# Natural *CMT2* variation is associated with genome-wide methylation changes and temperature seasonality

**DOI:** 10.1101/004119

**Authors:** Xia Shen, Jennifer De Jonge, Simon Forsberg, Mats Pettersson, Zheya Sheng, Lars Hennig, Örjan Carlborg

## Abstract

As *Arabidopsis thaliana* has colonized a wide range of habitats across the world it is an attractive model for studying the genetic mechanisms underlying environmental adaptation. Here, we used public data from two collections of *A. thaliana* accessions to associate genetic variability at individual loci with differences in climates at the sampling sites. We use a novel method to screen the genome for plastic alleles that tolerate a broader climate range than the major allele. This approach reduces confounding with population structure and increases power compared to standard genome-wide association methods. Sixteen novel loci were found, including an association between Chromomethylase 2 (*CMT2*) and temperature seasonality where the genome-wide CHH methylation was different for the group of accessions carrying the plastic allele. *Cmt2* mutants were shown to be more tolerant to heat-stress, suggesting genetic regulation of epigenetic modifications as a likely mechanism underlying natural adaptation to variable temperatures, potentially through differential allelic plasticity to temperature-stress.

**AUTHOR SUMMARY:** A central problem when studying adaptation to a new environment is the interplay between genetic variation and phenotypic plasticity. *Arabidopsis thaliana* has colonized a wide range of habitats across the world and it is therefore an attractive model for studying the genetic mechanisms underlying environmental adaptation. Here, we study two collections of *A. thaliana* accessions from across Eurasia to identify loci associated with differences in climates at the sampling sites. A new genome-wide association analysis method was developed to detect adaptive loci where the alleles tolerate different climate ranges. Sixteen novel such loci were found including a strong association between Chromomethylase 2 (*CMT2*) and temperature seasonality. The reference allele dominated in areas with less seasonal variability in temperature, and the alternative allele existed in both stable and variable regions. Our results thus link natural variation in *CMT2* and epigenetic changes to temperature adaptation. We showed experimentally that plants with a defective *CMT2* gene tolerate heat-stress better than plants with a functional gene. Together this strongly suggests a role for genetic regulation of epigenetic modifications in natural adaptation to temperature and illustrates the importance of re-analyses of existing data using new analytical methods to obtain deeper insights into the underlying biology from available data.

## INTRODUCTION

*Arabidopsis thaliana* has colonized a wide range of habitats across the world and it is therefore an attractive model for studying the genetic mechanisms underlying environmental adaptation [1]. Several large collections of *A. thaliana* accessions have either been whole-genome re-sequenced or high-density SNP genotyped [1–7]. The included accessions have adapted to a wide range of different climatic conditions and therefore loci involved in climate adaptation will display genotype by climate-at-sampling-site correlations in these populations. Genome-wide association or selective-sweep analyses can therefore potentially identify signals of natural selection involved in environmental adaptation, if those can be disentangled from the effects of other population genetic forces acting to change the allele frequencies. Selective-sweep studies are inherently sensitive to population-structure and, if present, the false-positive rates will be high as the available statistical methods are unable to handle this situation properly. Further experimental validation of inferred sweeps (e.g. [1,8]) is hence necessary to suggest them as adaptive. In GWAS, kinship correction is now a standard approach to account for population structure that properly controls the false discovery rate. Unfortunately, correcting for genomic kinship often decreases the power to detect individual adaptive loci, which is likely the reason that no genome-wide significant associations to climate conditions were found in earlier GWAS analyses [1,8]. Nevertheless, a number of candidate adaptive loci could despite this be identified using extensive experimental validation [1,2,8], showing how valuable these populations are as a resource for finding the genomic footprint of climate adaptation.

Genome-wide association (GWA) datasets based on natural collections of *A. thaliana* accessions, such as the RegMap collection, are often genetically stratified. This is primarily due to the close relationships between accessions sampled at nearby locations. Furthermore, as the climate measurements used as phenotypes for the accessions are values representative for the sampling locations of the individual accessions, these measurements will be confounded with the general genetic relationship [9]. Unless properly controlled for, this confounding might lead to excessive false-positive signals in the association analysis; this as the differences in allele-frequencies between loci in locations that differ in climate, and at the same time are geographically distant, will create an association between the genotype and the trait. However, this association could also be due to other forces than selection. In traditional GWA analyses, mixed-model based approaches are commonly used to control for population-stratification. The downside of this approach is that it, in practice, will remove many true genetic signals coming from local adaptation due to the inherent confounding between local genotype and adaptive phenotype. Instead, the primary signals from such analyses will be due to effects of alleles that exist in, and have similar effects across, the entire studied population. In general, studies into the contributions of genetic variance-heterogeneity to the phenotypic variability in complex traits is a novel and useful approach with great potential [10]. Here, we have developed and used a new approach that combines a linear mixed model and a variance-heterogeneity test, which addresses these initial concerns and shown that it is possible to infer statistically robust results of genetically regulated phenotypic variability in GWA data from natural populations.

This study describes the results from a reanalysis of data from the RegMap collection to find loci contributing to climate adaptation through an alternative mechanism:genetic control of plasticity. Such loci are unlikely to be detected with standard GWAS or selective-sweep analyses as they have a different genomic signature of selection and distribution across climate envelopes. The reason for this difference is that plastic alleles are less likely to be driven to fixation by directional selection, but rather that multiple alleles remain in the population under extended periods of time by balancing selection [11]. To facilitate the detection of such loci, we extend and utilize an approach [12,13] that instead of mapping loci by differences in allele-frequencies between local environments, which is highly confounded by population structure, infer adaptive loci using a heterogeneity-of-variance test. This identifies loci where the minor allele is associated with a broader range of climate conditions than the major allele [12]. As such widely distributed alleles will be present across the entire population, they are less confounded with population structure and detectable in our GWAS analysis that utilizes kinship correction to account for population stratification.

## RESULTS

### Genome-wide association analysis to detect loci with plastic response to climate

A genome-wide association analysis was performed for thirteen climate variables across ∼215,000 SNPs in 948 *A. thaliana* accessions from the RegMap collection, representing the native range of the species [1,9]. In total, sixteen genome-wide significant loci were associated with eight climate variables (Table 1), none of which could be found using standard methods for GWAS analyses [1,8,14–16]. The effects were in general quite large, from 0.3 to 0.5 residual standard deviations (Table 1), meaning that the minor allele is associated with a climate that is between 21–35% more variable than that of the major allele. The detailed results from the association analysis for each of these climate variables are reported in Text S1 (Supplementary Figure 1–13). As expected, there was low confounding between the alleles associated with a broader range of climate conditions and population structure. This is illustrated by the plots showing the distributions of these alleles across the population strata in relation to their geographic origin and the climate envelopes in Text S1 (Supplementary Figure 14–35).

**Figure 1.**
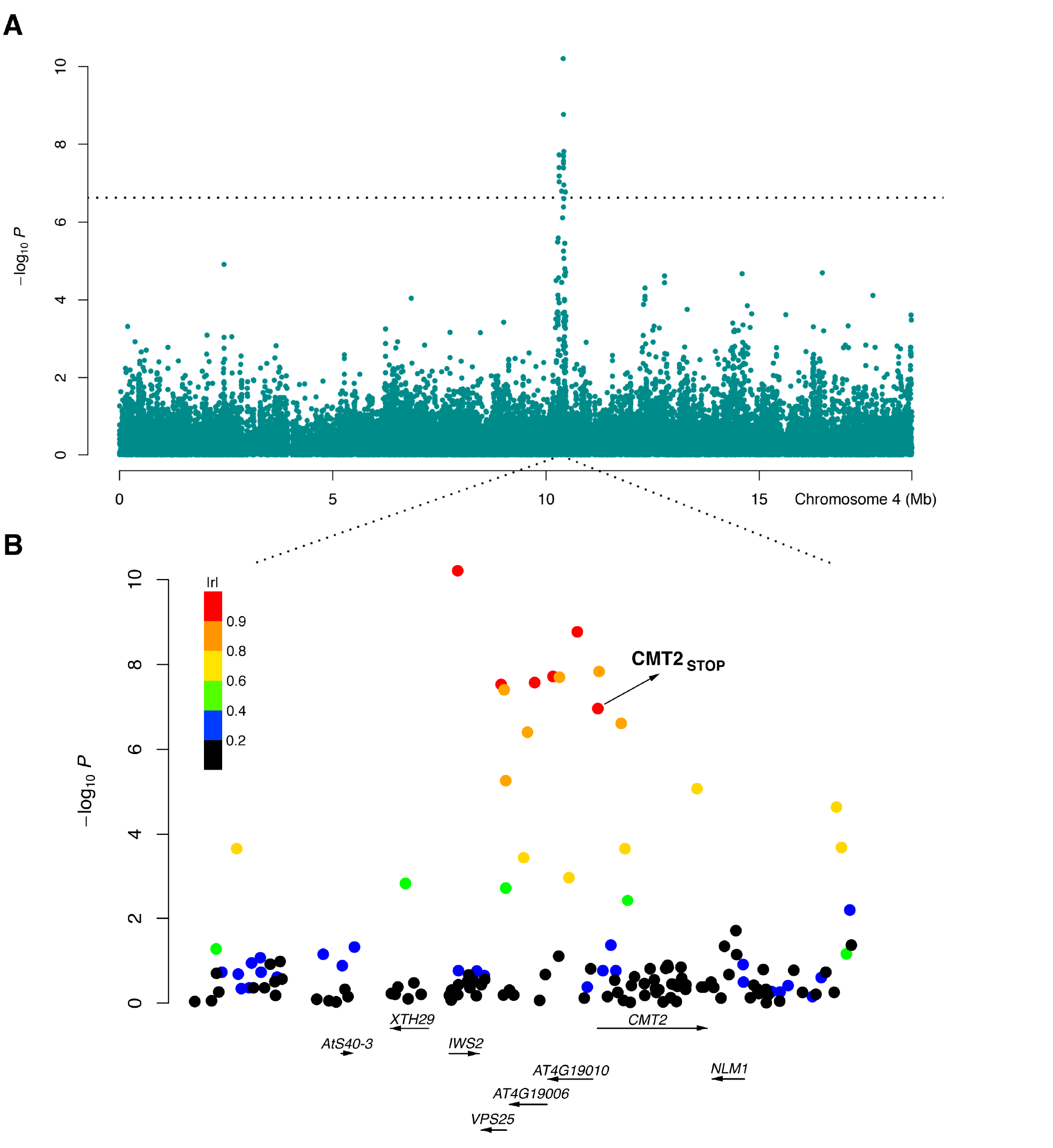
An LD block associated with temperature seasonality contains *CMT2*. A genome-wide significant variance-heterogeneity association signal was identified for temperature seasonality in the RegMap collection of natural *Arabidopsis thaliana* accessions [1]. The peak on chromosome 4 around 10 Mb (A) mapped to a haplotype block (B) containing a nonsense mutation *(CMT2_STOP_)* early in the first exon of the Chromomethylase 2 *(CMT2)* gene. Color coding based on |r| (the absolute value of the correlation coefficient) as a measure of LD between each SNP in the region and the leading SNP in the association analysis.

**Table I.** Loci with genome-significant, non-additive effects on climate adaptation and a functional analysis of nearby genes (*r*^2^ > 0.8) containing missense or nonsense mutations.

| Trait | Leading SNP |  |  |  |  | Selected Candidate Genes |  |  | Mutant Analysis<sup>1</sup> |  |
| --- | --- | --- | --- | --- | --- | --- | --- | --- | --- | --- |
|  | Chrom | Pos (bp) | MAF<sup>2</sup> | P-value<sup>3</sup> | Effect<sup>4</sup> | Gene name | Locus | # Mut<sup>5</sup> | PASE | MSA |
| Temperature seasonality |  |  |  |  |  |  |  |  |  |  |
|  | 2<sup>a</sup> | 12 169 701 | 0.08 | 2.0E-08 | 0.53±0.08 | *BGAL8* | *AT2G28470* | 4 | 0.36<sup>6</sup> | 0.72<sup>6</sup> |
|  | 4 | 10 406 018 | 0.05 | 6.1E-11 | 0.51±0.06 | *IWS2* | *AT4G19000* | 1 | 0.21 | 0.35 |
|  |  |  |  |  |  | *CMT2* | *AT4G19020* | 7 | STOP | STOP |
| Maximum temperature in the warmest month |  |  |  |  |  |  |  |  |  |  |
|  | 1 | 6 936 457 | 0.05 | 1.9E-07 | 0.34±0.07 |  | *AT1G19990* | 1 | 0.64 | 0.2 |
| Minimum temperature in the coldest month |  |  |  |  |  |  |  |  |  |  |
|  | 2<sup>b</sup> | 18 620 697 | 0.08 | 3.5E-08 | 0.33±0.05 |  |  |  |  |  |
|  | 2<sup>c</sup> | 19 397 389 | 0.05 | 5.0E-08 | 0.38±0.06 |  |  |  |  |  |
|  | 5 | 14 067 526 | 0.07 | 4.2E-08 | 0.33±0.05 |  | *AT5G35930* | 1 | 0.30 | 0.05 |
|  | 5<sup>d</sup> | 18 397 418 | 0.11 | 1.2E-07 | 0.28±0.05 |  |  |  |  |  |
| Number of consecutive cold days |  |  |  |  |  |  |  |  |  |  |
|  | 2<sup>b</sup> | 18 620 697 | 0.08 | 1.7E-07 | 0.33±0.05 |  |  |  |  |  |
|  | 2<sup>c</sup> | 19 397 389 | 0.05 | 7.2E-08 | 0.39±0.06 |  |  |  |  |  |
|  | 5 | 7 492 277 | 0.08 | 4.3E-09 | 0.38±0.06 |  | *AT5G22560* | 4 | 0.63 | 0.11 |
|  | 5<sup>d</sup> | 18 397 418 | 0.11 | 1.9E-07 | 0.29±0.05 |  |  |  |  |  |
| Day length in spring |  |  |  |  |  |  |  |  |  |  |
|  | 2<sup>a</sup> | 12 169 701 | 0.08 | 2.0E-07 | 0.46±0.08 | *BGAL8* | *AT2G28470* | 4 | 0.36 | 0.72 |
|  | 3 | 12 642 006 | 0.07 | 9.4E-08 | 0.29±0.05 |  |  |  |  |  |
|  | 4<sup>e</sup> | 14 788 320 | 0.08 | 2.2E-08 | 0.39±0.06 | *VEL1* | *AT4G30200* | 2 | 0.26 | 0.06 |
| Relative humidity in spring |  |  |  |  |  |  |  |  |  |  |
|  | 3 | 1 816 353 | 0.07 | 1.2E-08 | 0.39±0.06 |  |  |  |  |  |
|  | 4<sup>e</sup> | 14 834 441 | 0.06 | 6.4E-08 | 0.49±0.08 | *VEL1* | *AT4G30200* | 2 | 0.26 | 0.06 |
|  |  |  |  |  | 0.43±0.07 | *XTH19* | *AT4G30290* | 1 | 0.14 | 0.49 |
|  | 5 | 8 380 640 | 0.07 | 6.3E-08 |  |  |  |  |  |  |
| Length of the growing season |  |  |  |  |  |  |  |  |  |  |
|  | 3 | 576 148 | 0.08 | 1.4E-07 | 0.27±0.04 |  |  |  |  |  |
| Number of consecutive frost-free days |  |  |  |  |  |  |  |  |  |  |
|  | 1 | 953 031 | 0.24 | 8.0E-08 | 0.25±0.04 | *SOM* | *AT1G03790* | 3 | 0.25 | 0.55 |
|  | 1 | 6 463 065 | 0.08 | 1.9E-07 | 0.33±0.06 |  |  |  |  |  |
|  | 2 | 9 904 076 | 0.22 | 2.2E-08 | 0.22±0.04 |  |  |  |  |  |

<sup>a,b,c,d,e</sup>Loci affecting affect multiple traits; <sup>1</sup>The predicted functional effect score for the strongest mis-sense mutation in the gene based on amino-acid physiochemical properties (PASE) and evolutionary conservation (MSA) [39]; <sup>2</sup>MAF: Minor Allele Frequency; <sup>3</sup>P-value: significance after genomic-control from a linear regression analysis of squared z-scores accounting for population stratification. <sup>4</sup>Effect: Standardized genetic effect on adaptability (Chi-square distributed) of squared z-scores accounting for population stratification; <sup>5</sup>#Mut: number mis- and non-sense mutations in the gene in the 1001-genomes dataset [2]. <sup>6</sup>Locus contains two missense mutations with equally strong predicted effects.

### Identification of candidate mutations using re-sequencing data from the 1001-genomes project

Utilizing the publicly available whole-genome re-sequencing data from the 1001-genomes project [2–7] (http://1001genomes.org), we screened the loci with significant associations to the climate variables for candidate functional polymorphisms. Missense, nonsense or frameshift mutations in high linkage disequilibrium (LD; *r*^2^ > 0.8) with the leading SNPs were identified in five functional candidate genes associated with eight climate variables (for details on these see Table 1) and 11 less characterized genes (Table S1). 76 additional linked loci or genes without candidate mutations in their coding regions are provided in Table S2.

### Several loci are associated with multiple climate variables

Interestingly, three out of the eight loci with missense mutations affected more than one climate variable, even though these were only marginally correlated. One such potentially pleiotropic adaptive effect for day length and relative humidity in the spring was associated with a locus containing the genes *VEL1* and *XTH19* (Table 1). The major allele at this locus was predominant in short-day regions, whereas the alternative allele was more plastic in relation to day-length. *XTH19* has been implied as a regulator of shade avoidance [17], but information about its potential involvement in regulation of photoperiodic length is lacking. *VEL1*, is a Plant Homeo Domain (PHD) finger protein. PHD finger proteins are known to affect vernalization and flowering of *A. thaliana*, e.g. by silencing the key flowering locus *FLC* during vernalization, and is involved in photoperiod-mediated epigenetic regulation of MAF5 [18–20]. The finding that *VEL1* is associated with day length and relative humidity is thus consistent with the role of previous reports on PHD finger proteins. It also makes this protein an interesting target for future studies into the genetics underlying simultaneous adaptation to day-length and humidity.

Another potentially pleiotropic adaptive effect was identified for two more highly correlated traits, minimum temperature and number of consecutive cold days (Pearson’s *r*^2^ = 0.76). In total, 17 missense mutations were found at this locus. The top candidate gene containing a missense mutation is galactinol synthase 1 (*GolS1*). This gene has been reported to be involved in extreme temperature-induced synthesis [21,22], making it an interesting target for further studies regarding the genetics of temperature adaptation.

### Chromomethylase 2 (CMT2) *is associated with temperature seasonality in the regmap collection*

A strong association to temperature seasonality, i.e. the ratio between the standard deviation and the mean of temperature records over a year, was identified near Chromomethylase 2 (*CMT2*; Table 1; Figure 1). Stable areas are generally found near large bodies of water (e.g. London near the Atlantic 11 ± 5°C; mean ± SD) and variable areas inland (e.g. Novosibirsk in Siberia 1 ± 14°C). A premature *CMT2* stop codon located on chromosome 4 at 10,414,556 bp (the 31st base pair of the first exon) segregated in the RegMap collection, with minor allele frequency of 0.05. This *CMT2_STOP_* allele had a genome-wide significant association with temperature seasonality (*P* = 1.1 × 10^−7^) and was in strong LD (*r*^2^ = 0.82) with the leading SNP (Figure 1B). The geographic distributions of the wild-type (*CMT2_WT_*) and the alternative (*CMT2_STOP_*) alleles in the RegMap collection shows that the *CMT2_WT_* allele dominates in all major sub-populations sampled from areas with low or intermediate temperature seasonality. The plastic *CMT2_STOP_* allele is present, albeit at lower frequency, across all sub-populations in low- and intermediate temperature seasonality areas, and is more common in areas with high temperature seasonality (Figure 2A; Figure 3; Text S1, Supplementary Figure 36). Such global distribution across the major population strata indicates that the allele has been around in the Eurasian population sufficiently long to spread across most of the native range and that the allele is not deleterious but rather maintained through balancing selection [11], perhaps by mediating an improved tolerance to variable temperatures.

**Figure 2.**
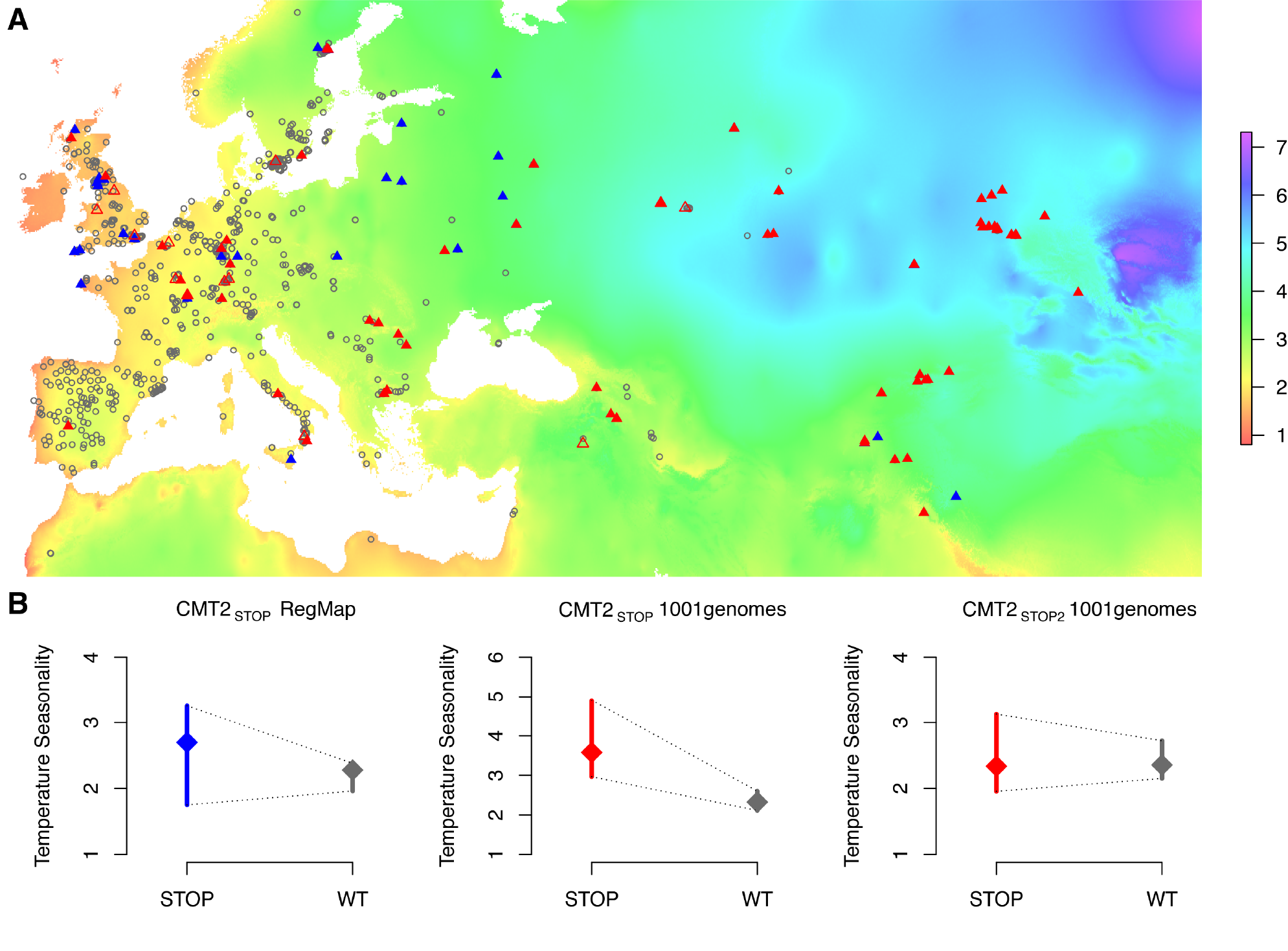
Geographic distribution of, and heterogenous variance for, three *CMT2* alleles in two collections of *A. thaliana* accessions. The geographic distributions (A) of the wild-type (*CMT2_WT_*; gray open circles) and two nonsense alleles *(CMT2_STOP_/CMT2_STOP_2*; filled/open triangles) in the *CMT2* gene that illustrates a clustering of *CMT2_WT_* alleles in less variable regions and a greater dispersion of the nonsense alleles across different climates both in the RegMap [1] (blue) and the 1001-genomes [2](red) *A. thaliana* collections. The resulting variance-heterogeneity in temperature seasonality between genotypes is highly significant, as illustrated by the quantile plots in (B) where the median is indicated by a diamond and a bar representing the 25% to 75% quantile range. The color scale indicate the level of temperature seasonality across the map. The colorkey in (A) represent the temperature seasonality values, given as the standard-deviation in % of the mean temperature (K).

**Figure 3.**
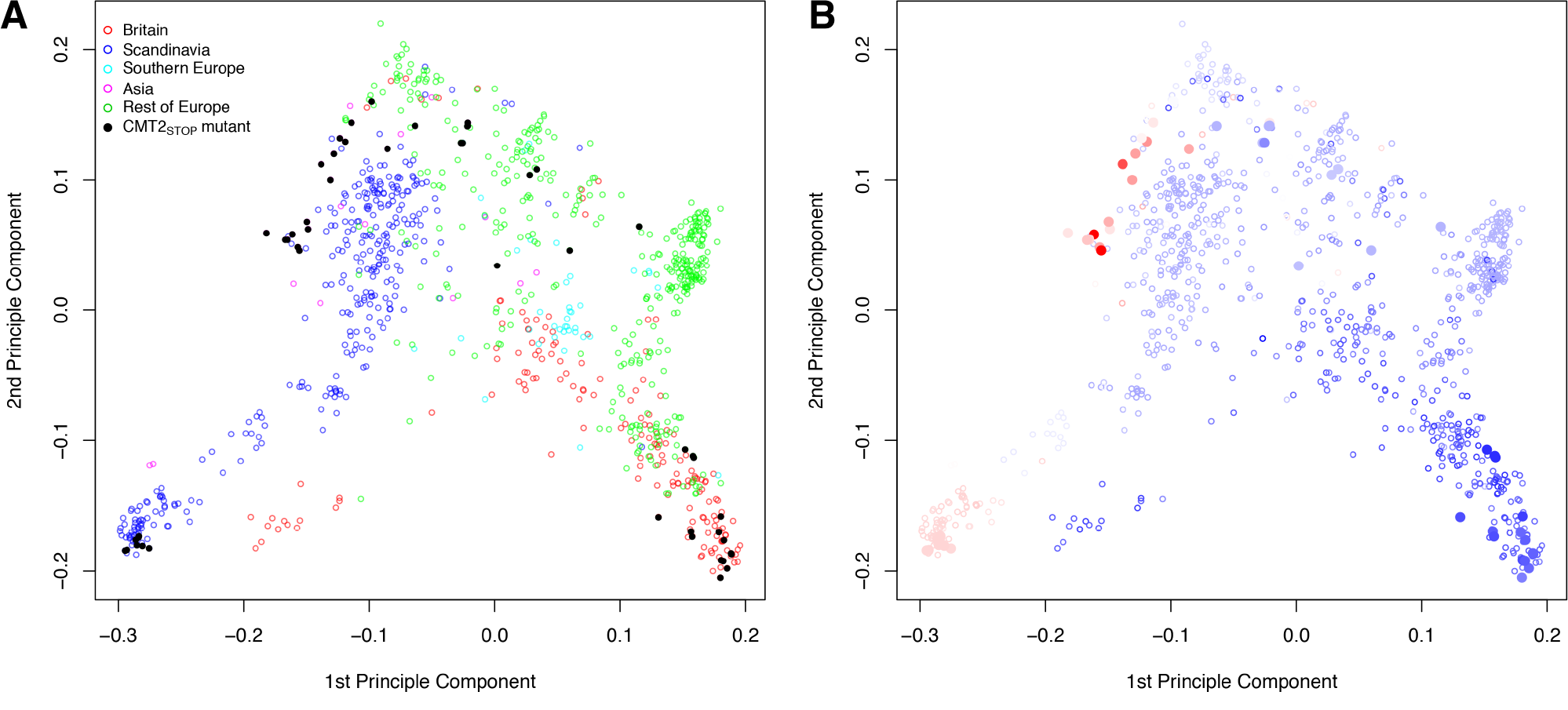
Principle components of the genomic kinship in the RegMap collection for the accessions carrying the alternative alleles at the Chromomethylase 2 locus *(CMT2_STOP_* and *CMT2_WT_* as filled and empty circles, respectively). Coloring is based on (A) geographical regions (defined as in Supplementary Fig. 37) and (B) temperature seasonality, ranging from dark blue (least variable) to red (most variable).

### Broader geographic distribution of the cmt2_stop_ allele in the 1001-genomes collection

To confirm that the *CMT2_STOP_* association was not due to sampling bias in the RegMap collection, we also scored the *CMT2* genotype and collected the geographical origins from 665 accessions that were part of the 1001-genomes project (http://1001genomes.org) [2,3,5–7]. In this more geographically diverse set (Figure 2A), *CMT2_STOP_* was more common (MAF = 0.10) and had a similar allele distribution across Eurasia as in RegMap (Text S1, Supplementary Figure 36–37). Two additional mutations were identified on unique haplotypes (*r*^2^ = 0.00)-one nonsense *CMT2_STOP2_* at 10,416,213 bp (MAF = 0.02) and a frameshift mutation at 10,414,640 bp (two accessions). Both *CMT2_STOP_* and *CMT2_STOP2_* had genotype-phenotype maps implying a plastic response to variable temperature (Figure 2B) and the existence of multiple mutations disrupting *CMT2* further suggest lack of *CMT2* function as a potentially evolutionary beneficial event [23].

### Accessions with the cmt2_stop_ allele has an altered genome-wide chh-methylation pattern

*CMT2* is a plant DNA methyltransferase that methylates mainly cytosines in CHH (H = any base but G) contexts, predominantly at transposable elements (TEs) [24,25]. We tested the effect of *CMT2_STOP_* on genome-wide DNA methylation using 135 *CMT2_WT_* and 16 *CMT2_STOP_* accessions, for which high-quality MethylC-sequencing data was publicly available [7]. In earlier studies [24,25], it has been shown that *CMT2*-mediated CHH methylation primarily affects TE-body methylation. In *CMT2* knock-outs in a Col-0 genetic background, this results in a near lack of CHH methylation at such sites. Here, we compared the levels of CHH-methylation across TE’s between *CMT2_STOP_* and *CMT2_WT_* accessions. Our analyses revealed that the accessions carrying the *CMT2_STOP_* allele on average had a small (1%) average decrease in CHH-methylation across the TE-body compared to the *CMT2_WT_* accessions. A more detailed analysis showed that this difference was primarily due to two of 16 *CMT2_STOP_* accessions, Kz-9 and Neo-6, showing a TE-body CHH methylation pattern resembling that of the *cmt2* knockouts in the data of [24]. Interestingly, none of the 135 *CMT2_WT_* accessions displayed such a decrease in TE-body CHH methylation, and hence there is a significant increase in the frequency of the *cmt2* knock-out TE-body CHH methylation pattern among the natural *CMT2_STOP_* accessions (*P* = 0.01; Fisher’s exact test). Our analyses show that the methylation-pattern is more heterogeneous among the natural accessions than within the Col-0 accession, both for the *CMT2_STOP_* and *CMT2_WT_* accessions (both P = 0.01; Brown-Forsythe heterogeneity of variance test; Figure 4). There is thus a significant association between the *CMT2_STOP_* polymorphism and decreased genome-wide TE-body CHH-methylation levels, and we show that this is apparently due to an increased frequency of the *cmt*2-mutant methylation phenotype. Further, the results also show a variable contribution of *CMT2*-independent CHH methylation pathways in the natural accessions. The reason why not all *CMT2_STOP_* accessions behave like null alleles is unclear, but the variability amongst in the level of CHH-methylation across the natural accessions suggest that it is possible that *CMT2*-independent pathways, such as the RNA-dependent DNA-methylation pathway, compensate for the lack of *CMT2* due to segregating polymorphisms also at these loci. Alternatively, *CMT2_STOP_* alleles may not be null, maybe due to stop codon read-through, which is more common than previously thought [26]. Although our analyses of genomewide methylation data have established that *CMT2_STOP_* allele has a quantitative effect on CHH methylation, further studies are needed to fully explore the link between the *CMT2_STOP_* allele, other pathways affecting genome-wide DNA-methylation and their joint contributions to the inferred association to temperature seasonality.

**Figure 4.**
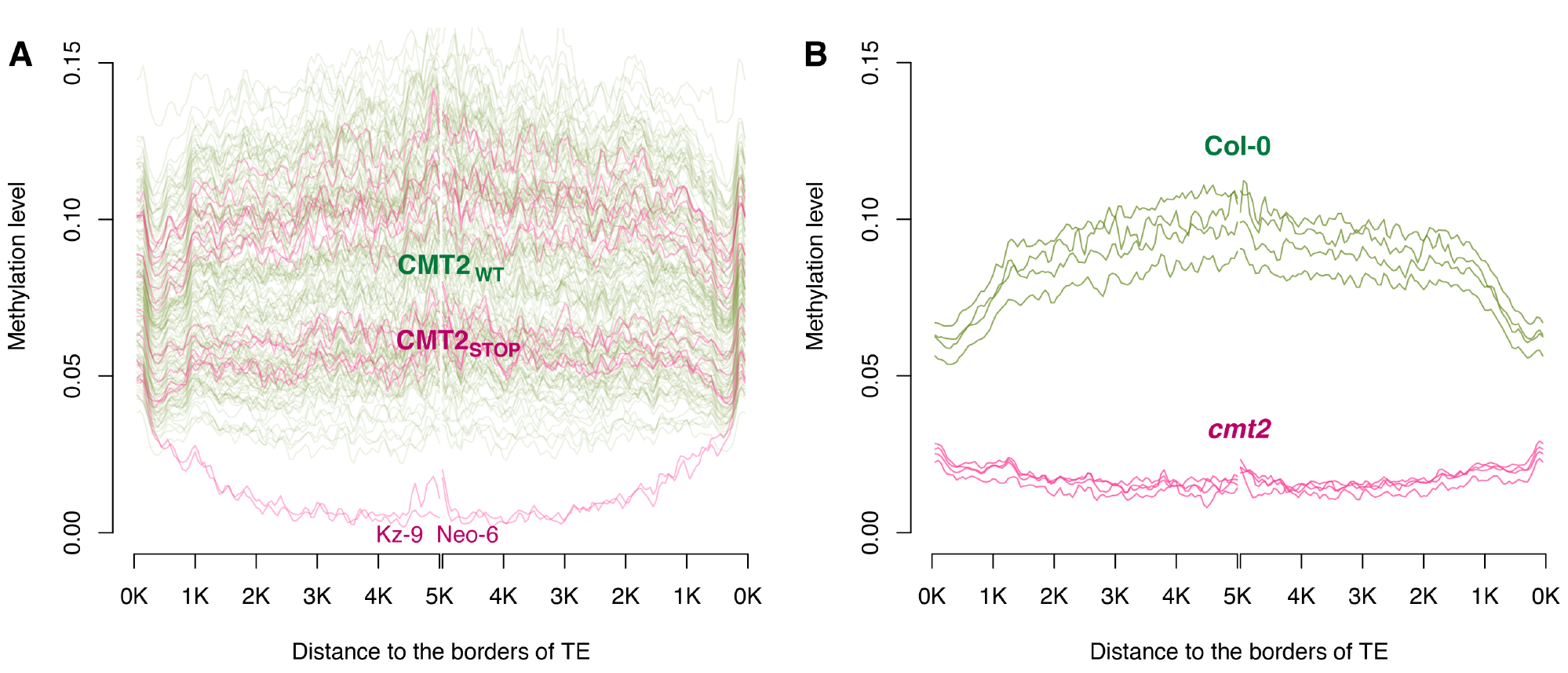
Comparison of CHH methylation patterns inside TE-bodies, (A) between *CMT2_WT_* and *CMT2_STOP_* accessions using the data from [7], and (B) between four replicate Col-0 wild-type and *CMT2* knock-outs from [24]. For each accession, the curve is to illustrate the moving average methylation level in a sliding 100 bp window. On the x-axis, the two different strands of DNA are aligned in the middle, truncated at 5 kb from the edge of the TEs.

### Cmt2 *mutant plants have an improved heat-stress tolerance*

To functionally explore whether *CMT2* is a likely contributor to the temperature-stress response, we have subjected *CMT2* mutants to two types of heat-stress. First, we tested the reaction of Col-0 and the *cmt2-5* null mutant (Supplementary Fig. 45) to severe heat-stress (24h at 37°C). This treatment was used because it can release transcriptional silencing of some TEs [27] and could thus be a good starting point to evaluate potential stress effects on *cmt2*. Under these conditions, the *cmt2* mutant had significantly higher survival-rate (1.6-fold; *P* = 9.1 × 10^−3^; Figure 5A) than Col-0. To evaluate whether a similar response could also be observed under less severe, non-lethal stress, we subjected the same genotypes to heat-stress of shorter duration (6h at 37°C) and measured root growth after stress as a measure of the ability of plants to recover. Also under these conditions, the *cmt2* mutant was found to be more tolerant to heat-stress, as its growth was less affected after being stressed (Figure 5B; 1.9-fold higher in *cmt2; P* =0.026, one-sided t-test). This striking improvement in tolerance to heat-stress of *cmt2* plants suggests *CMT2*-dependent CHH methylation as an important alleviator of stress responses in *A. thaliana* and a candidate mechanism for temperature adaptation.

**Figure 5.**
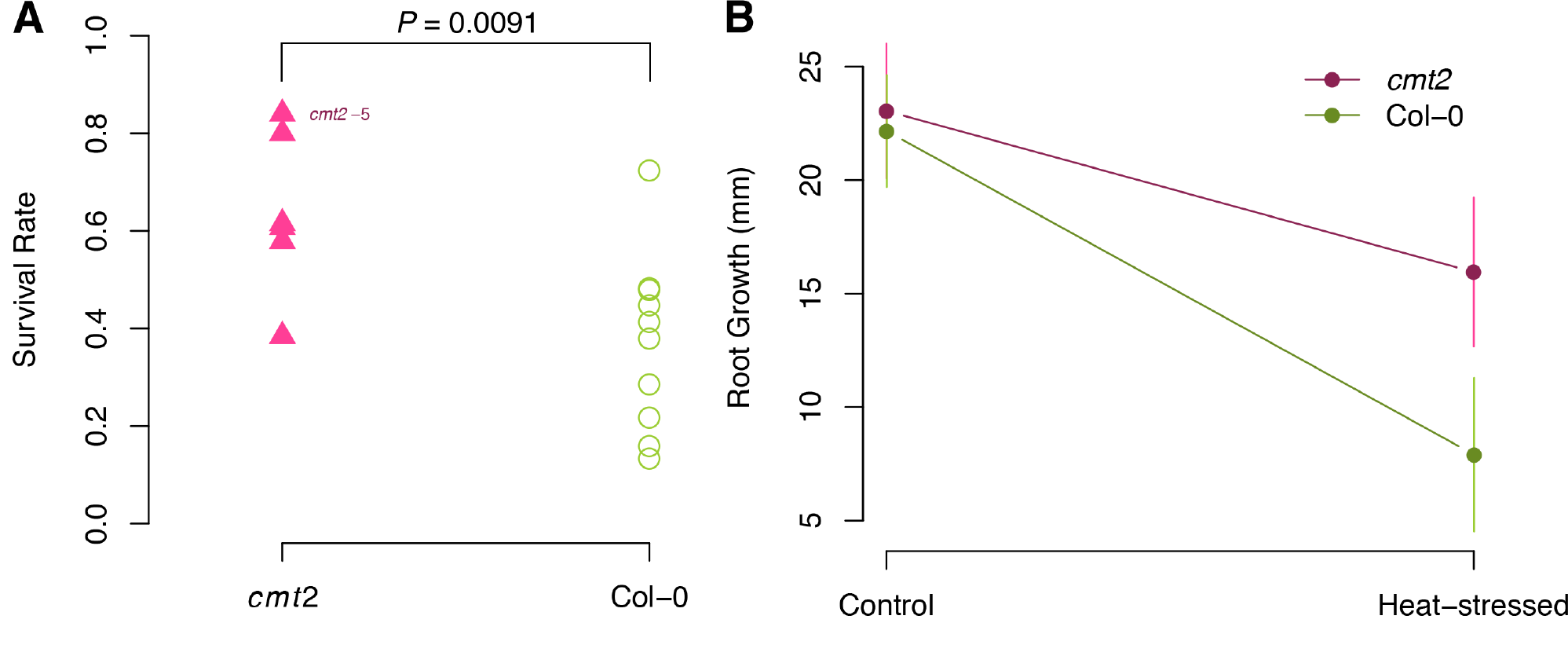
*cmt2* mutant plants display an increased tolerance to heat-stress. A. The survival rate is significantly higher for *cmt2-5* mutant than for Col-0 plants under severe heat-stress (24 h at 37.5°C). *P*-values in A were obtained using a log-linear regression. B. The *CMT2-5* mutant was also more tolerant to less severe heat-stress heat-stress (6 h at 37.5°C) than Col-0, here illustrated by its significantly faster growth of the root (*P* = 0.026; one-sided t-test) during the first 48h following heat stress.

### The CMT2_STOP_ allele is associated with increased leaf serration and higher disease presence after bacterial inoculation

To also explore the potential effects of the *CMT2_STOP_* allele on other phenotypes measured in collections of natural accessions, we tested for associations between this *CMT2* polymorphism and the 107 phenotypes measured as part of a previous study [28]. Three phenotypes were found to be significantly associated with the genotype at this locus (Supplementary Figure 39).

Associations were found to two phenotypes related to disease presence following inoculation with *Pseudomonas viridiflava* (strains PNA3.3a and ME3.1b; *P* = 4.8 × 10^−3^ and *P* = 1.3 × 10^−4^, respectively). Scoring of disease was done by eye four days after inoculation in 6 replicates / strain × accession using a scale from 0 (no visible symptom) to 10 (leaves collapse and turn yellow) with an increment of 1[28]. The connection between an increased susceptibility (0.6 and 0.7 units for PNA3.3a and ME3.1b, respectively) to disease and an increased tolerance to temperature seasonality is not obvious. However, recent work by [29] has shown that widespread dynamic CHH-methylation is important for the response to *Pseudomonas syringae* infection. In light of this finding, it is therefore not unlikely that these phenotypes are functionally related via an altered CMT2-mediated CHH-methylation in response to abiotic and biotic stress.

An association was also found for the level of leaf serration (increase by 0.23 units for the *CMT2_STOP_* allele; *P* = 3.3 × 10^−3^), determined after growth for 8 weeks at 10°C (level from 0: entire lamina, to 1.5: sharp/ jagged serration), across 4 plants/accession[28]. Measures of leaf serration were also available at 16 and 22°C, and interestingly there was a significant *CMT2* genotype × temperature interaction (*P* = 0.048). The *CMT2_STOP_* accessions have the same level of serration across the three measured temperatures, whereas the level of serration decreases with temperature for the *CMT2_WT_* accessions (Text S1, Supplementary Fig. 38). Although we are not aware of any earlier results connecting leaf serration to the *CMT2* locus or the level of CHH-methylation in the plant, this result further indicate that the effects of the *CMT2_STOP_* and the *CMT2_WT_* alleles depends on temperature.

## DISCUSSION

A major challenge in attempts to identify individual loci involved in climate adaptation is the strong confounding between geographic location, climate and population structure in the natural *A. thaliana* population. Earlier genome-wide association analyses in large collections of natural accessions experienced a lack of statistical power when correcting for population-structure [1,8]. We used an alternative GWAS approach [12] to test for a variance-heterogeneity, instead of a mean difference, between genotypes. This analysis identifies loci where the minor allele is more plastic (i.e. exist across a broader climatic range) than the major allele. As it has low power to detect cases where the minor allele is associated with a lower variance (here with local environments), it will not map private alleles in local environments in a genome-wide analysis[12,30]. In contrast, a standard GWAS map loci where the allele-frequencies follow the climatic cline. Although plastic alleles might be less frequent in the genome, they are easier to detect in this data due to their lower confounding with population-structure. This overall increase in power is also apparent when comparing the signals that reach a lower, sub-GWAS significance level (Text S1, Supplementary Figure 40–44).

Several novel genome-wide significant associations were found to the tested climate variables, and a locus containing *VEL1* was associated to both day length and relative humidity in the spring. *A thaliana* is a facultative photoperiodic flowering plant and hence non-inductive photoperiods will delay, but not abolish, flowering. A genetic control of this phenotypic plasticity is thus potentially an adaptive mechanism. *VEL1* regulates the epigenetic silencing of genes in the FLC-pathway in response to vernalization [19] and photoperiod length [20] resulting in an acceleration of flowering under non-inductive photoperiods. Our results suggest that genetically plastic regulation of flowering, via the high-variance *VEL1* allele, might be beneficial under short-day conditions where both accelerated and delayed flowering is allowed. In long-daytime areas, accelerated flowering is potentially detrimental hence the wild-type allele has the highest adaptive value. It can be speculated whether this is connected to the fact that day-length follows a latitudinal cline, where early flowering might be detrimental in northern areas where accelerated flowering, when the day-length is short, could lead to excessive exposure to cold temperatures in the early spring and hence a lower fitness.

A particularly interesting finding in our vGWAS was the strong association between the *CMT2*-locus and temperature seasonality. Here the allele associated with higher temperature seasonality (i.e the plastic allele) had an altered genome-wide CHH methylation pattern where some accessions displayed a TE-body CHH methylation pattern similar to that of *cmt2* mutant plants. Interestingly, we also found that *cmt2* mutants were more tolerant to both mild and severe heat-stress, a finding that both strongly implicates *CMT2* as an adaptive locus and clearly illustrates the potential of our method as a useful approach to identify novel associations of functional importance.

It is not clear via which mechanism *CMT2*-dependent CHH methylation might affect plant heat tolerance. Although our results show that the *CMT2_STOP_* allele is present across regions with both low and high temperature seasonality, it remains to be shown whether this is due to this allele being generally more adaptable across all environments, or whether the *CMT2_WT_* allele is beneficial in environments with stable temperature and the *CMT2_STOP_* in high temperature seasonality areas. Regardless, we consider it most likely that the effect will be mediated by TEs in the immediate neighborhood of protein-coding genes. Heterochromatic states at TEs can affect activity of nearby genes and thus potentially plant fitness [31]. Consistent with a repressive role of *CMT2* on heat stress responses, *CMT2* expression is reduced by several abiotic stresses including heat [32]. Because global depletion of methylation has been shown to enhance resistance to biotic stress [29], it is possible that DNA-methylation has a broader function in shaping stress responses than currently thought.

Our results show that *CMT2_STOP_* accessions have more heterogeneous CHH methylation patterns than *CMT2_WT_* accessions. The *CMT2_STOP_* polymorphism is predicted to lead to a non-functional *CMT2* protein, and hence a genome-wide CHH-methylation profile resembling that of a complete *CMT2* mutant [24]. Although some of the accessions carrying the *CMT2_STOP_* allele displayed this pattern with a lower CHH-methylation inside TE-bodies, most of these accessions did not have any major loss of genome-wide CHH methylation. Such heterogeneity might indicate the presence of compensatory mechanisms and hence that the effects of altered *CMT2* function could be dependent on the genetic-background. This is an interesting finding that deserves further investigation, although such work is beyond the scope of the current study. Our interpretation of the available results is that our findings reflect the genetic heterogeneity among the natural accessions studied. In light of the recent report by [25], who showed a role also of *CMT3* in TE-body CHH methylation, it is not unlikely that the regulation of CHH methylation may result from the action and interaction of several genes.

We identified several alleles associated with a broader range of climates across the native range of *A. thaliana*, suggesting that a genetically mediated plastic response might of important for climate adaptation. Using publicly available data from several earlier studies, we were able to show that an allele at the *CMT2* locus displays an altered genome-wide CHH-methylation pattern was strongly associated with temperature seasonality. Using additional experiments, we also found that *CMT2* mutant plants tolerated heat-stress better than wild-type plants. Together, these findings suggest this genetically determined epigenetic variability as a likely mechanism contributing to a plastic response to the environment that has been of adaptive advantage in natural environments.

## MATERIALS AND METHODS

### Climate data andgenotyped arabidopsis thaliana accessions

Climate phenotypes and genotype data for a subset of the *A. thaliana* RegMap collection were previously analyzed by [1]. We downloaded data on 13 climate variables and genotypes of 214,553 single nucleotide polymorphisms (SNPs) for 948 accessions from: http://bergelson.uchicago.edu/regmap-data/climate-genome-scan. The climate variables used in the analyses were: aridity, number of consecutive cold days (below 4 degrees Celsius), number of consecutive frost-free days, day-length in the spring, growing-season length, maximum temperature in the warmest month, minimum temperature in the coldest month, temperature-seasonality, photosynthetically active radiation, precipitation in the wettest month, precipitation in the driest month, precipitation-seasonality, and relative humidity in the spring. More information on these variables is provided by [1]. No squared pairwise Pearson’s correlation coefficients between the phenotypes were greater than 0.8 (Figure S7 of [1]).

We calculated the temperature seasonality for at sampling locations of a selection of 1001-genomes (http://1001genomes.org) accessions. Raw climate data was downloaded from http://www.worldclim.org/, re-formatted and thereafter processed by the raster package in R. The R code for generating this data is provided in Text S1. The genotype for the *CMT2_STOP_* polymorphism was obtained by extracting the corresponding SNP data for the 1001-genomes accessions.

### Statistical modeling in genome-wide scans for adaptability

The climate data at the geographical origins of the *A. thaliana* accessions were treated as phenotypic responses. Each climate phenotype vector y for all the accessions were normalized via an inverse-Gaussian transformation. The squared normalized measurement 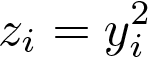of accession *i* is modeled by the following linear mixed model to test for an association with climate adaptability (i.e. a greater plasticity to the range of the environmental condition):

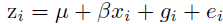

where *μ* is an intercept, *x_i_* the SNP genotype for accession *i* β the genetic SNP effect, 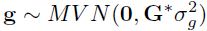 the polygenic effects and 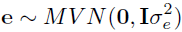 the residuals. *x_i_* is coded 0 and 2 for the two homozygotes (inbred lines). The genomic kinship matrix G* is constructed via the whole-genome generalized ridge regression method HEM (heteroscedastic effects model) [13] as **G\*** = **ZWZ′**, where **Z** is a number of individuals by number of SNPs matrix of genotypes standardized by the allele frequencies. **W** is a diagonal matrix with element 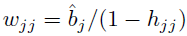 for the *j* -th SNP, where 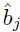 is the SNP-BLUP (SNP Best Linear Unbiased Prediction) effect estimate for the *j* -th SNP from a whole-genome ridge regression, and *h_jj_* is the hat-value for the *j* –th SNP. Quantities in **W** can be directly calculated using the **bigRR** package [13] in R. An example R source code for performing the analysis is provided in the Text S1.

The advantage of using the HEM genomic kinship matrix **G\***, rather than an ordinary genomic kinship matrix **G** = **ZZ′**, is that HEM is a significant improvement of the ridge regression (SNP-BLUP) in terms of the estimation of genetic effects [13,33]. Due to this, the updated genomic kinship matrix **G**\* better represents the relatedness between accessions and also accounts for the genetic effects of the SNPs on the phenotype.

### Testing and quality control for association with climate adaptability

The test statistic for the SNP effect β is constructed as the score statistic [34]:

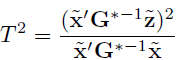
implemented in the GenABEL package [35],where 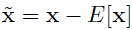 are the centered genotypic values and 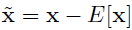 the centered phenotypic measurements. The *T*^2^ statistic has an asymptotic χ^2^ distribution with 1 degree of freedom. Subsequent genomic control (GC) [36] of the genome-wide association results was performed under the null hypothesis that no SNP has an effect on the climate phenotype. SNPs with minor allele frequency (MAF) less than 0.05 were excluded from the analysis. A 5% Bonferroni-corrected significance threshold was applied. As suggested by [30], the significant SNPs were also analyzed using a Gamma generalized linear model to exclude positive findings that might be due to low allele frequencies of the high-variance SNP.

### Statistical testing for associations between the CMT2_STOP_ polymorphism and phenotypes measured in a collection of natural accessions

The *CMT2_STOP_* genotype was extracted from the publicly available genome-wide genotype data with 107 phenotype measured from [28]. The association between the *CMT2_STOP_* genotype and each phenotype was tested by fitting a normal linear mixed model to account for population stratification, where the genomic kinship matrix was calculated by the ibs(, weight = ‘freq’) procedure in the GenABEL package [35], and the linear mixed model was fitted using the hglm package [37].

### Functional analysis of polymorphisms in loci with significant genome-wide associations to climate

All the loci that showed genome-wide significance in the association study was further characterized using the genome sequences of 728 accessions sequenced as part of the 1001-genomes project (http://1001genomes.org). Mutations within a ± 100Kb interval of each leading SNP and that are in LD with the leading SNP (*r*^2^ > 0.8) were reported (Table S1). The consequences of the identified polymorphisms were predicted using the Ensembl variant effect predictor [38] and their putative effects on the resulting protein estimated using the PASE (Prediction of Amino acid Substitution Effects) tool [39].

### Evaluation of TE-body methylation of CMT2_STOP_ and cmt2wt natural accessions

In a previous study, the methylation levels were scored at 43,182,344 sites across the genome using MethylC-sequencing in 152 natural *A. thaliana* accessions (data available at http://www.ncbi.nlm.nih.gov/geo/query/acc.cgi?acc=GSE43857) [7]. 135 of these accessions carried the *CMT2_WT_* and 17 the *CMT2_STOP_* alleles. Upon further inspection, the accession Rd-0 was excluded as it did not have sufficient sequence coverage to be used in the analyses. For each accession, across all TEs, moving averages of the CHH methylation level were calculated using a 100 bp sliding window from the borders of the TEs. The same analysis was also performed for four wild-type and four CMT2 knock-out accessions (data available at http://www.ncbi.nlm.nih.gov/geo/query/acc.cgi?acc=GSE41302) [24]. The results showing the TE-body CHH methylation patterns are visualized in Figure 4.

### Heat-stress treatments on cmt2 knockouts and natural CMT2_STOP_ accessions

A *CMT2* T-DNA insertion line (SAIL_906_G03, *cmt2–5* [24,40]) was ordered from NASC. Seeds of Col-0 wild-type and *cmt2-5* was then used for heat stress experiments based on a previously described protocol [27]. This treatment was used because it was shown to interfere with epigenetic gene silencing as evident from transcription of some TE [27]. Seeds were plated on 1/2 MS medium (0.8% agar, 1% sucrose), stratified for two days at 4°C in the dark and transferred to a growth chamber with 16 h light (110 μm^−2^ s^−1^, 22°C) and 8 h dark (20°C) periods. Ten-day-old seedlings were transferred to 4°C for one hour and subsequently placed for 6 h or 24 h at 37.5°C in the dark. Plant survival was scored two days after 24 h of heat stress with complete bleaching of shoot apices as lethality criterion (Text S1, Supplementary Fig. 46, Table S3). Experiments were repeated six times, each with ∼30 plants per genotype. Root length was measured immediately before the 6 h heat stress and two days after heat stress (Table S4).

A log-linear regression was conducted to test for the difference in survival rate between Col-0 and *cmt2-5* knockout, i.e. 
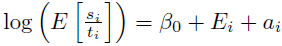

where *S_i_* is the number of surviving plants of accession *i, t_i_* the corresponding total number of plants, *E_i_* the experiment effect, *a_i_* the accession effect, and *β_0_* an intercept. The model fitting procedure was implemented using the `glm()` procedure in R, with option `family = gaussian(link = log)`, *s_i_* as response, *t_i_* as offset, and, *β_0_, E_i_, a_i_* as fixed effects.

## ACKNOWLEDGEMENTS

We thank Leif Andersson, Jennifer Lachowiec and Yanjun Zan for helpful input. Also, providers of pre-publication sequence data within the 1001-genomes project are acknowledged for their efforts in creating this community resource, including Monsanto Company, the Weigel laboratory at the Max Planck Institute for Developmental Biology, the IGS of the Center for Biotechnology of the University of Bielefeld, the DOE Joint Genome Institute (JGI), the Joint BioEnergy Institute, the Nordborg laboratory of the Gregor Mendel Institute of Molecular Plant Biology, the Bergelson lab of the University of Chicago and the Ecker lab of the Salk Institute for Biological Studies, La Jolla, CA.

## Supplementary Information for

### This PDF File Includes

Supplementary Text

Supplementary Fig. 1–46

Supplementary Table S1–S4

## Additional Results

### *CMT2* is a Potential Target for the Nonsense-Mediated RNA Decay (NMD) Pathway

The *CMT2_STOP_* allele is predicted to produce an mRNA with a pre-mature translation termination codon, which makes it a likely target for the nonsense-mediated mRNA decay (NMD) pathway (Nicholson and Muhlemann 2010). Thus, accessions carrying *CMT2_STOP_* may have lower transcript abundance than those with wild-type *CMT2*. To test whether this was the case, we evaluated the levels of *CMT2* mRNA using two publicly available datasets. First, we analyzed the 19 genomes project data (Gan et al. 2011) that contained full genome sequences and transcriptome for 19 *A. thaliana* accessions, two of which (Ct-1 and Kn-0) are part of the RegMap panel and carry the *CMT2_STOP_* mutation. The difference in mRNA abundance was highly significant between *CMT2* mutant and wild-type accessions (*P* = 6.9 × 10^−5^), with a higher expression in the wild type. A similar analysis was done using a larger data set from^7^, with CMT2 genotypes called from whole-genome re-sequencing data and using RNA-seq data from leaf tissue in 14 *CMT2_STOP_* and 92 *CMT2_WT_* accessions. As in the other dataset, the average mRNA abundance was higher for the CMT2 wild-type accessions, but here the difference was not significant (*P* = 0.14). However, the mRNA levels were significantly higher among the four *CMT2_STOP_* accessions that displayed a methylation pattern resembling that of the *CMT2_WT_* in the analysis above (t-test; *P* = 0.01). When those lines were omitted, the average mRNA level was significantly higher in the *CMT2_WT_* accessions than in the ten remaining *CMT2_STOP_* accessions (t-test; *P* = 0.02). Together, these results indicate that *CMT2* mRNA levels are often lower in *CMT2_STOP_* than in *CMT2* wild-type accessions. This is consistent with the notion that the premature stop codon does interfere with translation as expected.

## Methods

### Plant Material

A T-DNA insertion line (SAIL_906_G03, *cmt2-5*) (Figure S45) was ordered from NASC and seeds were produced from confirmed homozygous insertion plants. *CMT2* (AT4G19020) transcripts spanning the insertion site were assayed by RT-PCR with the following primers: forward_CMT2 CTCATTACCCCCAGAGGAGAG, reverse_CMT2 AAAGTTTGGGTCGAGGAAGAG. As a control a *PP2A* gene (AT1G13320) was used with primers forward_PP2A ATTCCGATAGTCGACCAAGC, reverse_PP2A: AACATCAACATCTGGGTCTTCA. RNA was extracted from five seedlings using the Trizol method as described previously (Steinbach and Hennig 2014). RNA was DNA treated with DNase I (Thermo Scientific) according to the manufacturer’s instructions. 1 *μ*g of RNA was used to synthesize cDNA with the RervertAid First Strand cDNA synthesis Kit (Thermo scientific) according to manufacturer’s instructions. No CMT2 transcripts spanning the insertion site were detected strongly suggesting that *cmt2-5* is a null allele.

## Source Code

### Example R source code for calculating hem genomic kinship matrix

Here, we use the example data in the **bigRR** package: http://cran.r-project.org/web/packages/bigRR/ to illustrate how an ordinary identity-by-state (IBS) kinship matrix can be update to a HEM genomic kinship matrix. The full theoretical details on this procedure are provided in^13^.

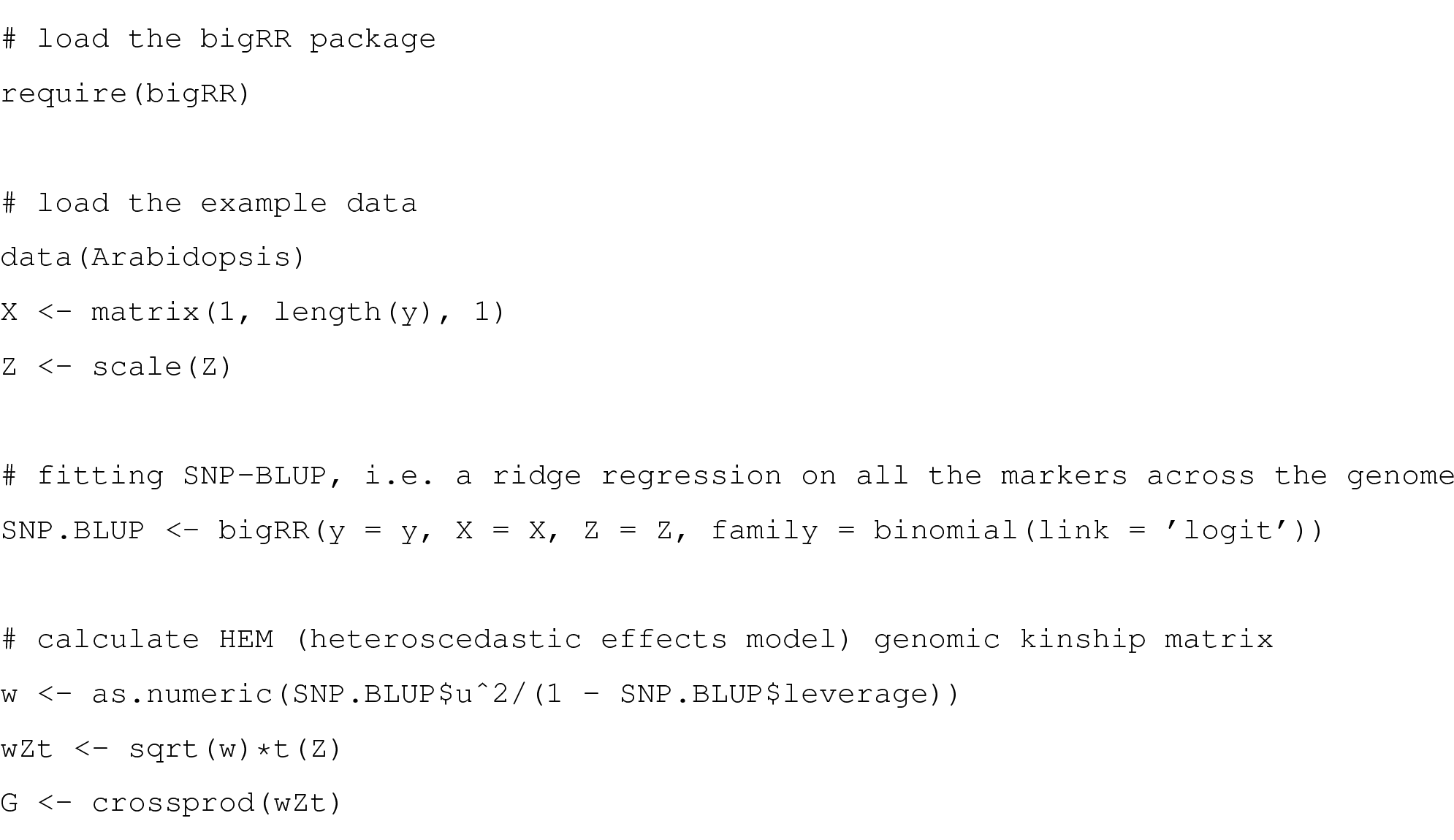

## Temperature seasonality phenotype preparation in R

We present the source code for phenotyping of temperature seasonality in Euro-Asia. The data downloaded from http://www.worldclim.org/ were processed using the following code to obtain an object readable by the **raster** package: http://cran.r-project.org/web/packages/raster/.

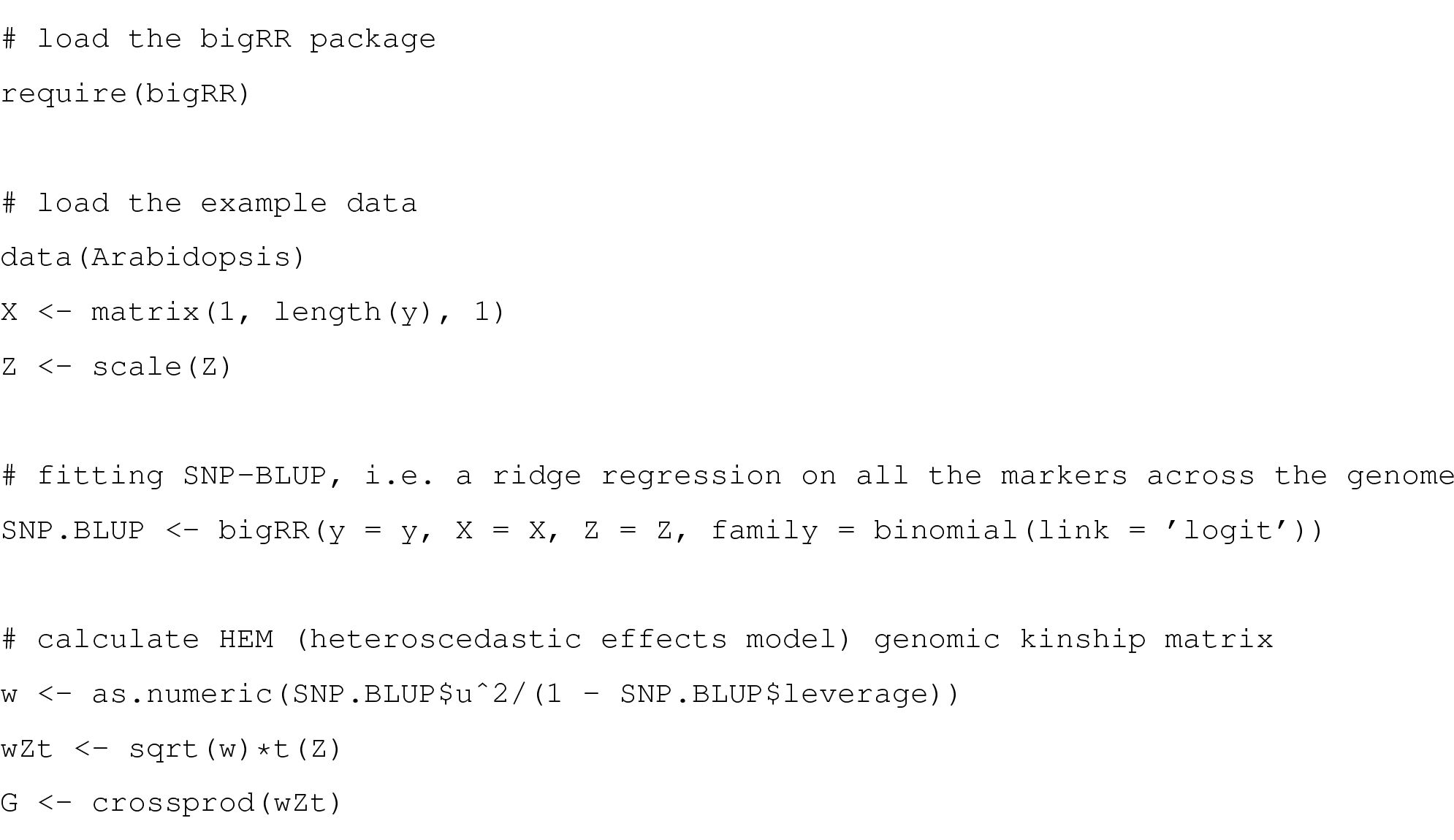

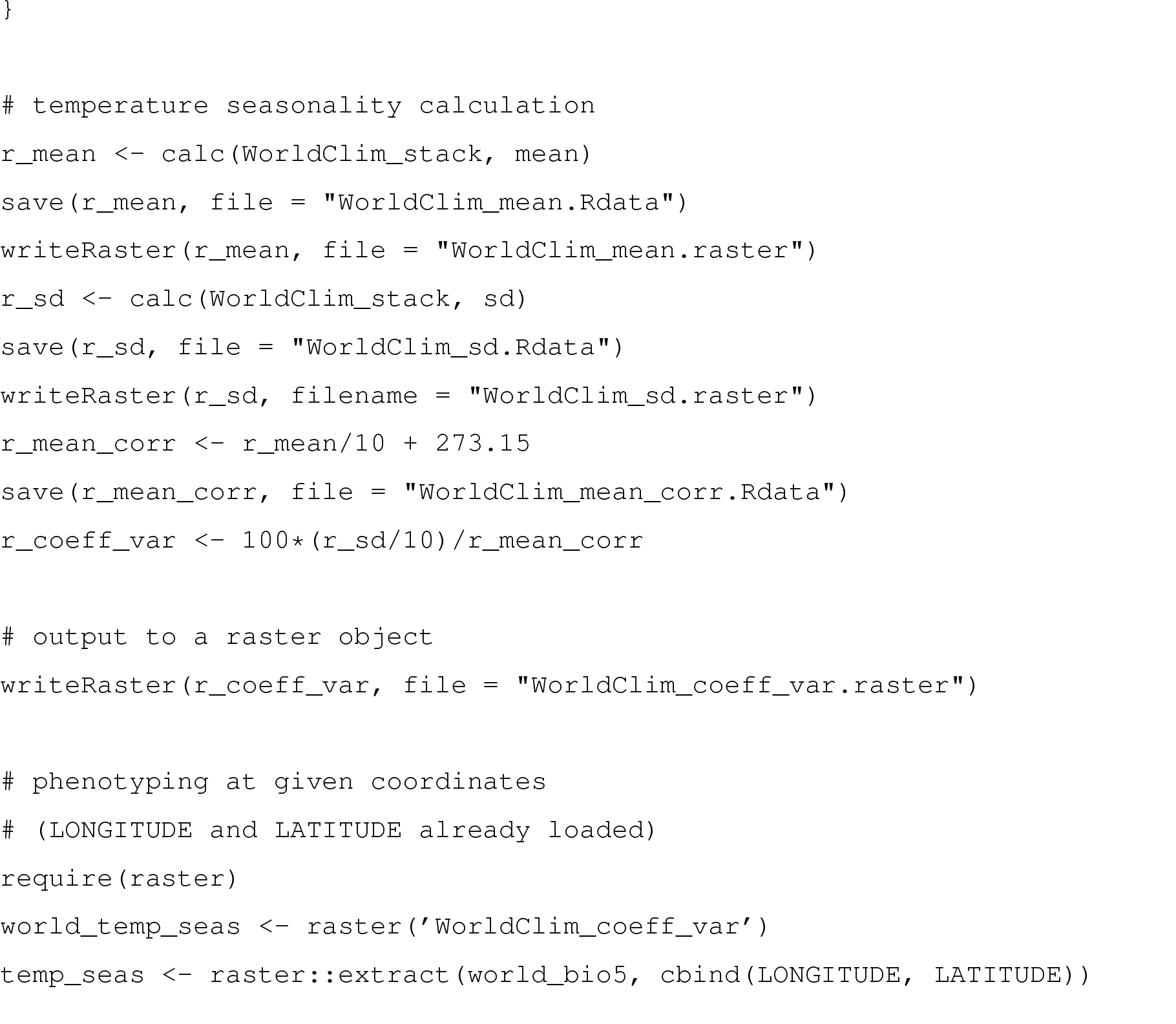

## Comments

### Statistical properties of the vGWAS in relation to population stratification

#### Inherent properties of the variance heterogeneity test decreases risk of identifying locally adapted alleles

An important property of the variance heterogeneity GWAS analysis is that it is inherently more powerful in detecting loci where the minor allele is associated with a higher variance than the major allele. In practice, there is no power to detect low-variance minor alleles in a GWAS setting^12,30^. Hence, the method facilitates detection of alternative *(i.e.* minor) alleles associated with a broader range of the climate variables than the reference (i.e. major) alleles. The method is powerful in finding associations to minor alleles associated with a broader range of climate-variables than the reference. Such alleles will, by definition, be present across a large proportion of the global population and due to this be considerably less affected by population structure. In Figure 3 or Figure S15 (top panel), we illustrate this property for the inferred loci using the *CMT2* locus as an example. The MDS-plot visualizes the distribution of the *CMT2_STOP_* allele across the population structure present in the RegMap collection using the pairwise genome-wide relationship between the accessions based on the first two principle-components of the kinship matrix. The link between the geographic origin of the accessions and kinship is visualized by coloring the dots for each accession based on geographic origin. As expected, accessions from nearby regions (*e.g.* UK, Scandinavia and mainland Europe) are more related. The *CMT2_STOP_* allele is, however, not heavily confounded with population-structure and is present in most major sub-groups of the population (albeit with a higher frequency in Asia-see Figure S36).

#### Mixed models based vGWAS analyses to account for population structure via modeling of genome-wide kinship

We statistically deal with the strong correlations that exist between climate & population structure (e.g. along east-west/north-south clines) using a mixed-model based approach accounting for genomic kinship combined with genomic control. This approach has earlier been shown to control type I errors (*i.e.* genome-wide P-value inflation) in structured populations. The major challenge in analyzing this population is thus not the false-positive rate, as also standard GWAS analyses can be implemented in the same mixed-models framework, but rather to avoid unacceptably high type II errors (*i.e.* low power) for traits confounded with population-structure. Traditional GWAS analyses model alleles to have a linear relationship with climate, which in practice means that they mostly coincide with the population-structure along geographic clines. Hence, analyses will either be prone to identify false associations (when population-structure is not accounted for), or be under-powered (when accounting for population-structure). Although this is not explicitly discussed in the earlier reports based on this data, this is the primary reason for their lack of genome-wide significant associations to individual adaptive loci. As illustrated in Figure 3, the variance-heterogeneity test identifies loci present across population strata, where the signal therefore remain even after accounting for population structure via the mixed-model approach. The independence between the effect of the inferred locus and population structure can be evaluated statistically by fitting a linear mixed model where the genotype is regressed on the genomic kinship, where the heritability differs from 0 when confounding is present. For *CMT2* this estimate is zero, showing that the *CMT2* genotype is not confounded with population structure in this data.

**On the power of vGWAS and GWAS analyses in highly structured populations** There are several reasons for why a low overlap is expected between the results from traditional GWAS/selective-sweep analyses (as performed earlier) and the variance-heterogeneity GWAS (vGWAS) used here. First, in the absence of population stratification, the GWAS is more powerful than the vGWAS. In the presence of population stratification, however, loci affecting the mean phenotype will often be highly confounded with population-structure as they are a main genetic mechanism leading to local adaptation. In order to infer such loci when controlling for population structure, the same alleles need to have been under selection in multiple, unrelated populations, which is apparently a rare event as no such loci could be detected in the earlier studies of climate adaptation. The population genetics forces acting on variance-controlling loci are still poorly explored. Studies have, however, shown that - and low-variance alleles are likely to co-exist in the population over extended periods of time at a frequency balanced depending on fluctuations in the surrounding environment that the population adapts to^11^. Due to this, both alleles are more likely to be present across different population strata than mean affecting alleles and therefore be less confounded with population structure. In Figure S36, we exemplify this by visualizing the allele-frequency of the *CMT2_STOP_* allele across the major sub-populations in the RegMap and 1001-genomes data. Although the uneven sampling across the regions in the 1001-genomes makes allele-frequency estimates uncertain, the overall picture shows that minor allele is present across all sub-populations at a lower frequency and that it has increased in frequency in Asia.

Second, a traditional GWAS searches for difference in means between genotypes, whereas the vGWAS searches for differences in variances between genotypes. As these are two different statistical properties of the phenotypic distribution, the basic assumption is that they are both statistically and biologically unrelated and consequently the loci identified by the two methods are not expected to overlap. Although some degree of overlap might be expected, *e.g*. in situations where the variance scales with the mean, the high-significance required to reach genome-wide significance in the testing, in practice only loci with strong, pure effects on one of the statistical moments seem to be able to reach such significance levels^12^. Formal comparisons between the results in the association studies will thus be misleading to the readers, as these will indicate that an overlap is to be expected. Results from evaluations of the overlap for sub-GWAS signals to explore the potential overlap of loci with weaker effects on both the mean and the variance shows some overlap (Figure S40–S44). It should be noted, however, that comparisons of overlap at individual loci is not appropriate at these significance levels due to the lack of proper control of the type I error rate. The overall conclusions from these comparisons is i) that the power is generally very low for the GWAS after control for population stratification and ii) that even the sub-GWAS overlap is low for the two methods, but the overlap that exists is consistent with the correlation between the climate variables.

Third, the earlier studies have also inferred loci using traditional selective-sweep mapping. These analyses are designed to infer hard selective-sweeps where (potentially) adaptive alleles are assumed to have increased in frequency due to directional selection. As discussed above, the population genomic dynamics of plastic alleles does not follow the same pattern as for alleles affecting the mean^11^, leading to a co-existence of the alleles over prolonged periods of time. This means that they will not be surrounded by a traditional genomic footprint of directional selection that can be detected in a selective-sweep analysis and one would not expect any overlap between the loci inferred in the selective sweep and vGWAS based analyses.

---

**Supplementary Figure 1.**
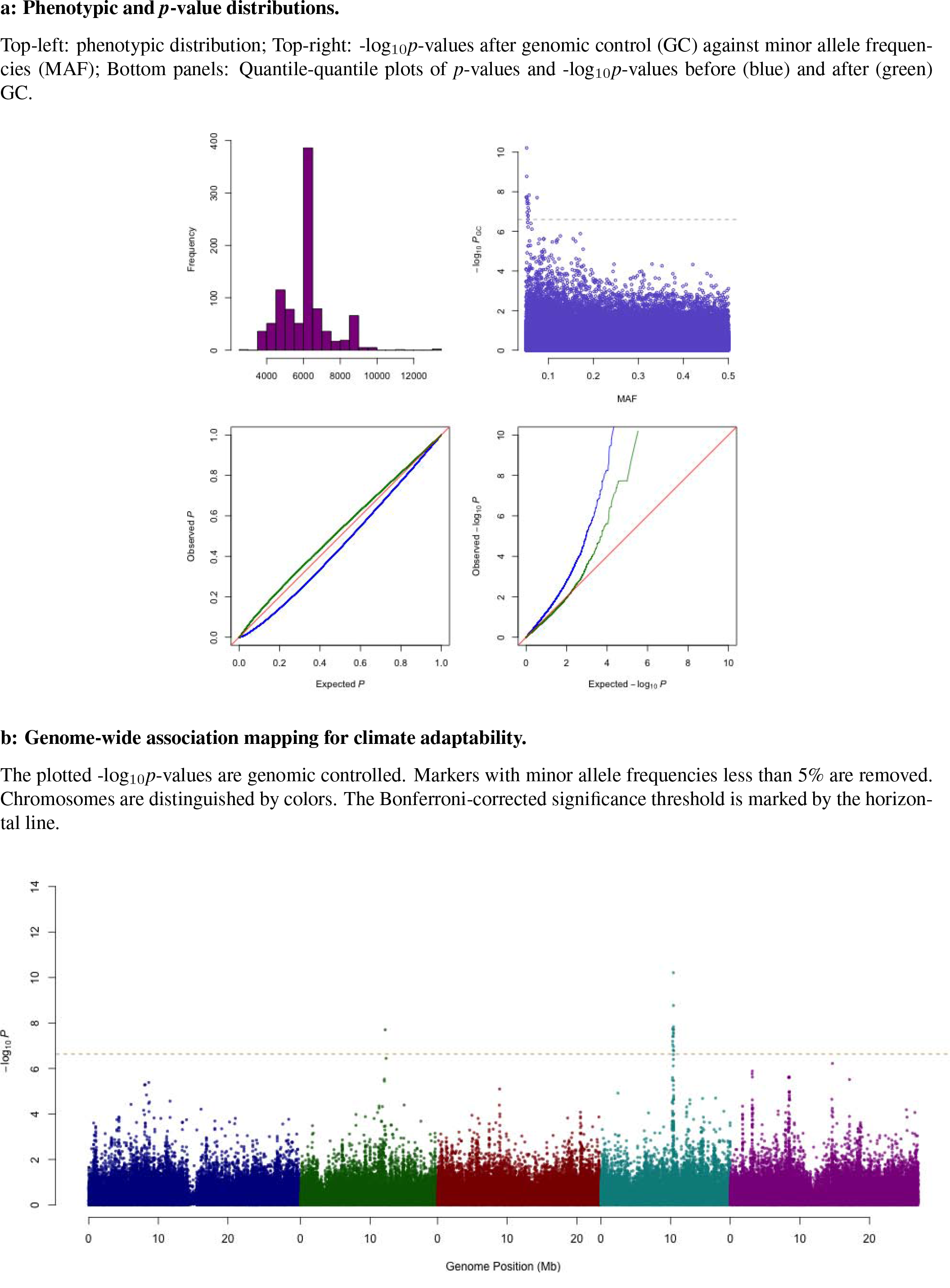
Summary of results for temperature seasonality.

**Supplementary Figure 2.**
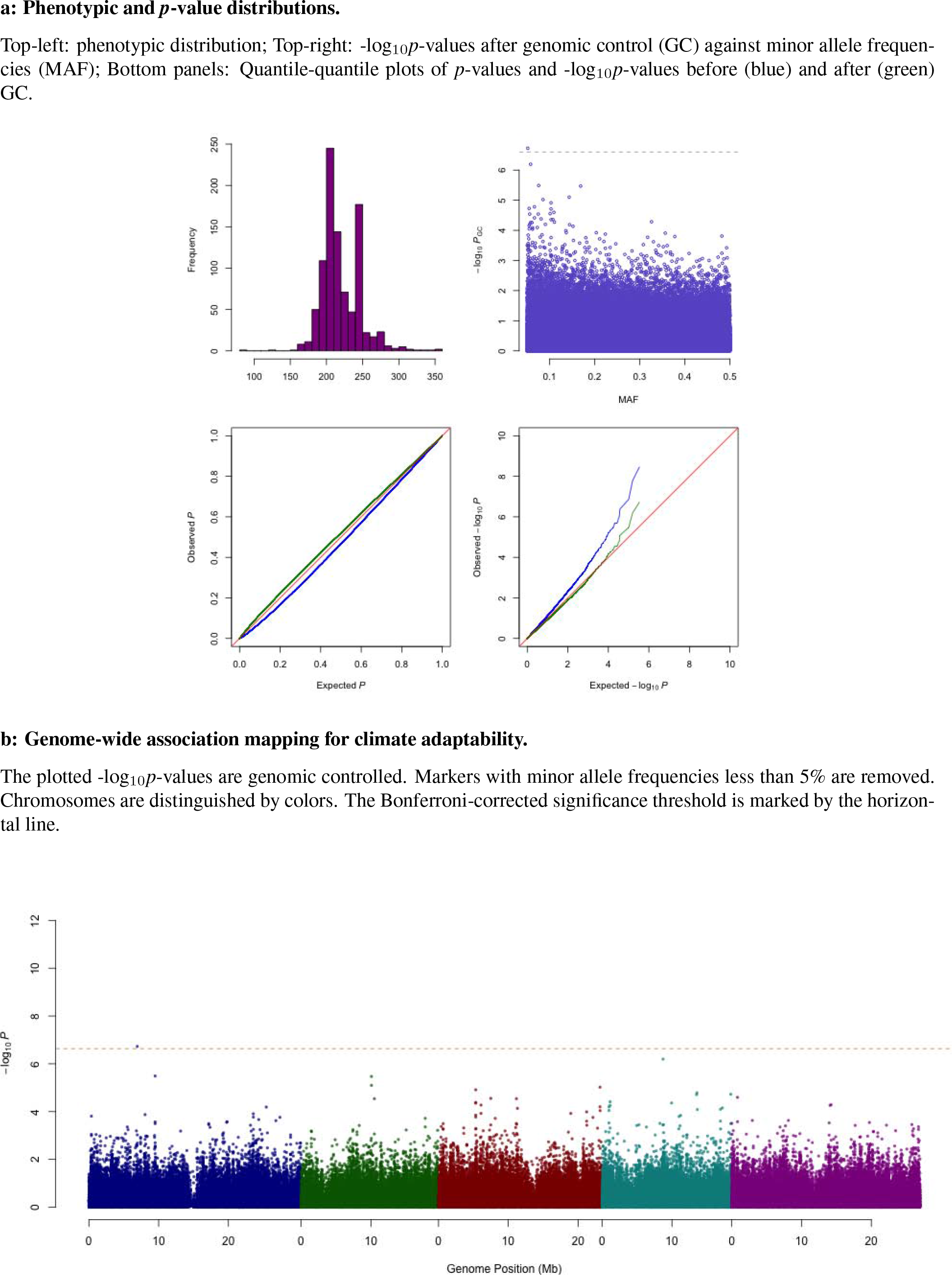
Summary of results for maximum temperature in the warmest month.

**Supplementary Figure 3.**
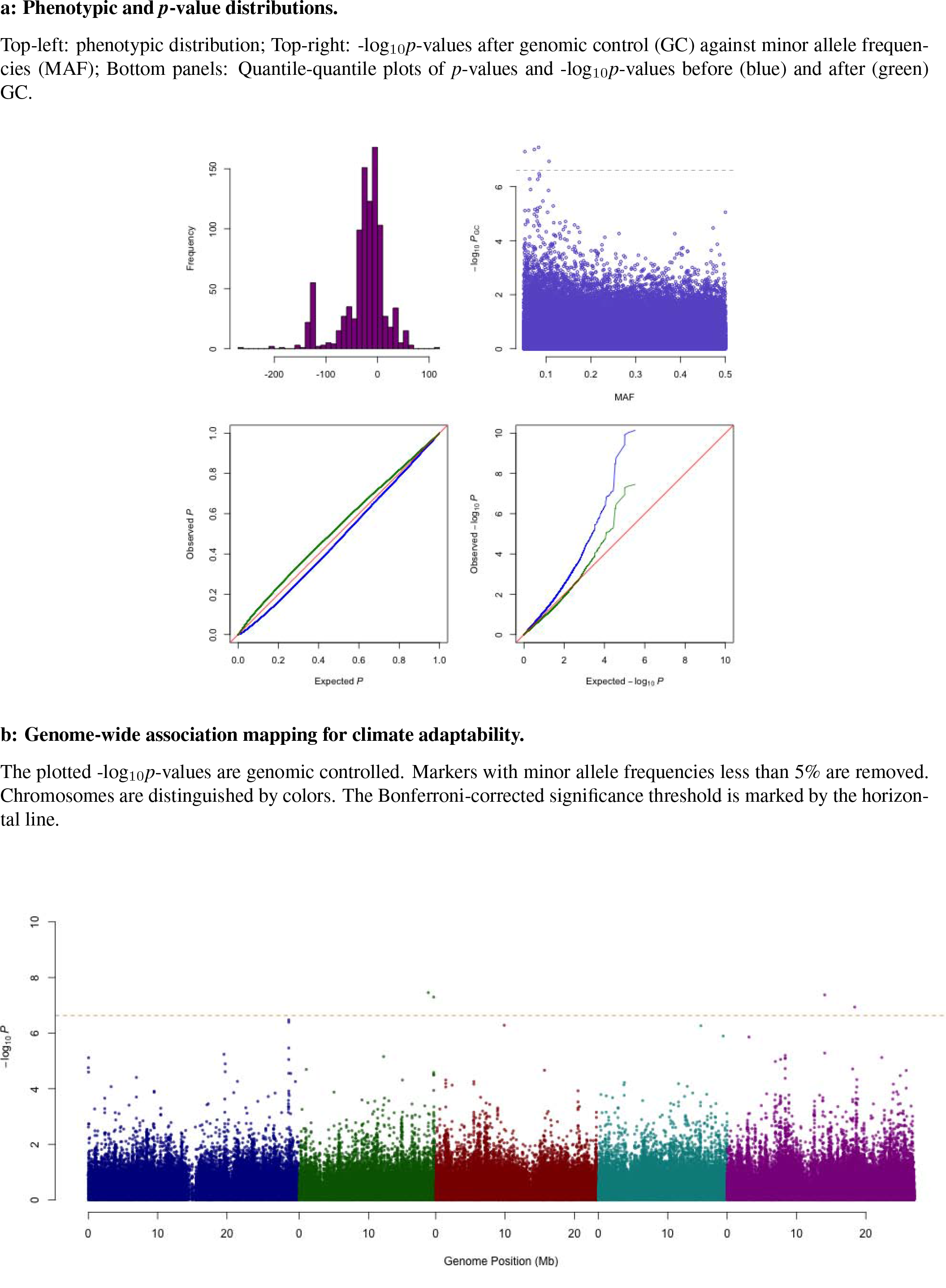
Summary of results for minimum temperature in the coldest month.

**Supplementary Figure 4.**
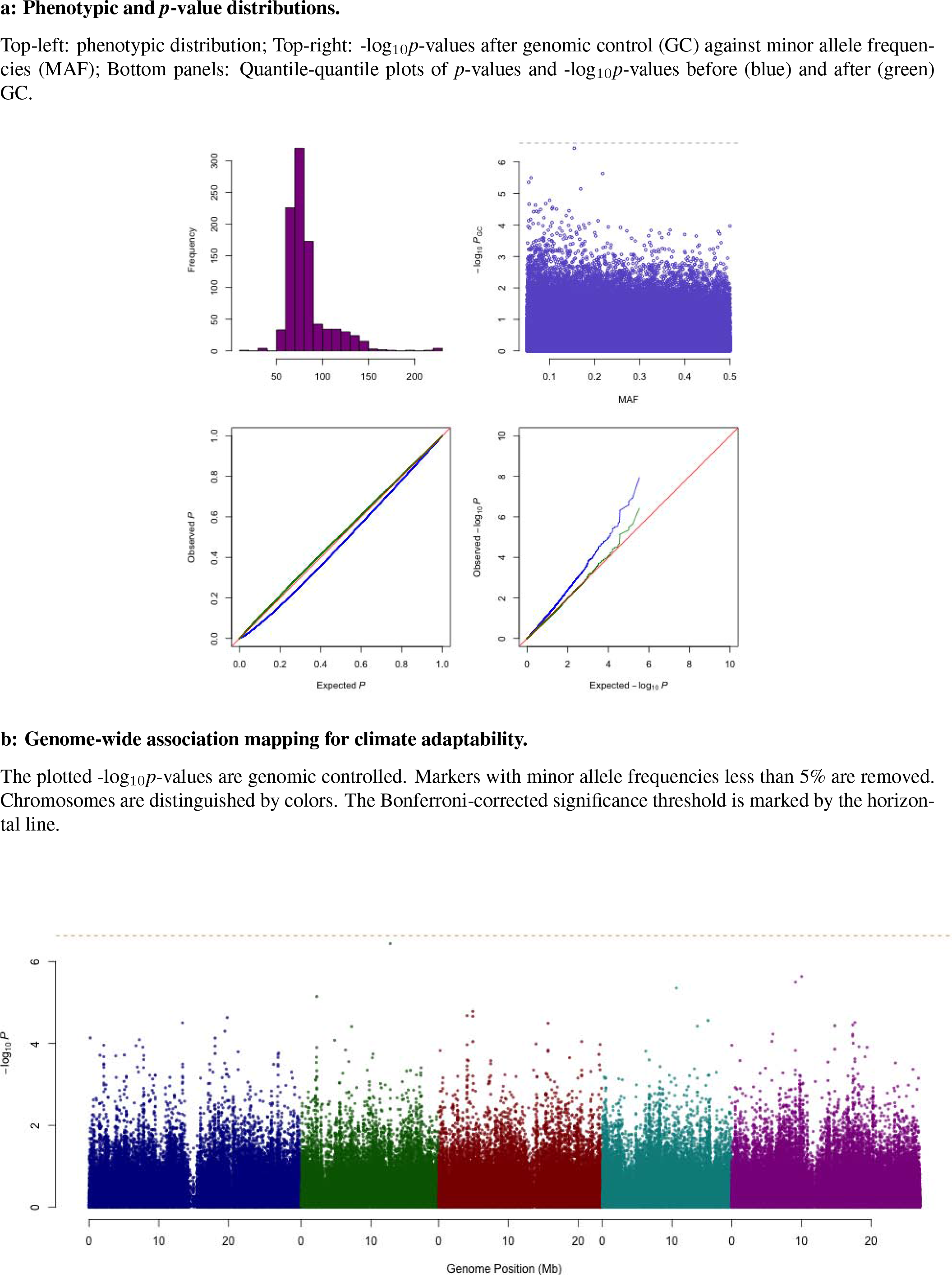
Summary of results for precipitation in the wettest month.

**Supplementary Figure 5.**
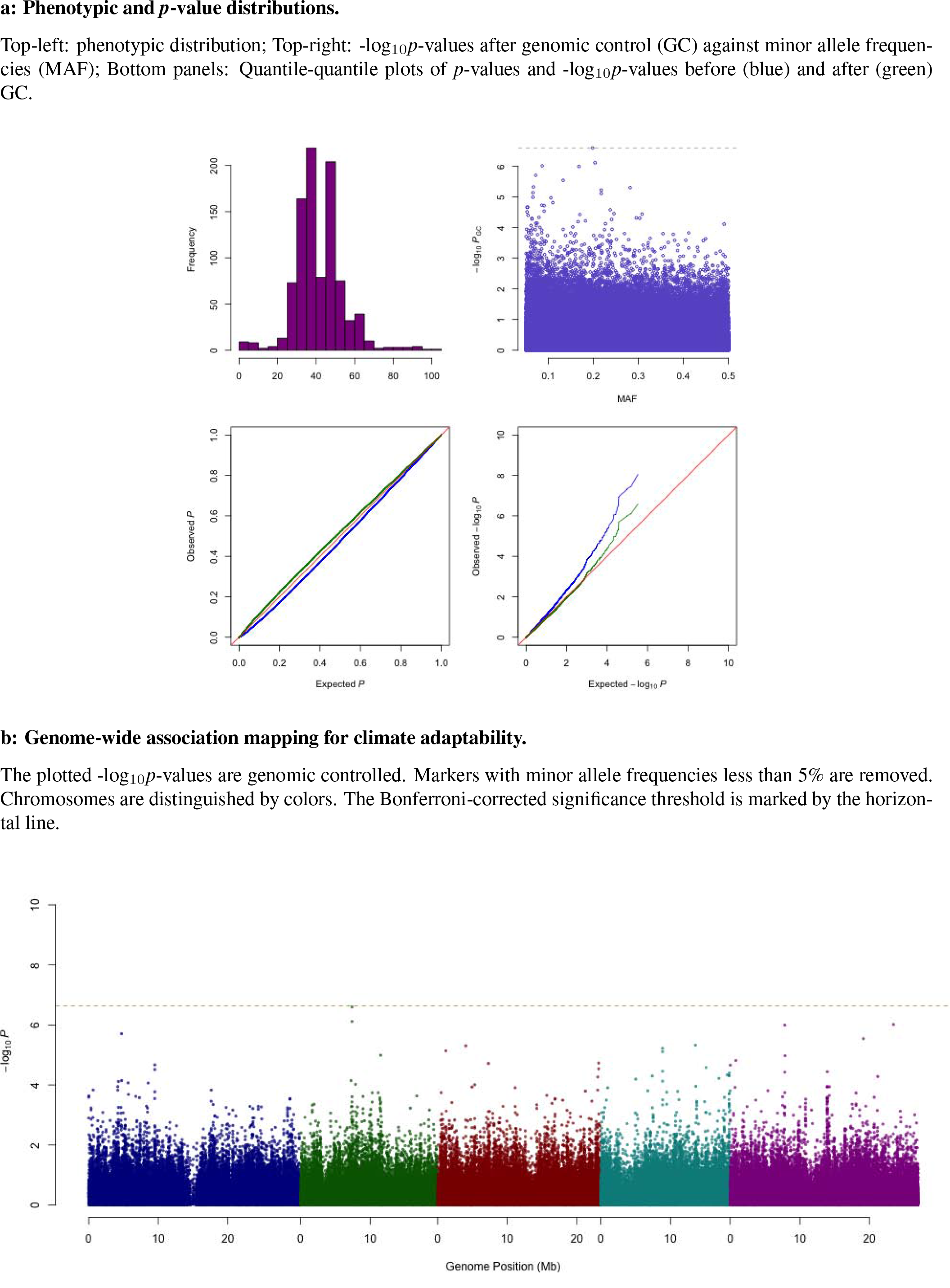
Summary of results for precipitation in the driest month.

**Supplementary Figure 6.**
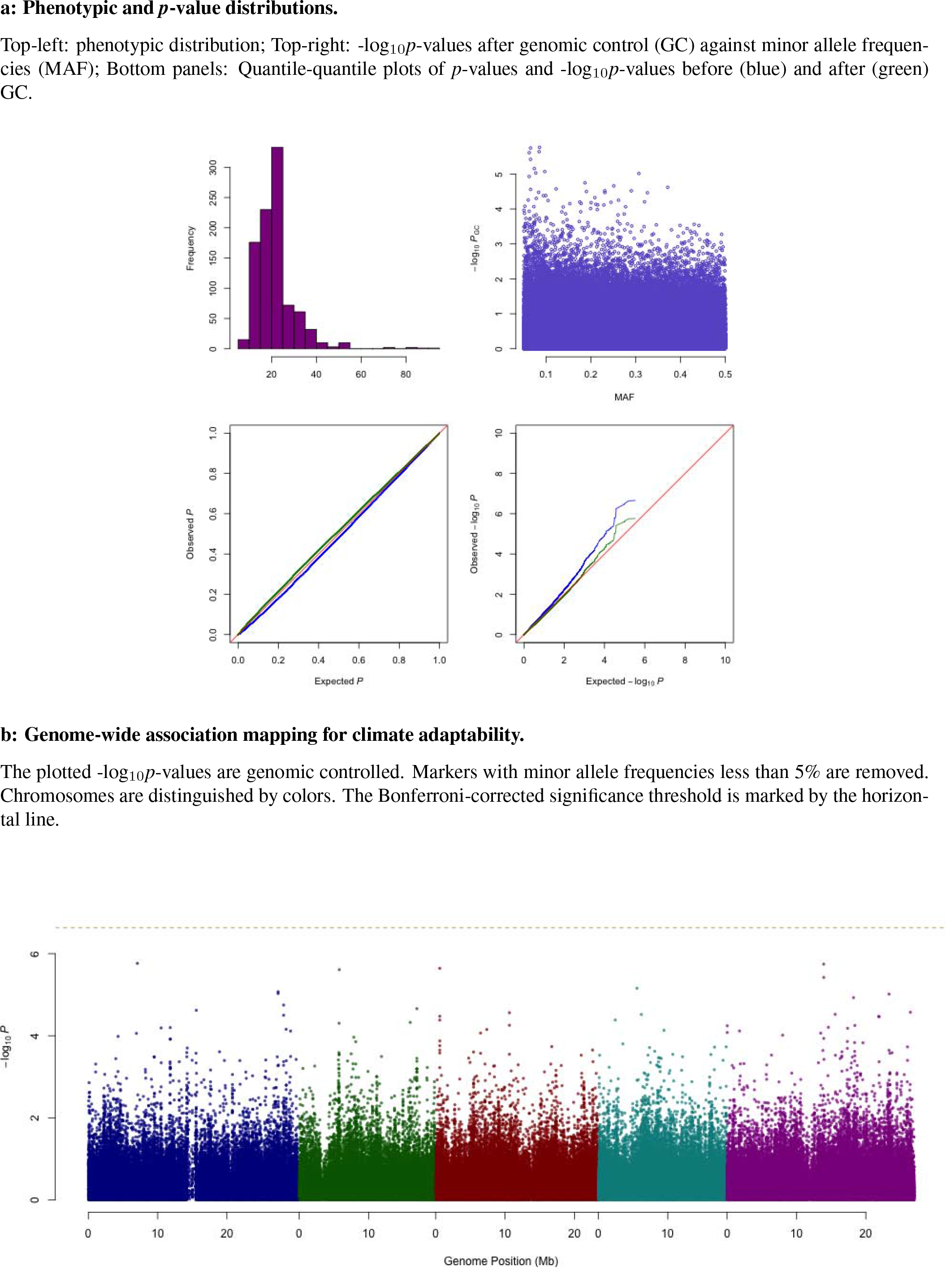
Summary of results for precipitation CV.

**Supplementary Figure 7.**
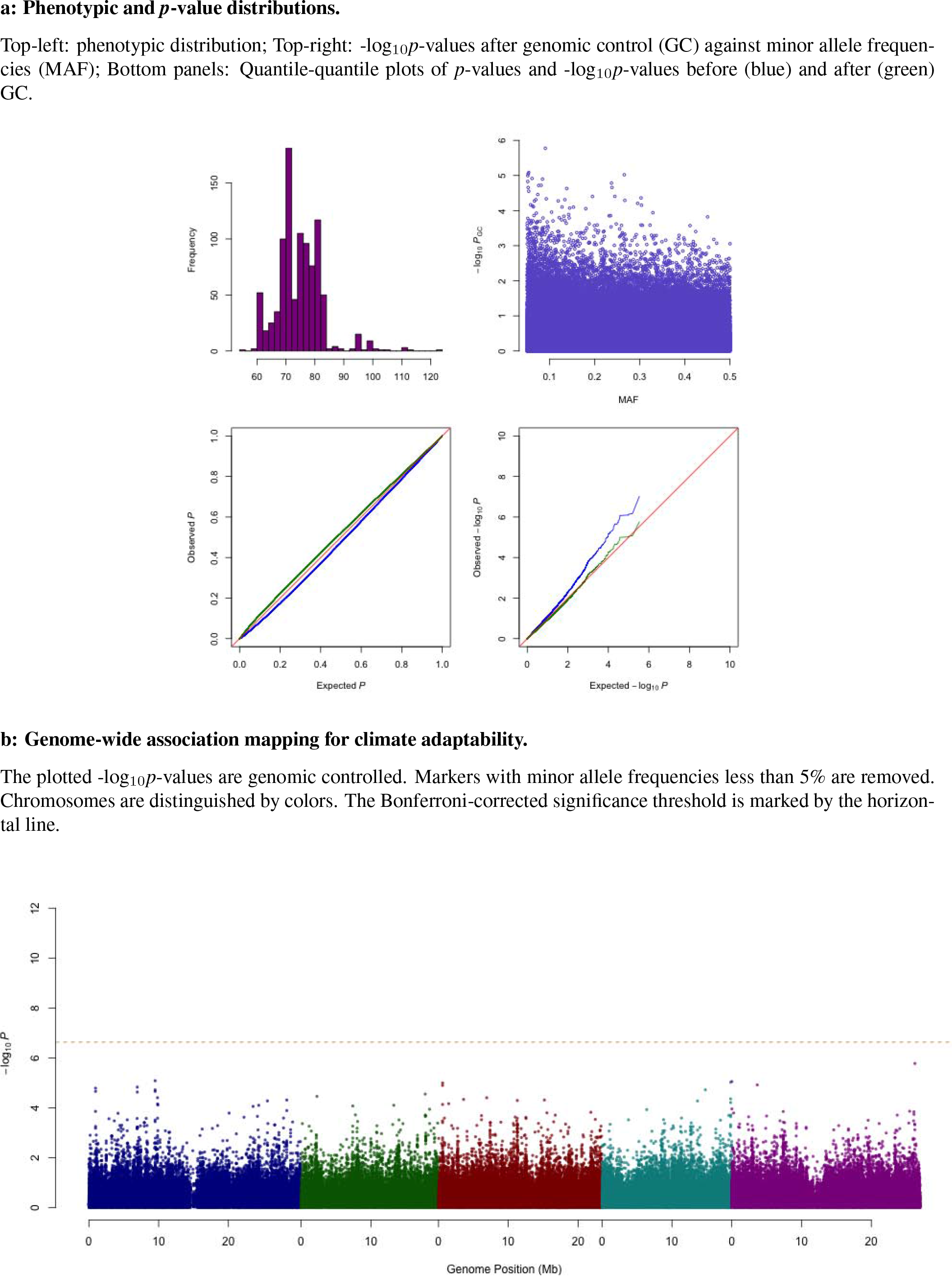
Summary of results for photosynthetically active radiation in spring.

**Supplementary Figure 8.**
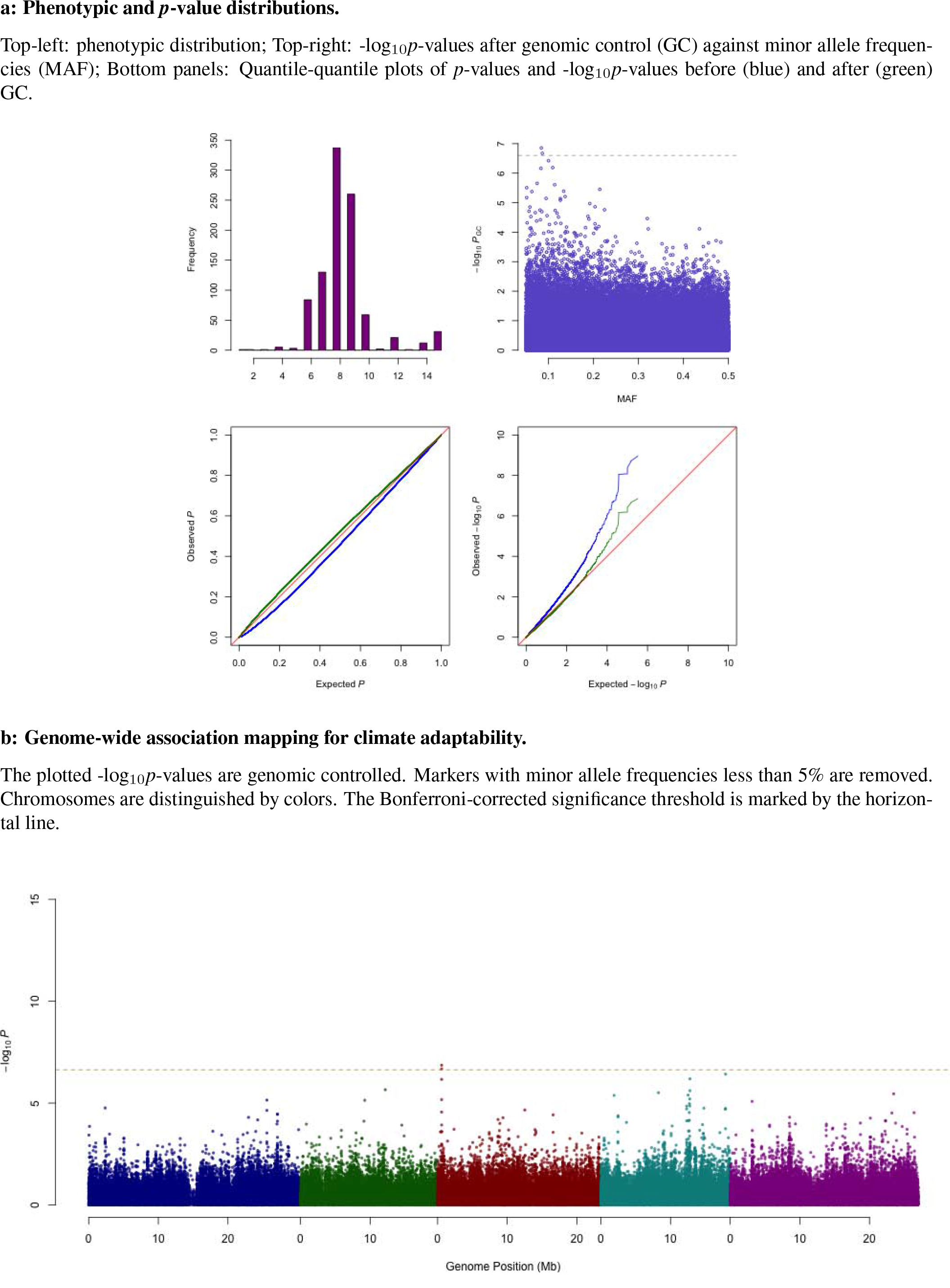
Summary of results for length of the growing season.

**Supplementary Figure 9.**
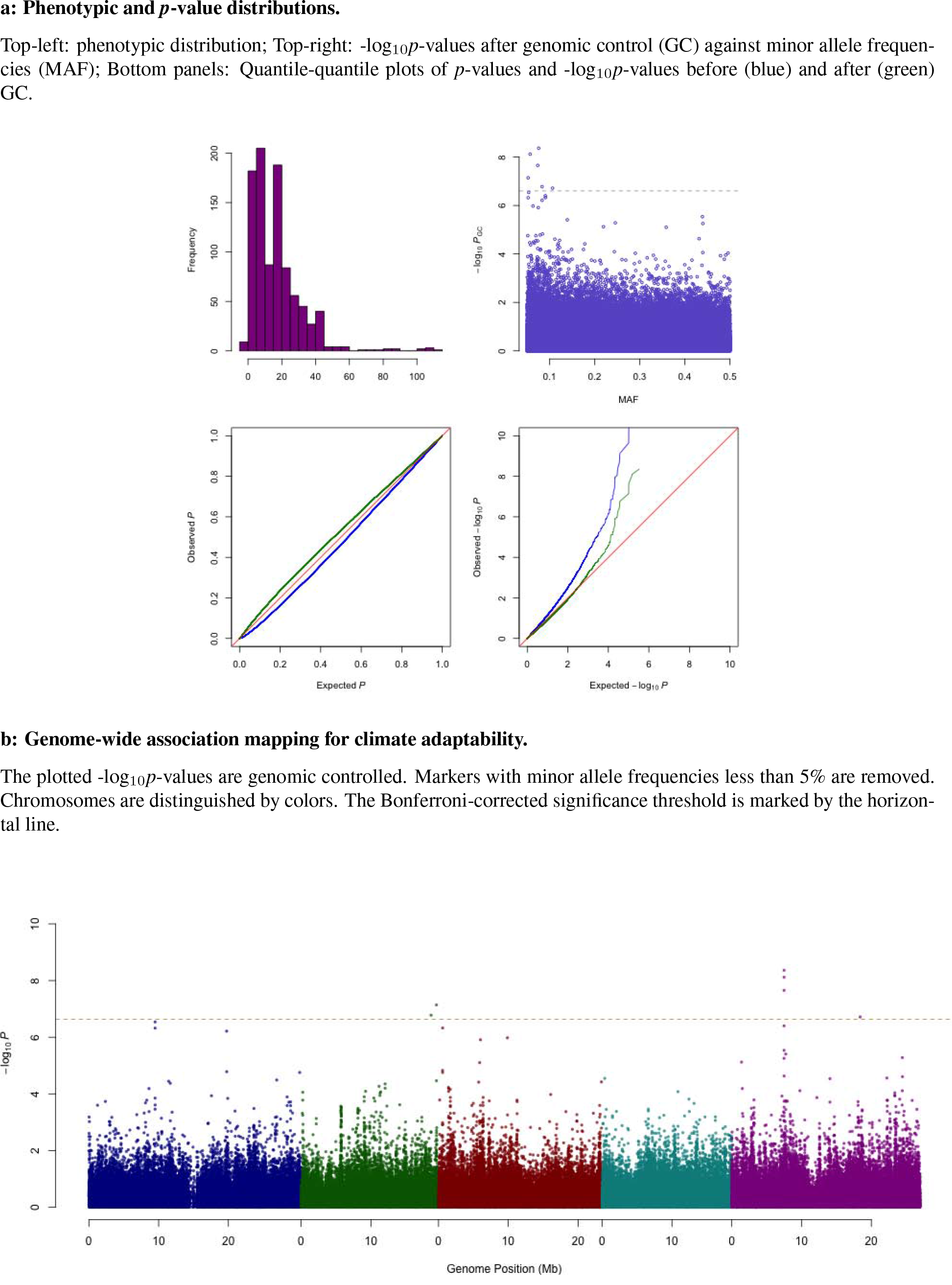
Summary of results for number of consecutive cold days.

**Supplementary Figure 10.**
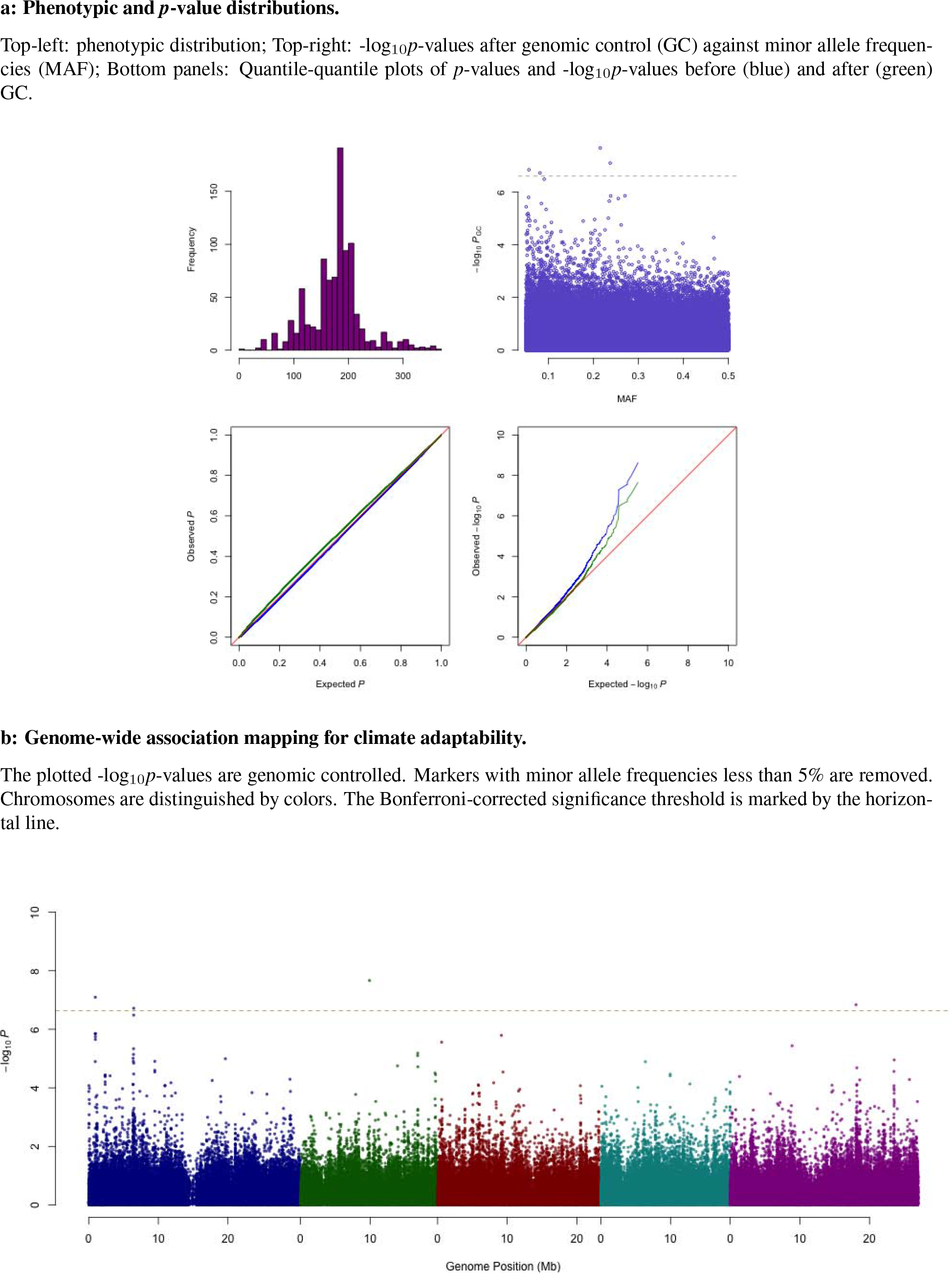
Summary of results for number of consecutive frost-free days.

**Supplementary Figure 11.**
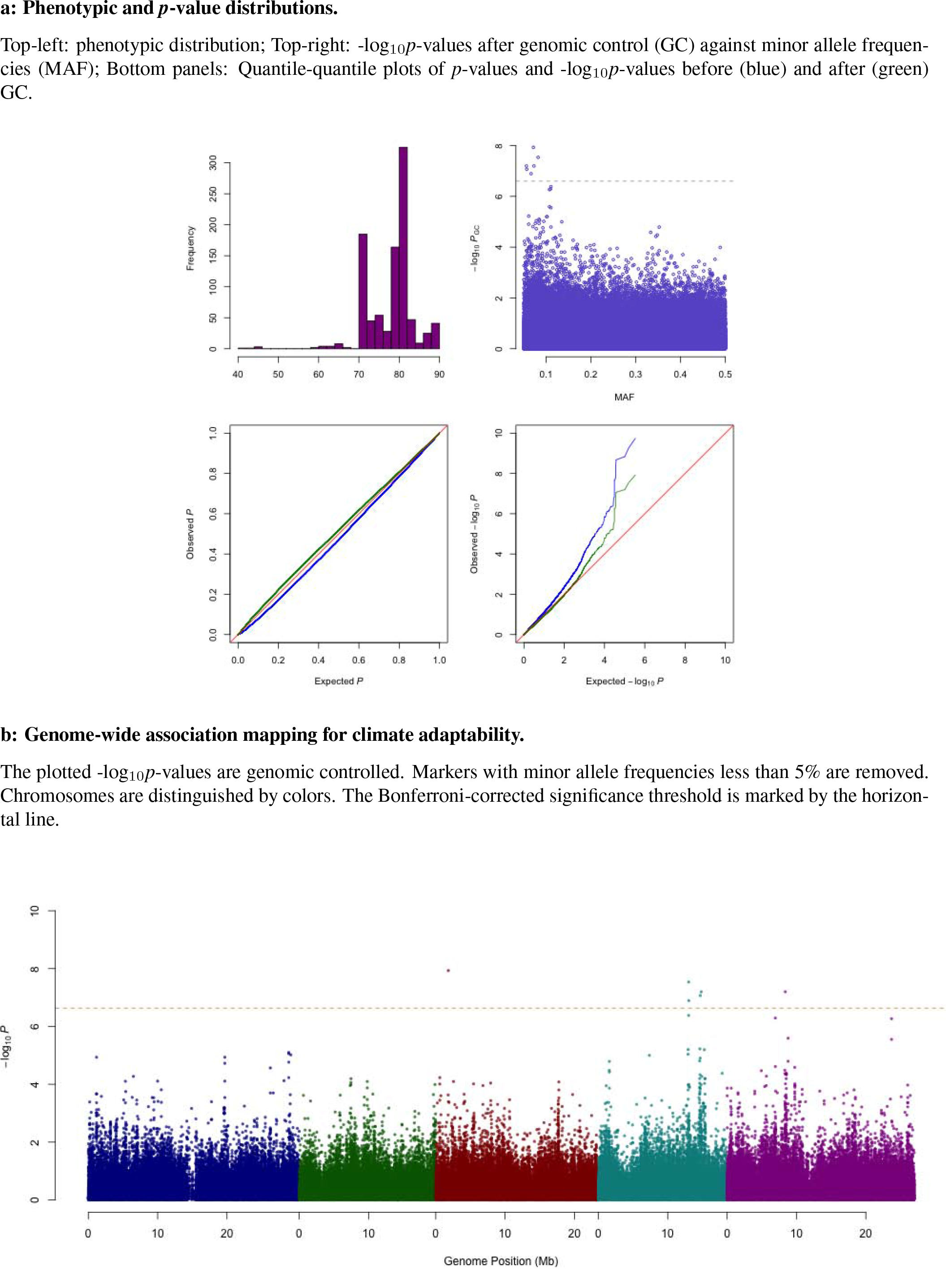
Summary of results for relative humidity in spring.

**Supplementary Figure 12.**
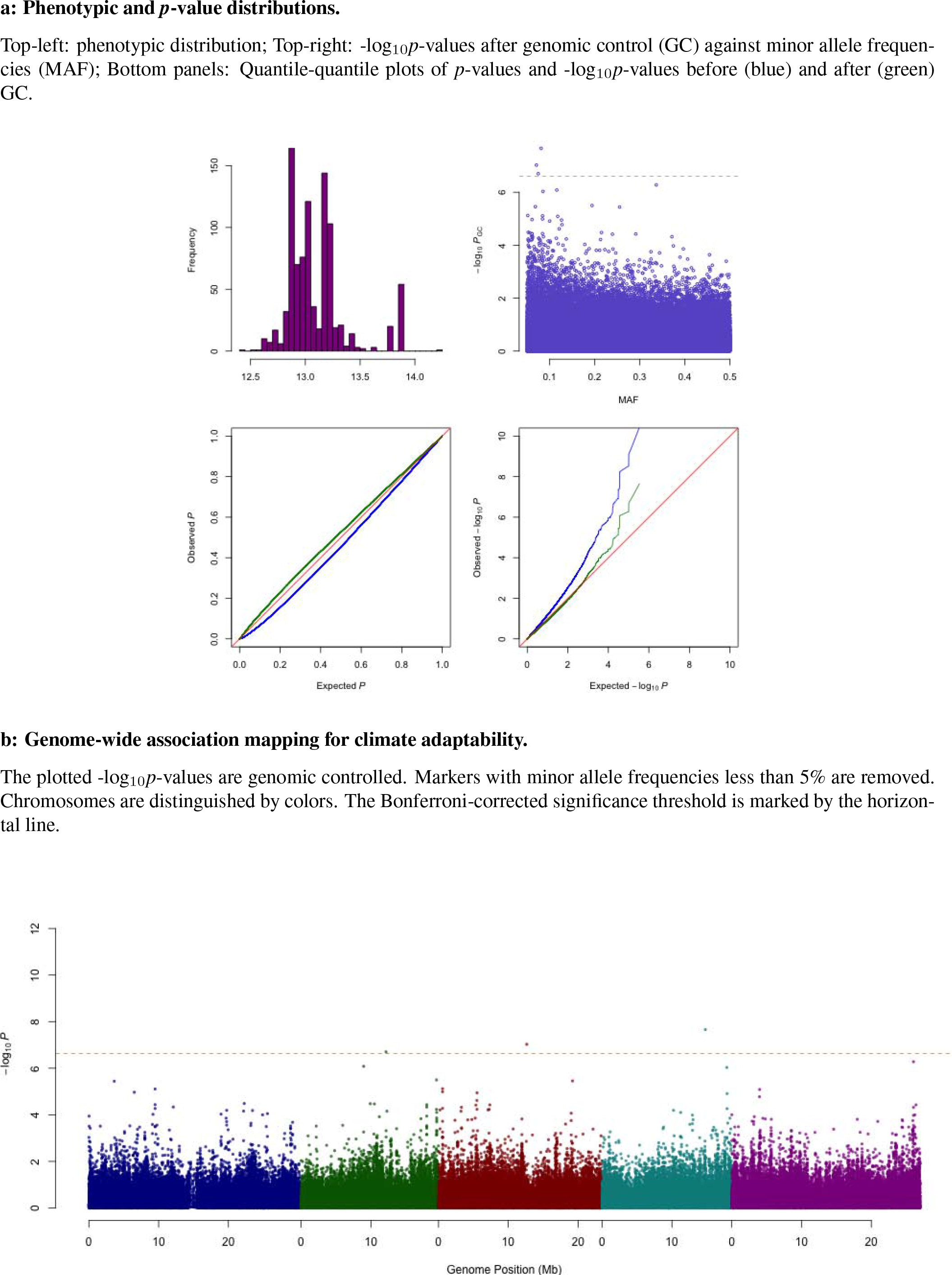
Summary of results for daylength in spring.

**Supplementary Figure 13.**
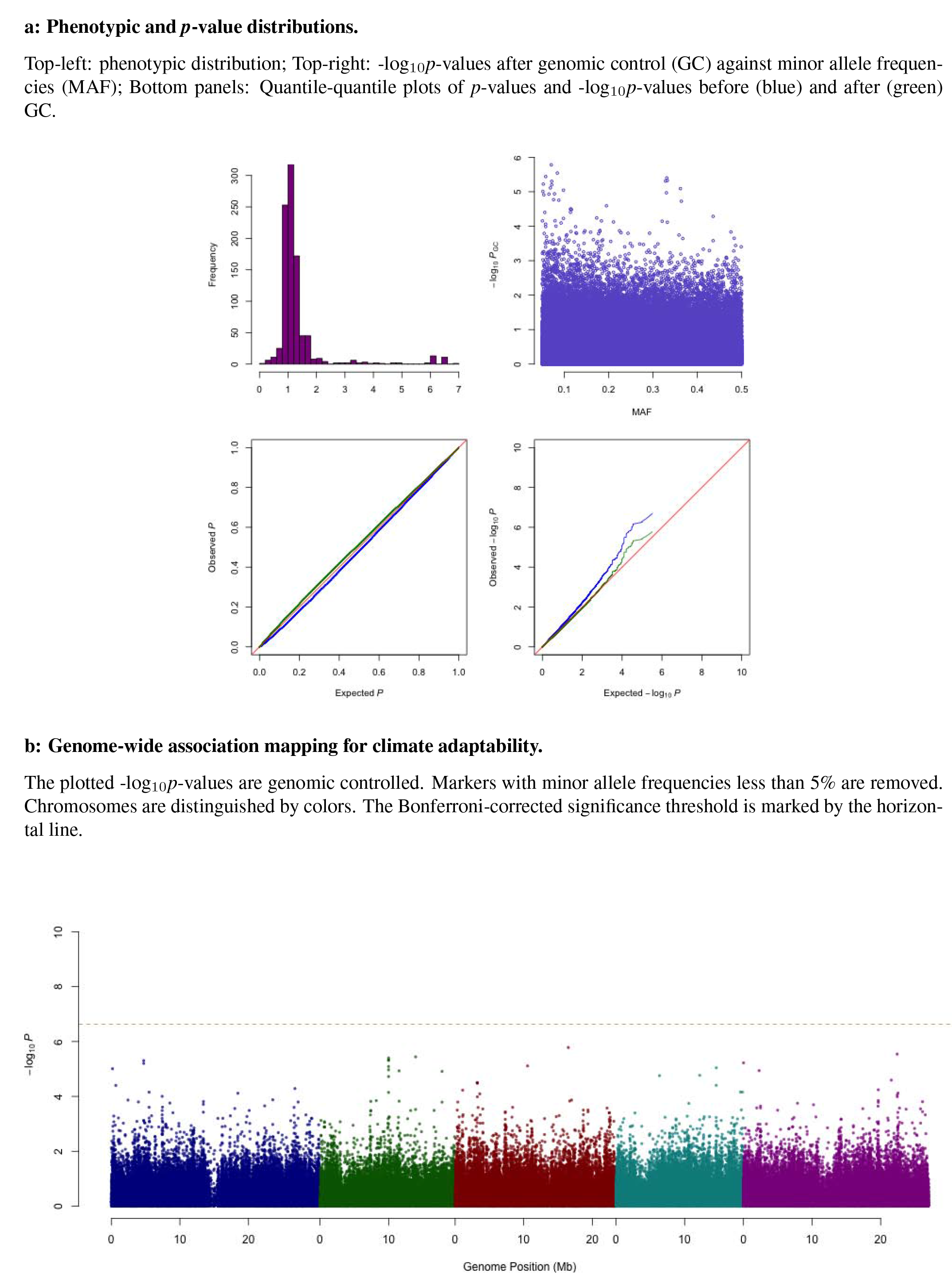
Summary of results for aridity index.

**Supplementary Figure 14.**
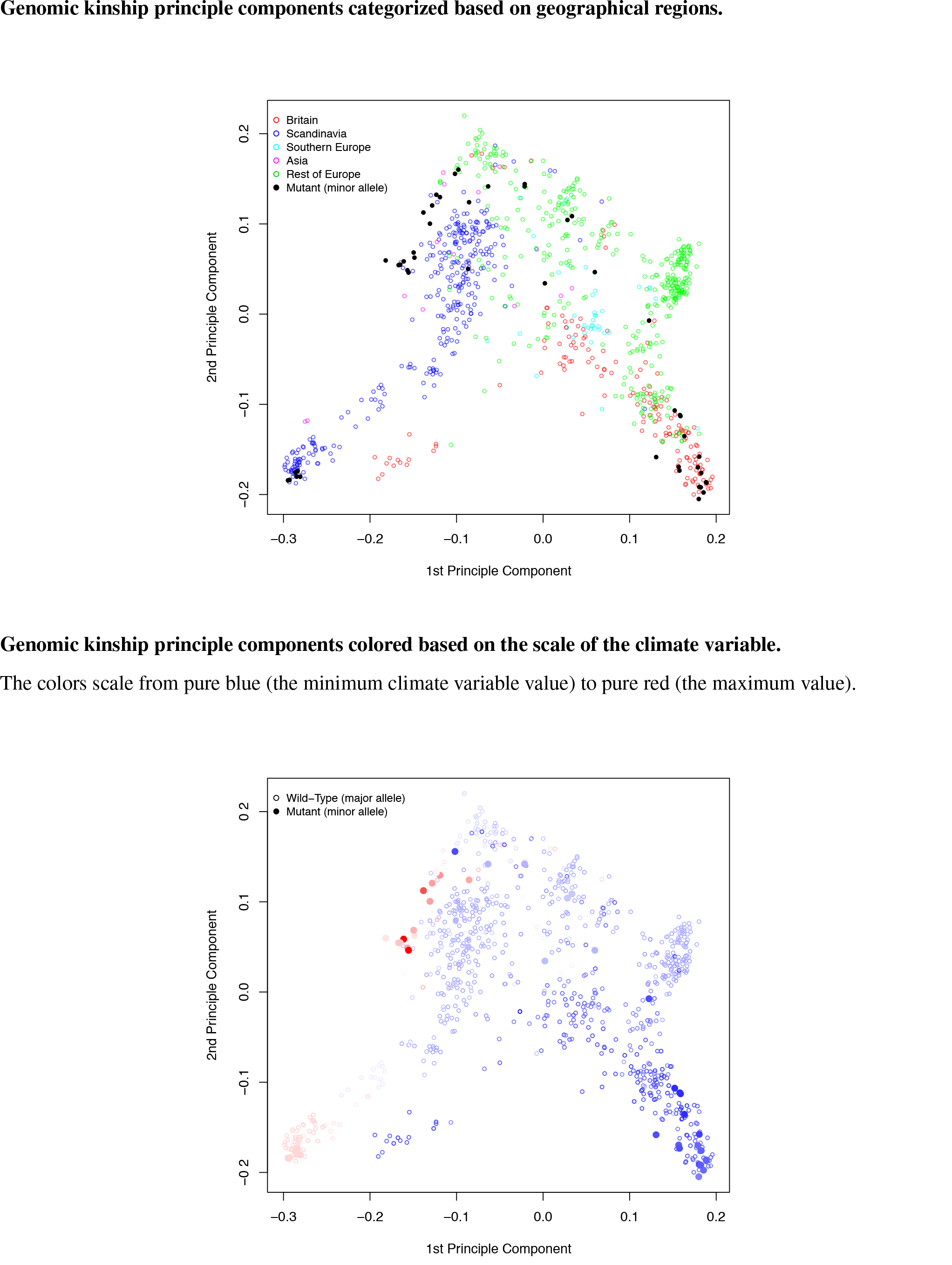
Principle components of the genomic kinship for the two alleles on chromosome 2 at 12169701 bp. Corresponding climate variable: temperature seasonality.

**Supplementary Figure 15.**
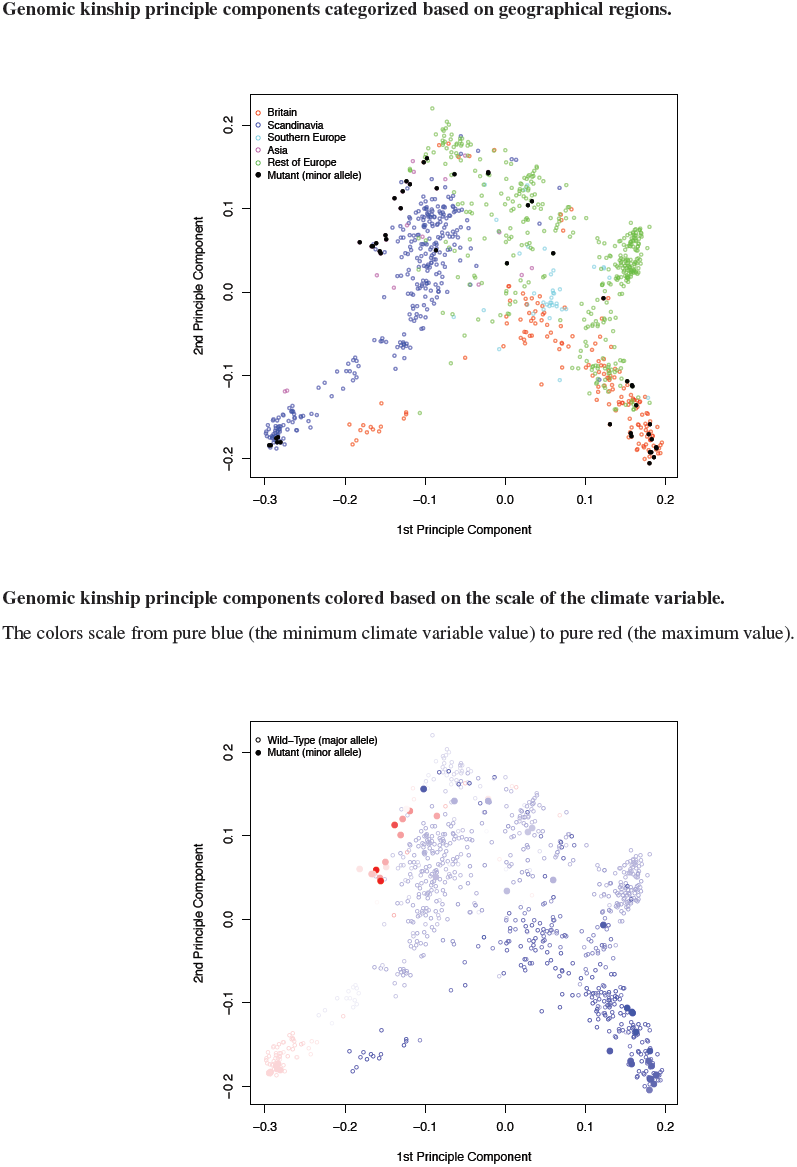
Principle components of the genomic kinship for the two alleles on chromosome 4 at 10406018 bp. Corresponding climate variable: temperature seasonality.

**Supplementary Figure 16.**
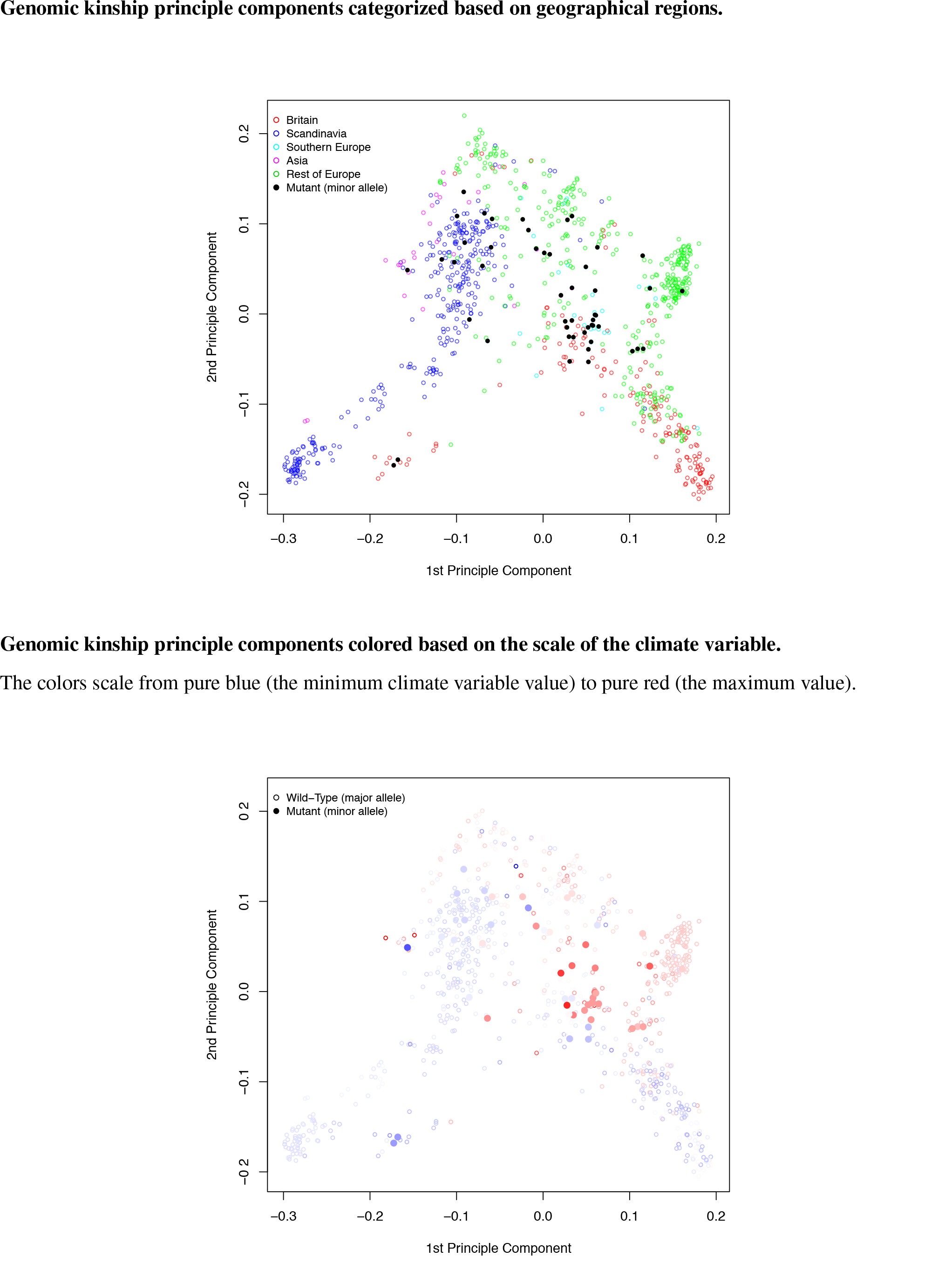
Principle components of the genomic kinship for the two alleles on chromosome 1 at 6936457 bp. Corresponding climate variable: maximum temperature in the warmest month.

**Supplementary Figure 17.**
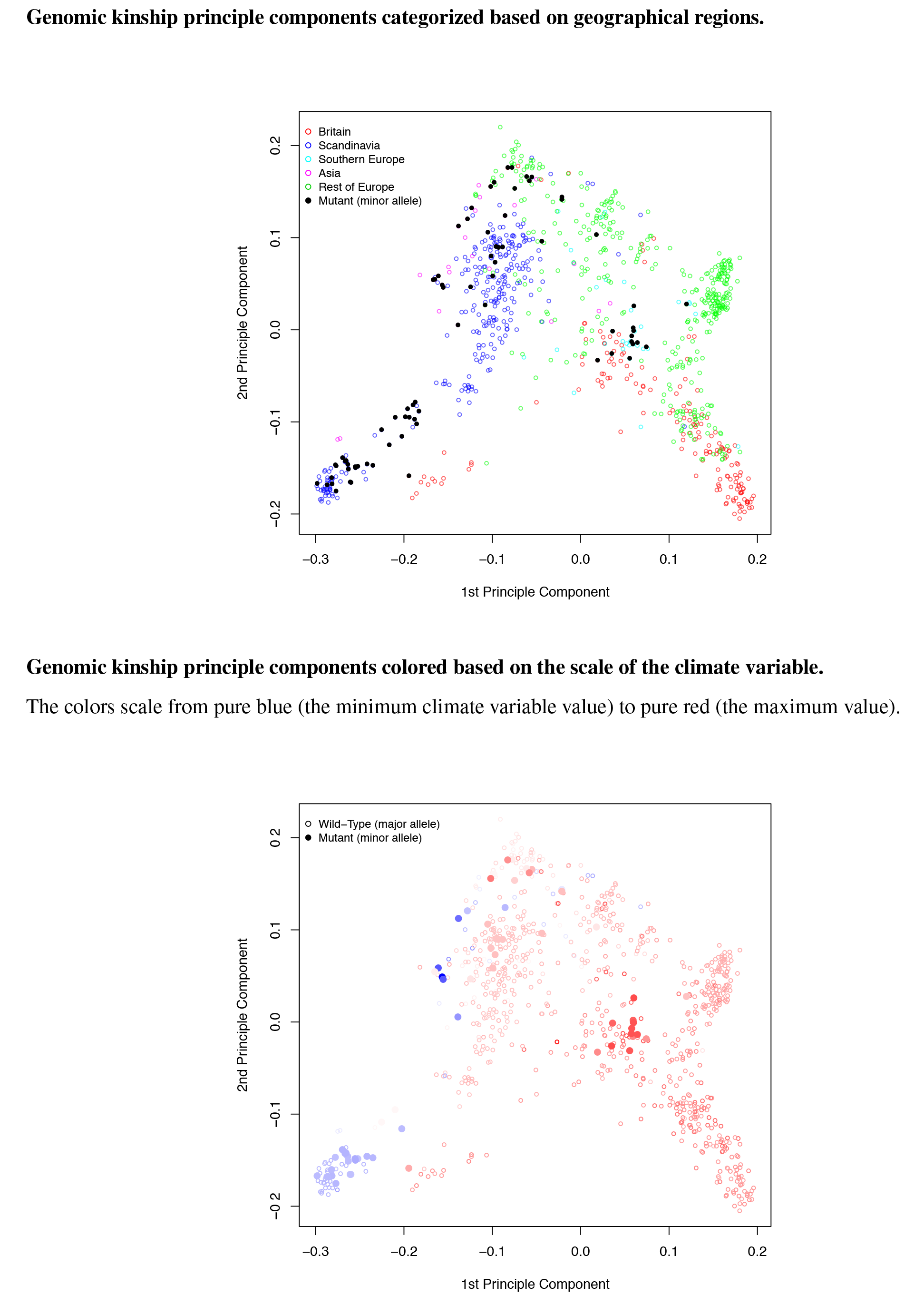
Principle components of the genomic kinship for the two alleles on chromosome 2 at 18620697 bp. Corresponding climate variable: minimum temperature in the coldest month.

**Supplementary Figure 18.**
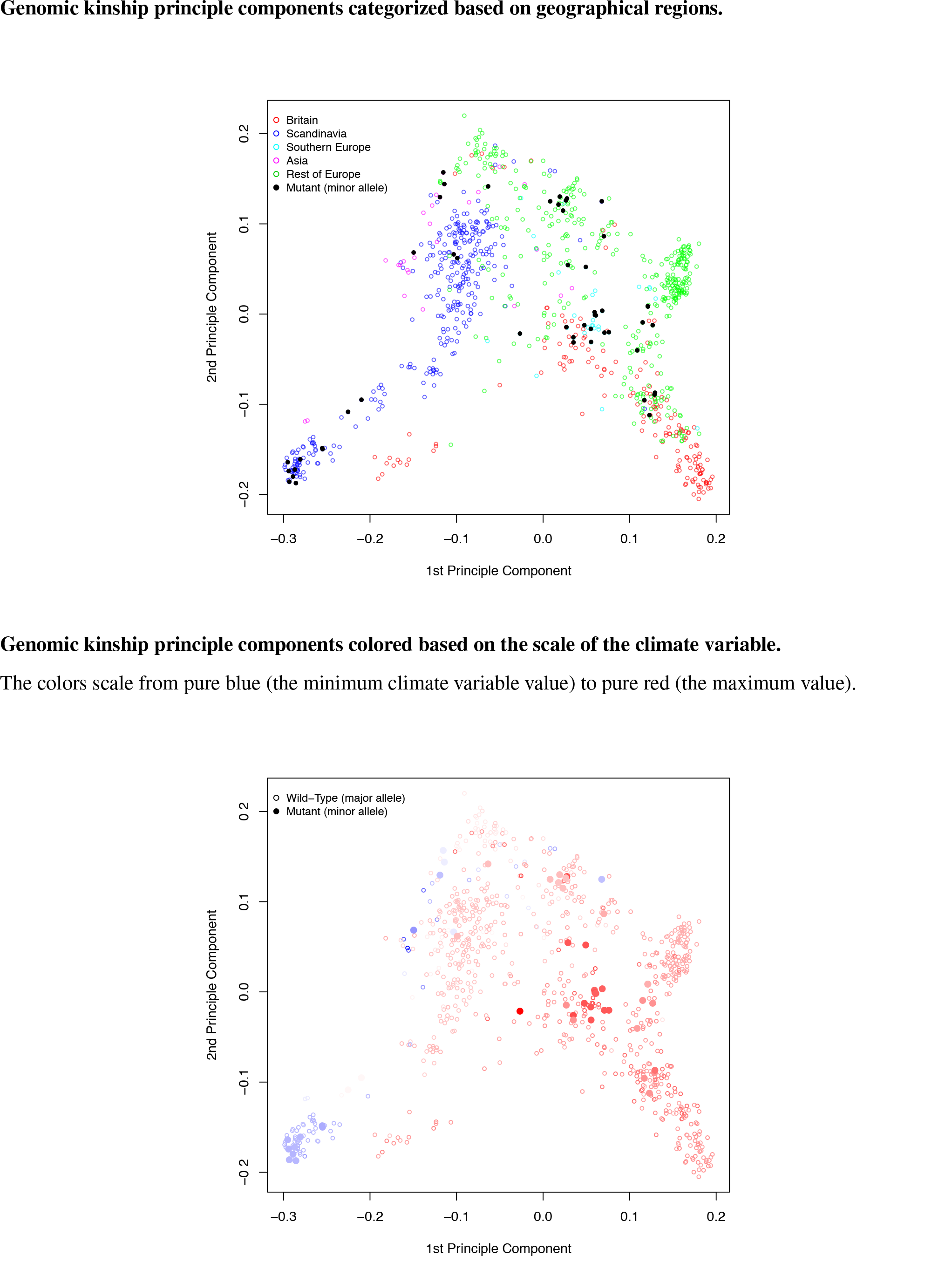
Principle components of the genomic kinship for the two alleles on chromosome 2 at 19397389 bp. Corresponding climate variable: minimum temperature in the coldest month.

**Supplementary Figure 19.**
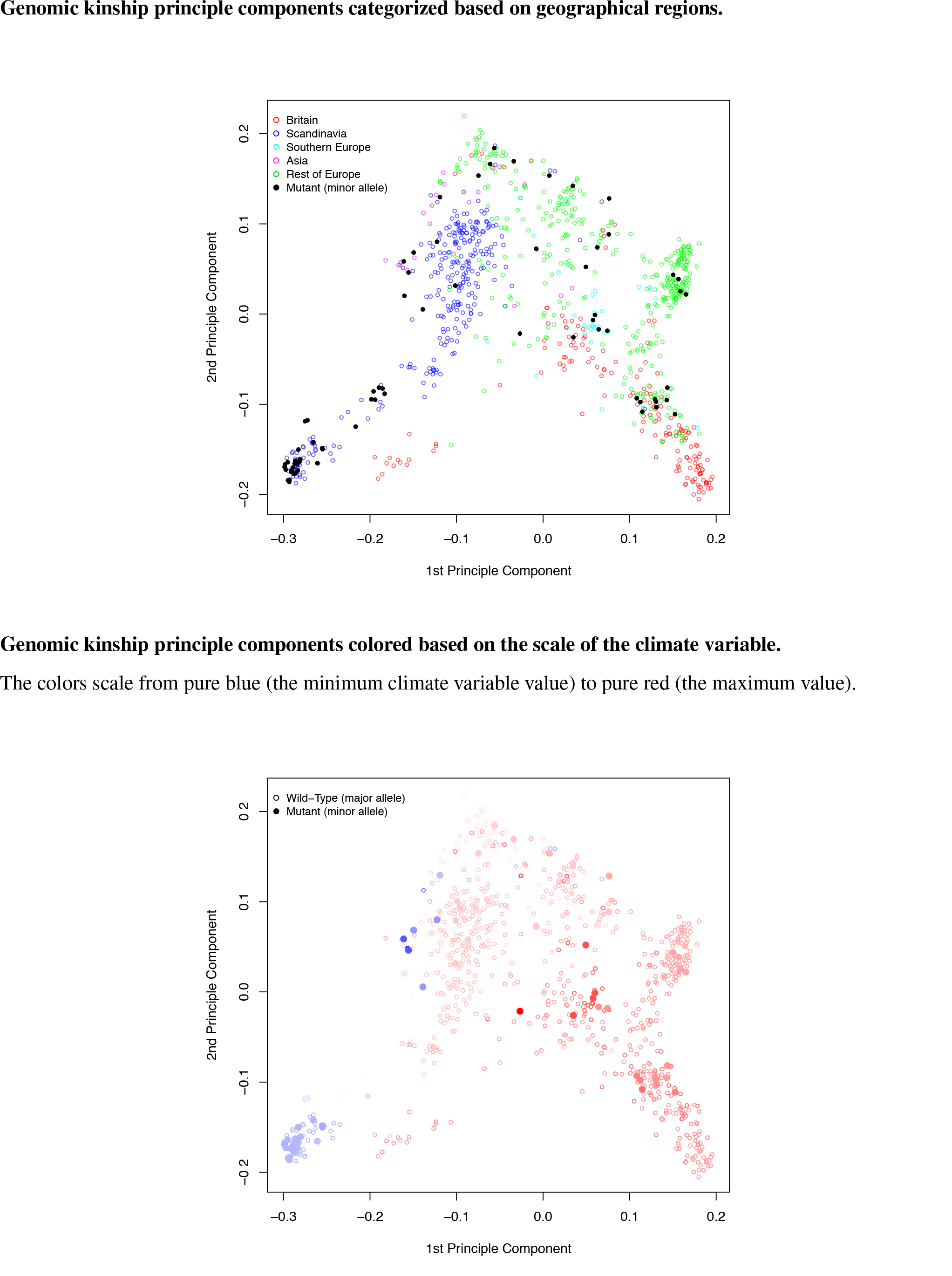
Principle components of the genomic kinship for the two alleles on chromosome 5 at 14067526 bp. Corresponding climate variable: minimum temperature in the coldest month.

**Supplementary Figure 20.**
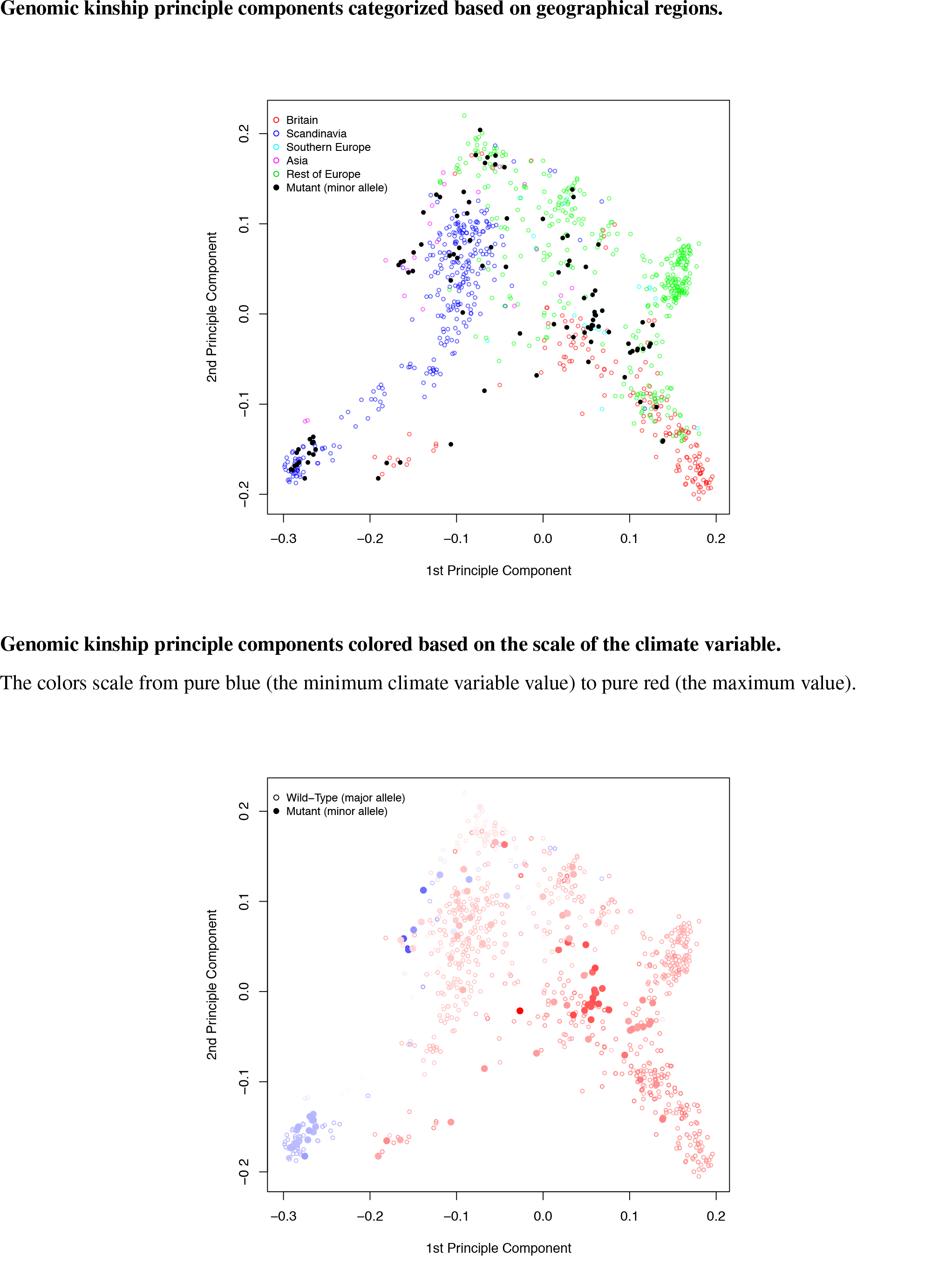
Principle components of the genomic kinship for the two alleles on chromosome 5 at 18397418 bp. Corresponding climate variable: minimum temperature in the coldest month.

**Supplementary Figure 21.**
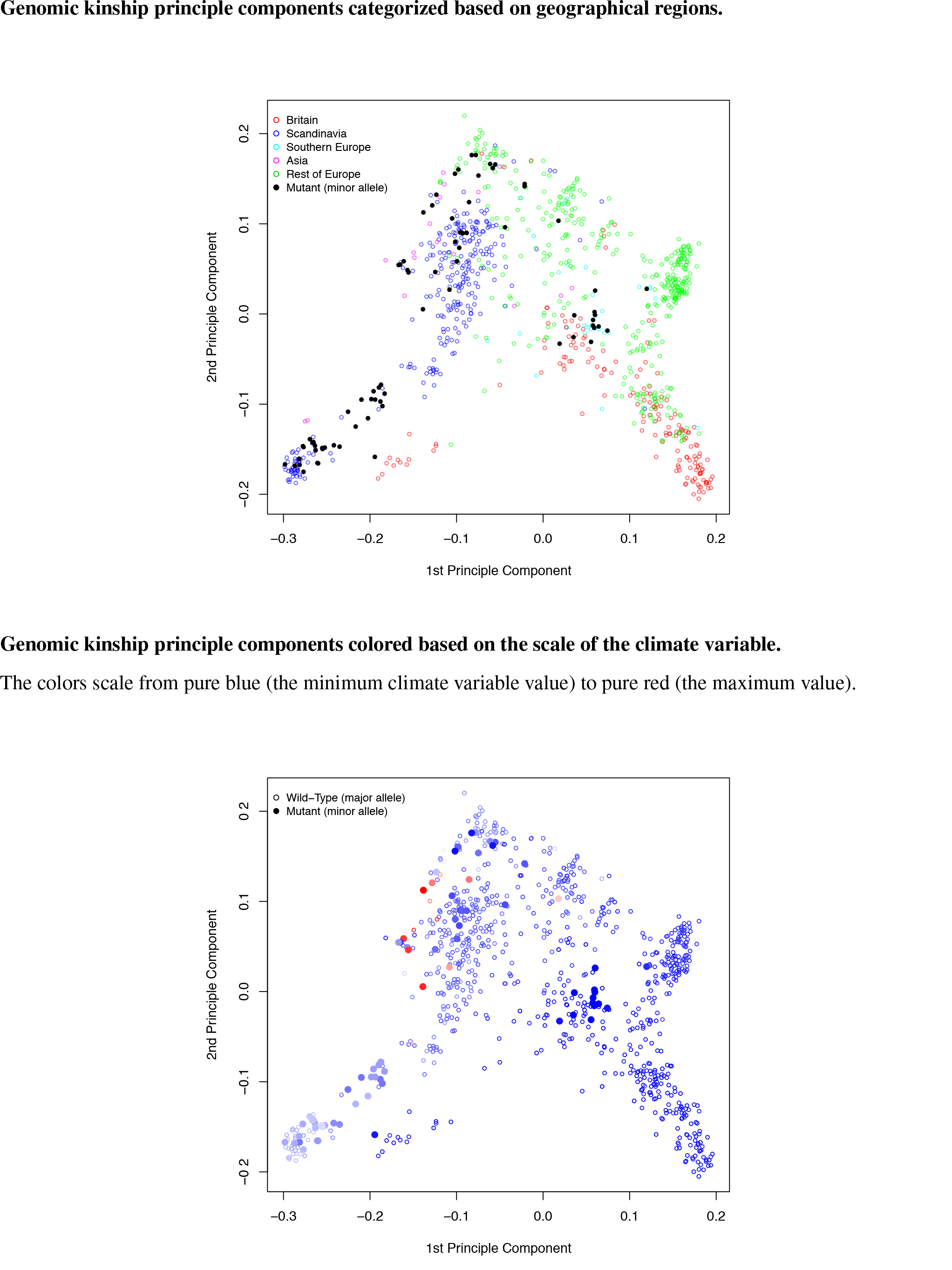
Principle components of the genomic kinship for the two alleles on chromosome 2 at 18620697 bp. Corresponding climate variable: number of consecutive cold days.

**Supplementary Figure 22.**
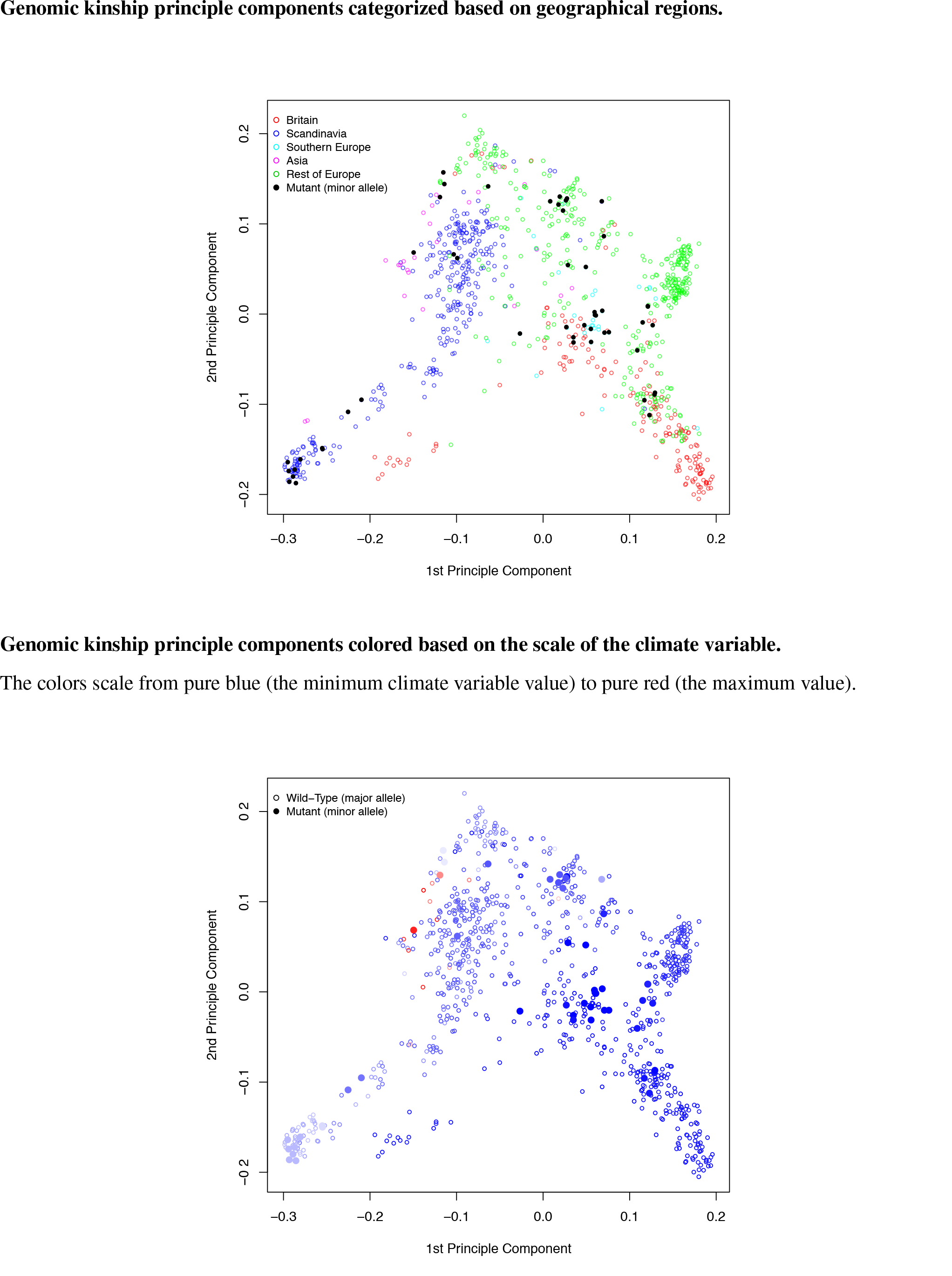
Principle components of the genomic kinship for the two alleles on chromosome 2 at 19397389 bp. Corresponding climate variable: number of consecutive cold days.

**Supplementary Figure 23.**
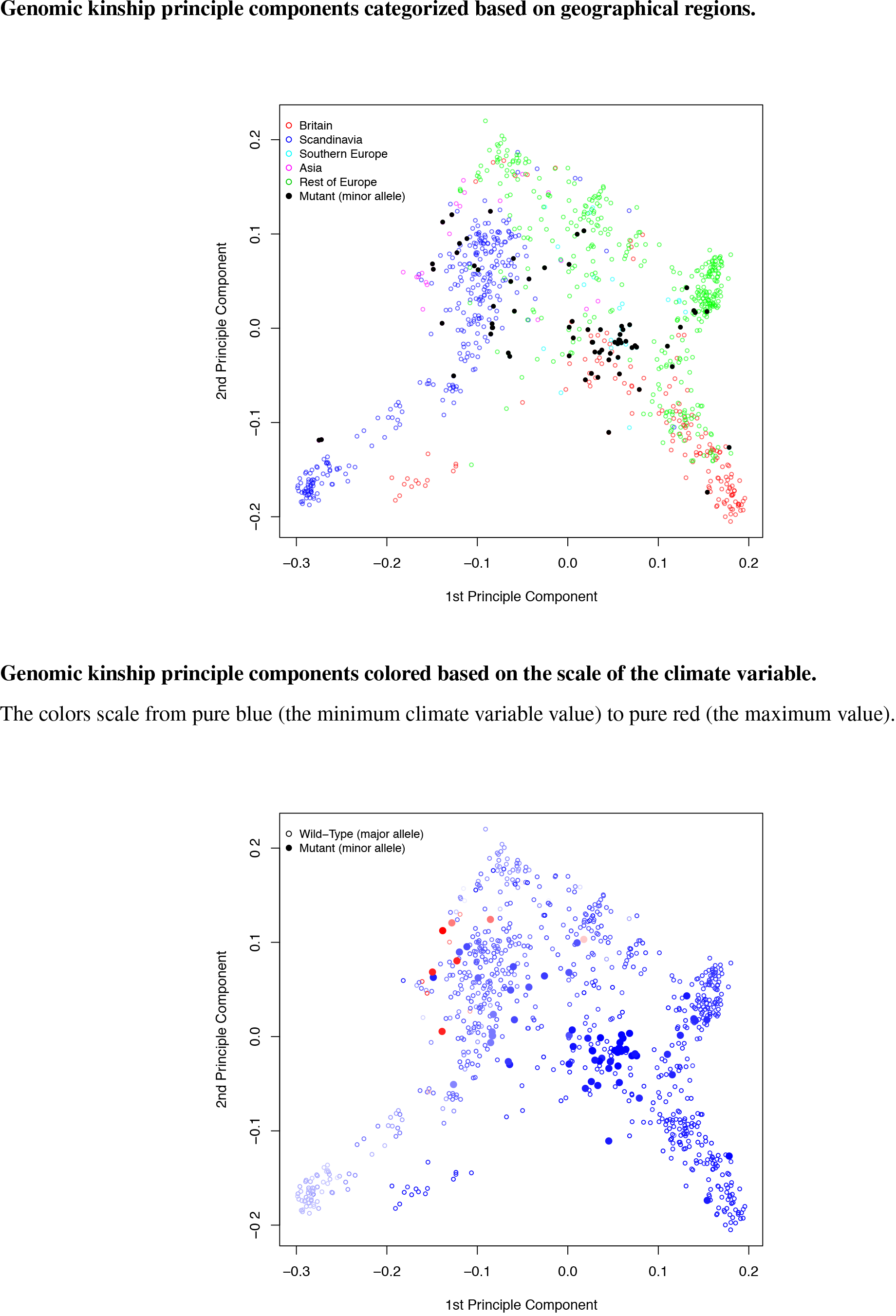
Principle components of the genomic kinship for the two alleles on chromosome 5 at 7492277 bp. Corresponding climate variable: number of consecutive cold days.

**Supplementary Figure 24.**
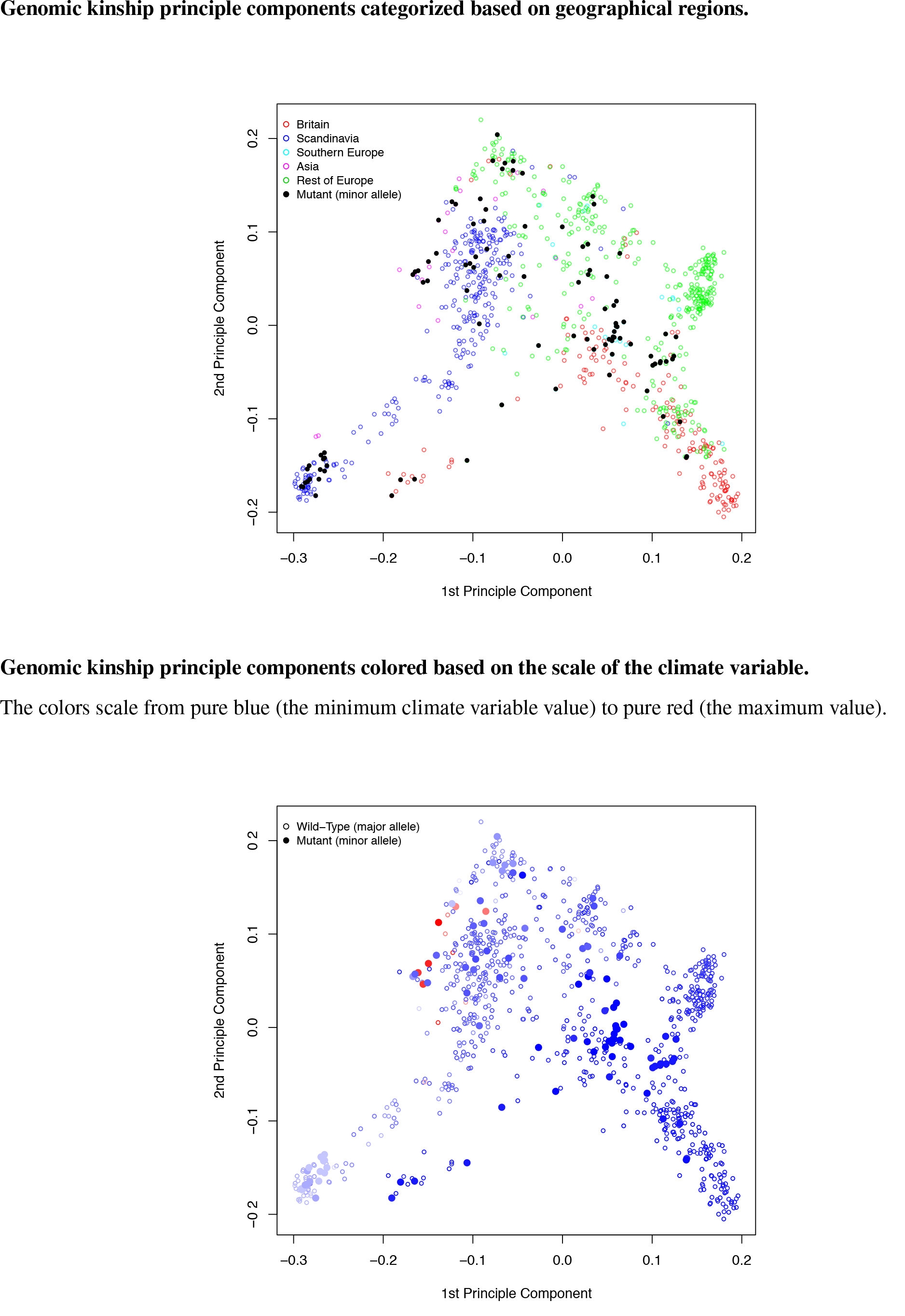
Principle components of the genomic kinship for the two alleles on chromosome 5 at 18397418 bp. Corresponding climate variable: number of consecutive cold days.

**Supplementary Figure 25.**
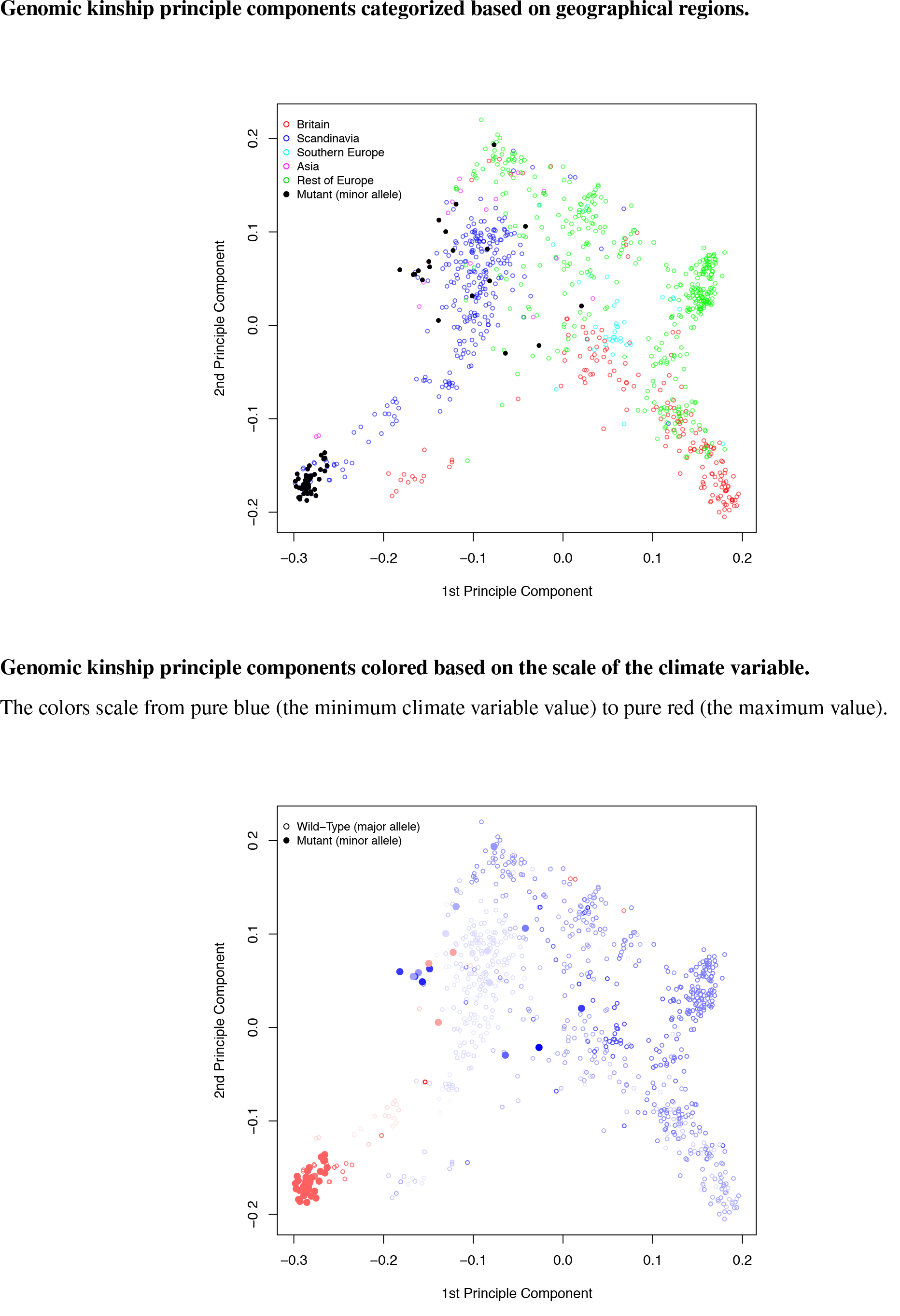
Principle components of the genomic kinship for the two alleles on chromosome 2 at 12169701 bp. Corresponding climate variable: day length in spring.

**Supplementary Figure 26.**
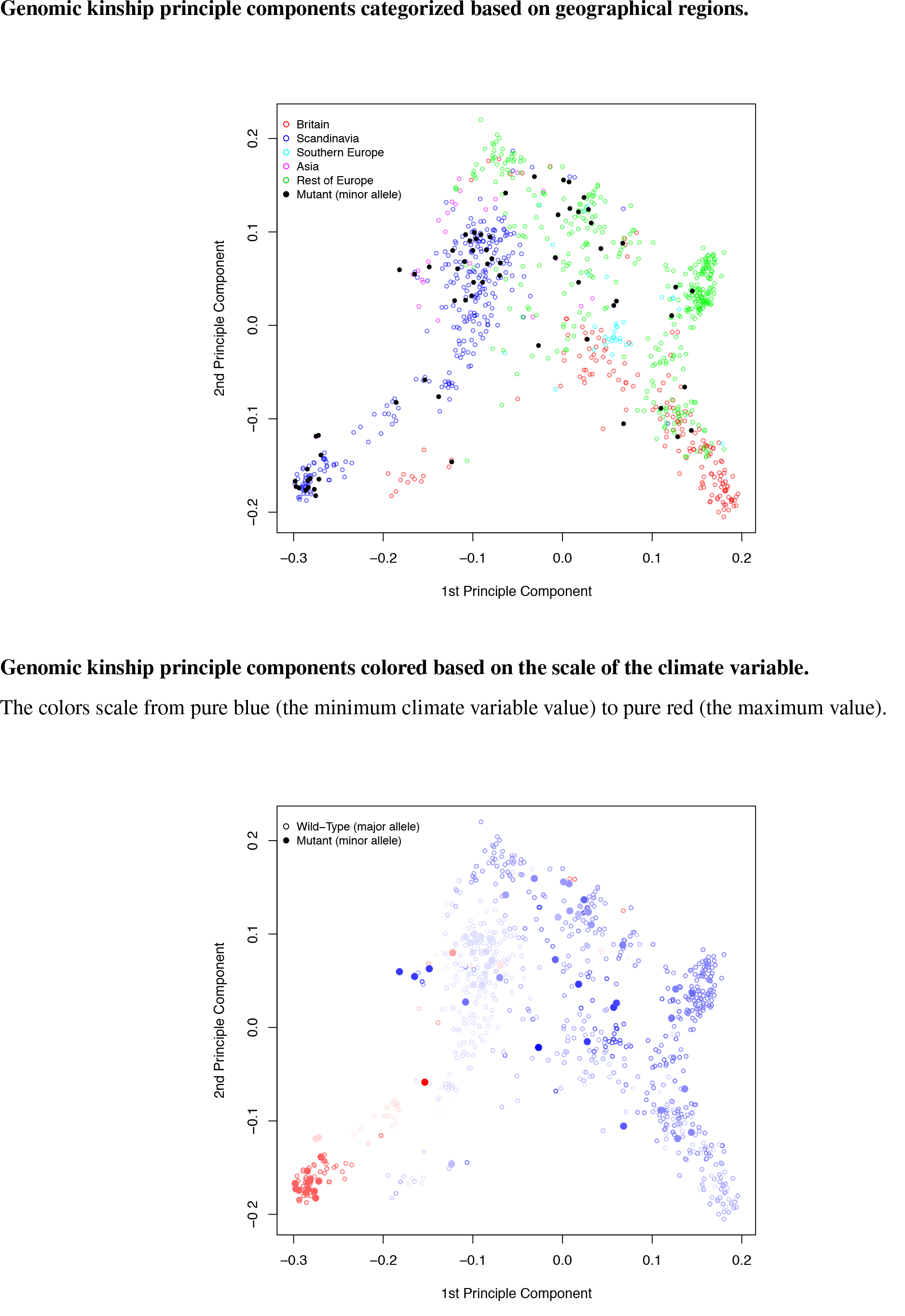
Principle components of the genomic kinship for the two alleles on chromosome 3 at 12642006 bp. Corresponding climate variable:day length in spring.

**Supplementary Figure 27.**
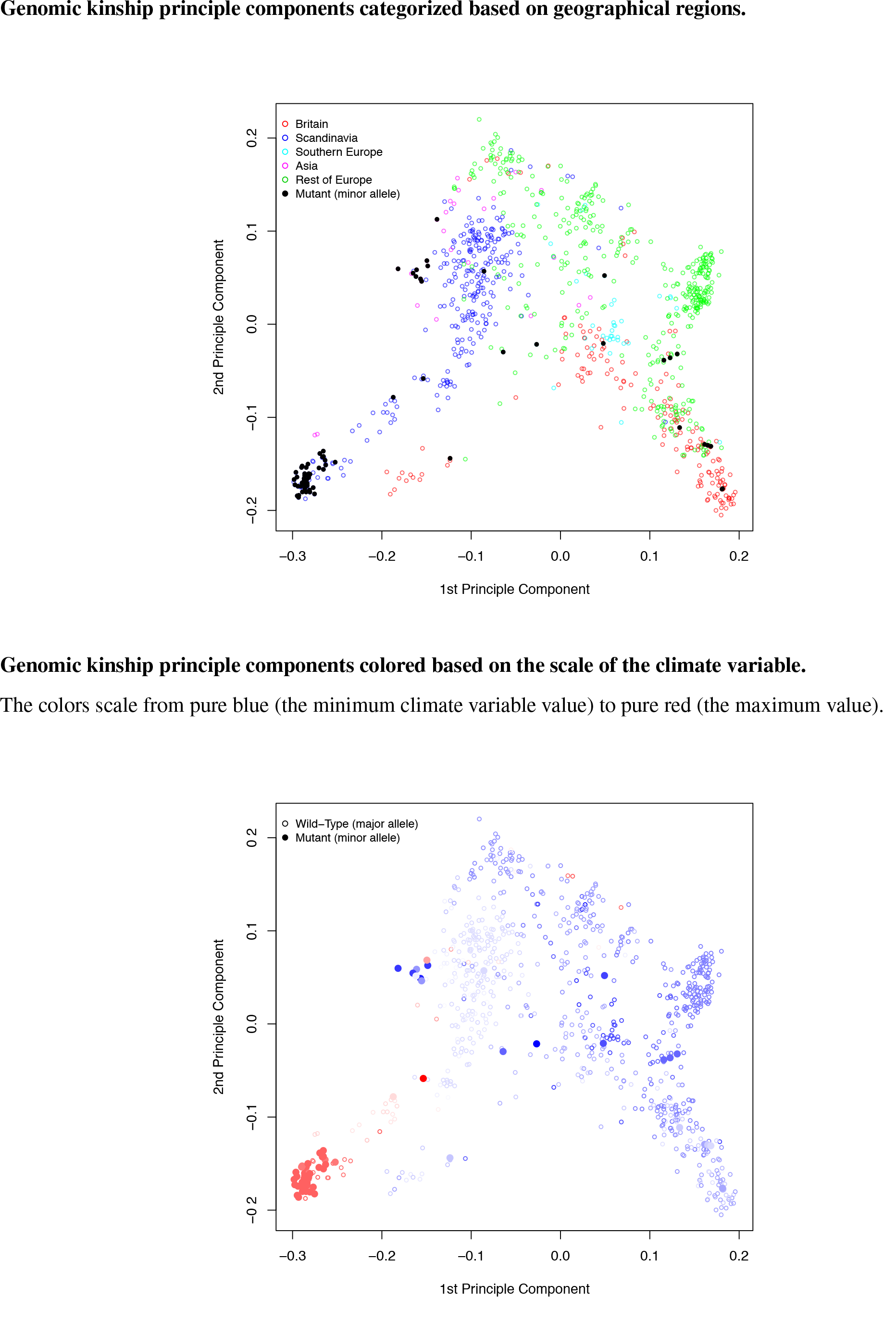
Principle components of the genomic kinship for the two alleles on chromosome 4 at 14788320 bp. Corresponding climate variable:day length in spring.

**Supplementary Figure 28.**
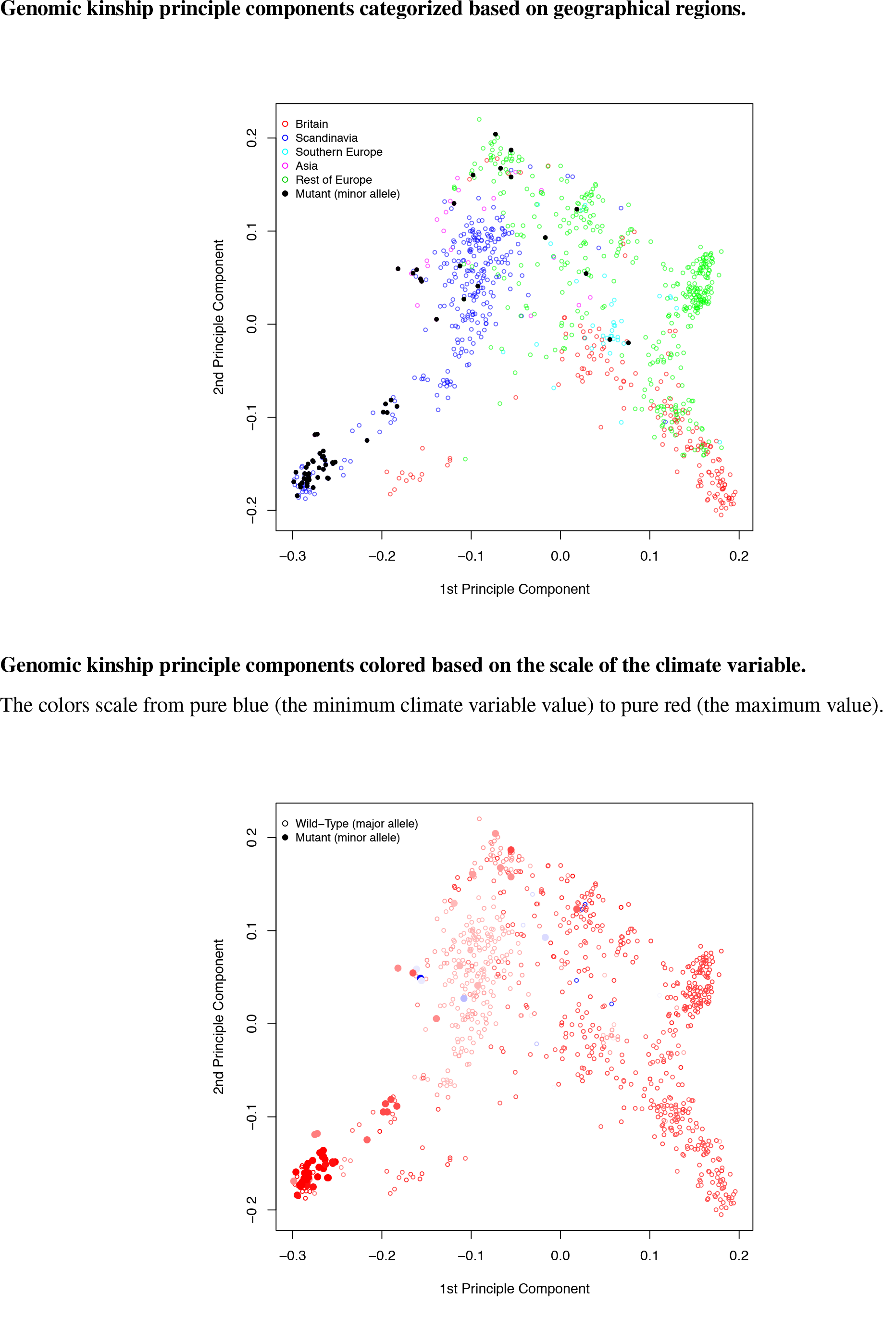
Principle components of the genomic kinship for the two alleles on chromosome 3 at 1816353 bp. Corresponding climate variable: relative humidity in spring.

**Supplementary Figure 29.**
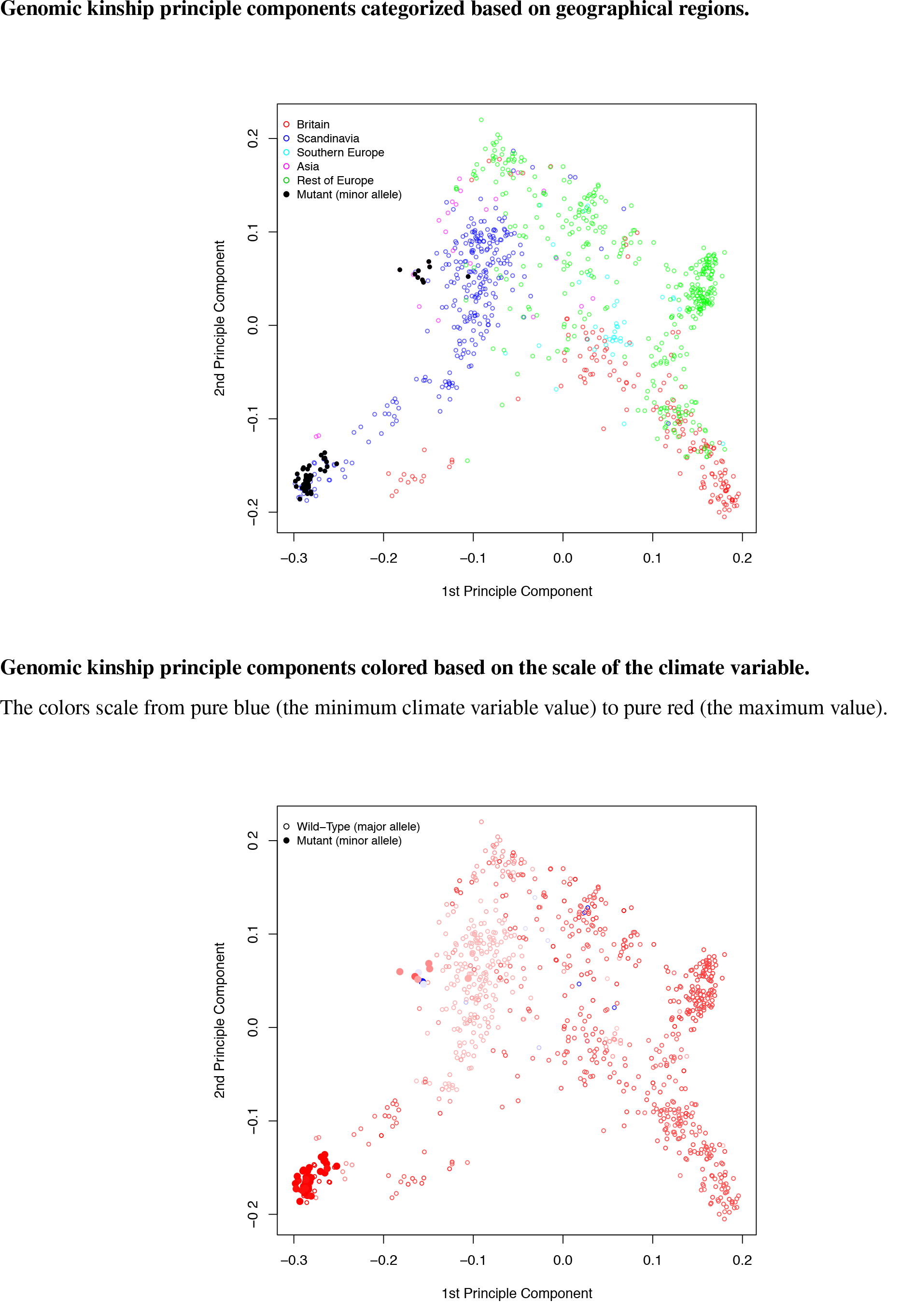
Principle components of the genomic kinship for the two alleles on chromosome 4 at 14834441 bp. Corresponding climate variable: relative humidity in spring.

**Supplementary Figure 30.**
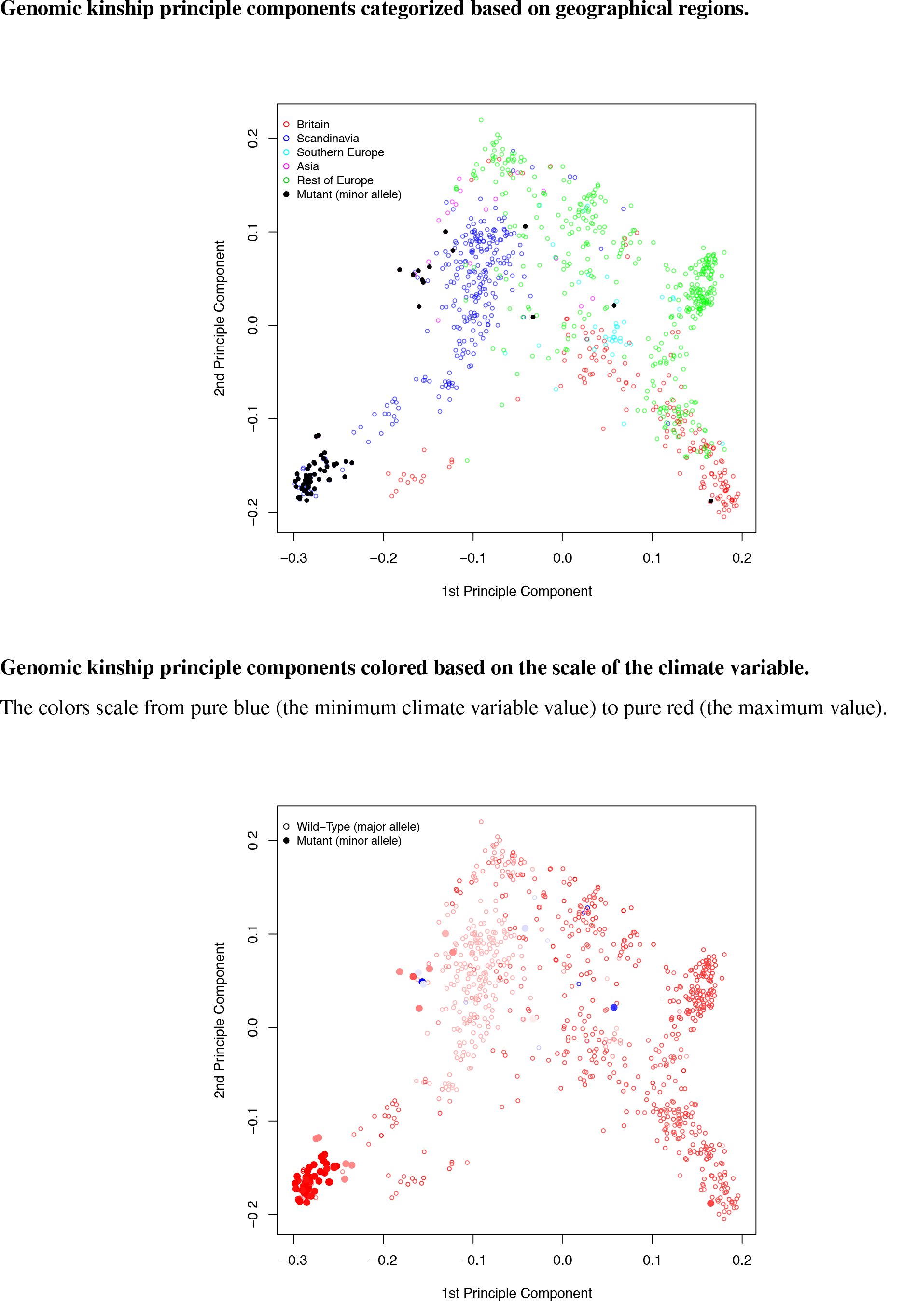
Principle components of the genomic kinship for the two alleles on chromosome 5 at 8380640 bp. Corresponding climate variable: relative humidity in spring.

**Supplementary Figure 31.**
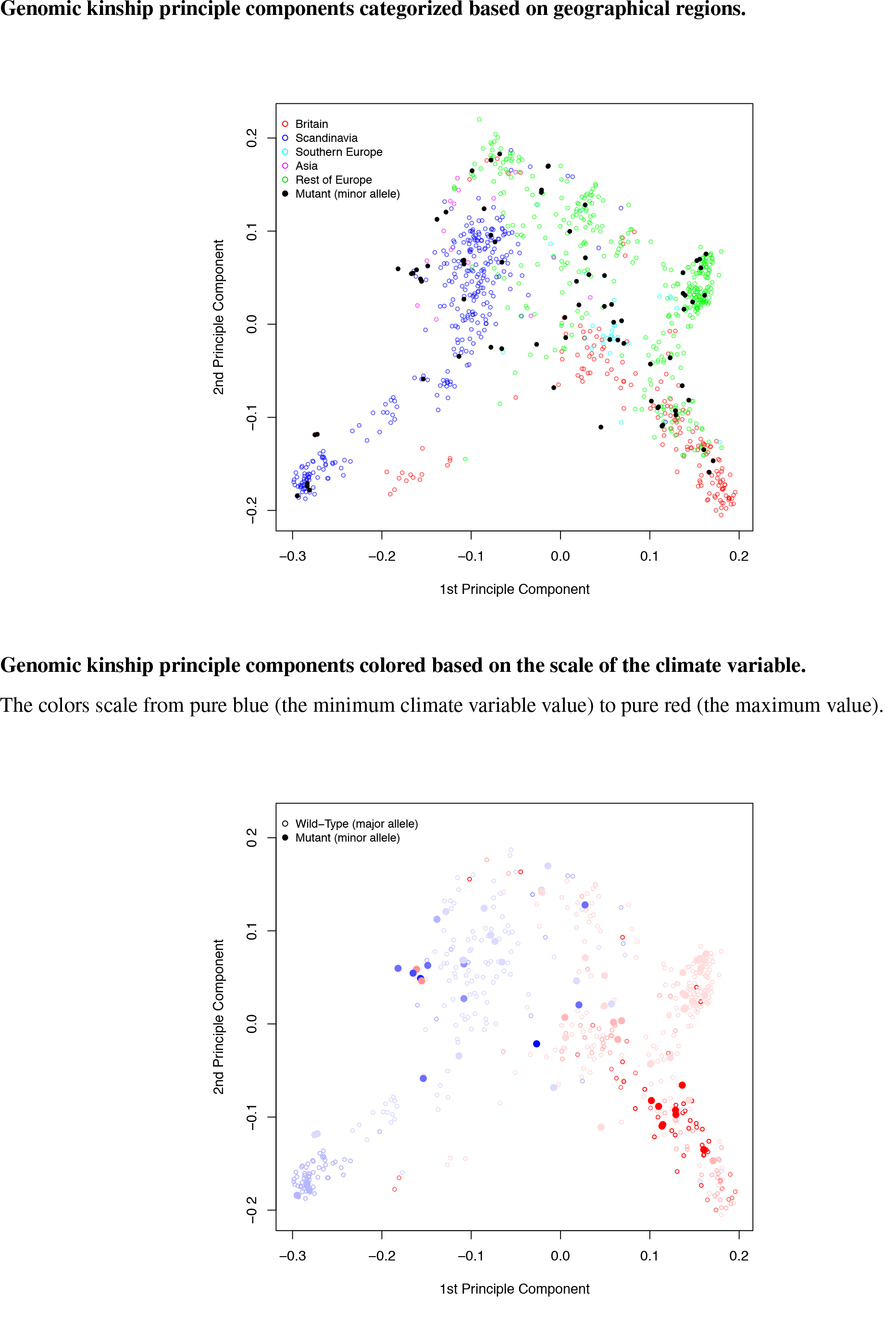
Principle components of the genomic kinship for the two alleles on chromosome 3 at 576148 bp. Corresponding climate variable: length of the growing season.

**Supplementary Figure 32.**
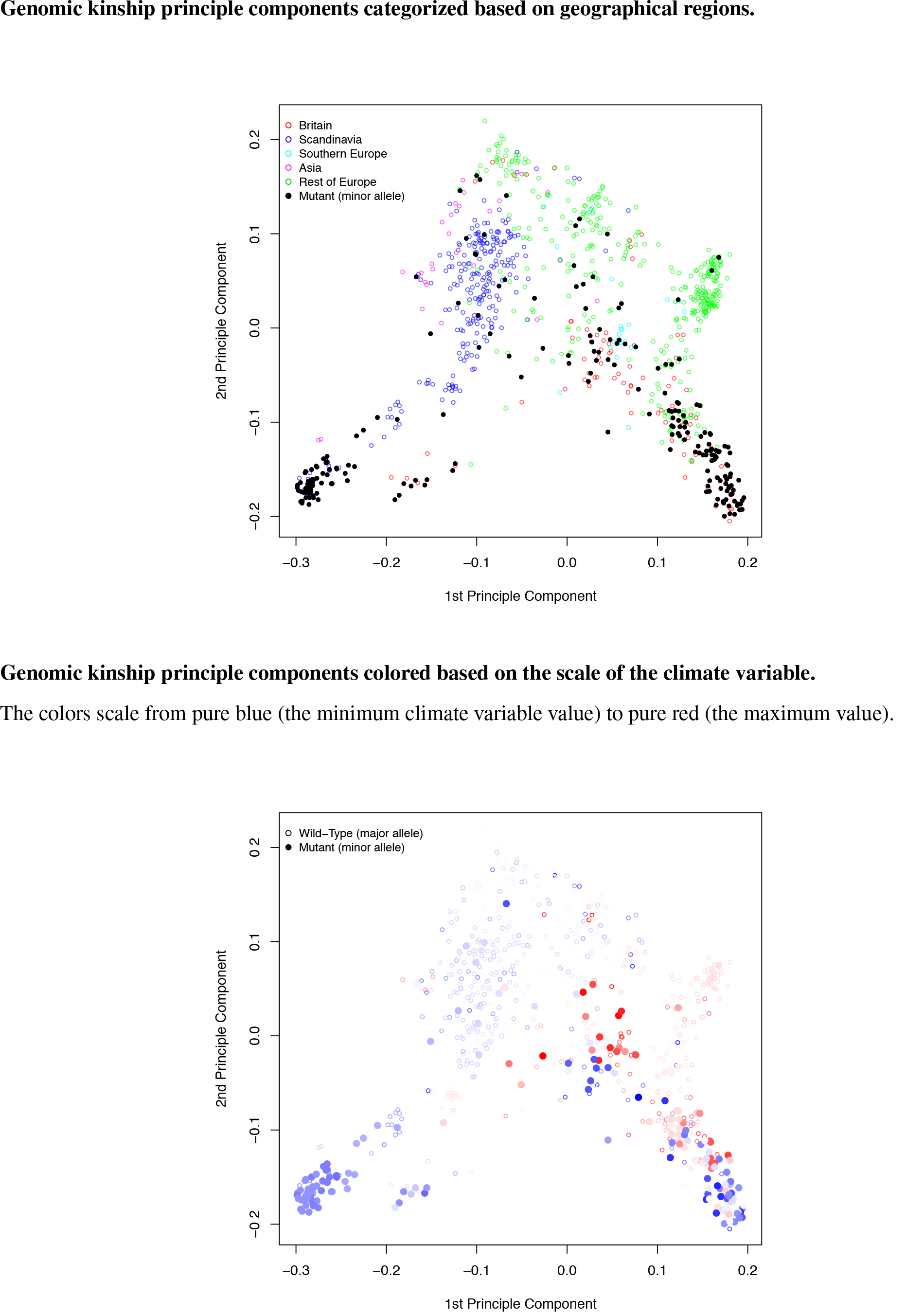
Principle components of the genomic kinship for the two alleles on chromosome 1 at 953031 bp. Corresponding climate variable: number of consecutive frost-free days.

**Supplementary Figure 33.**
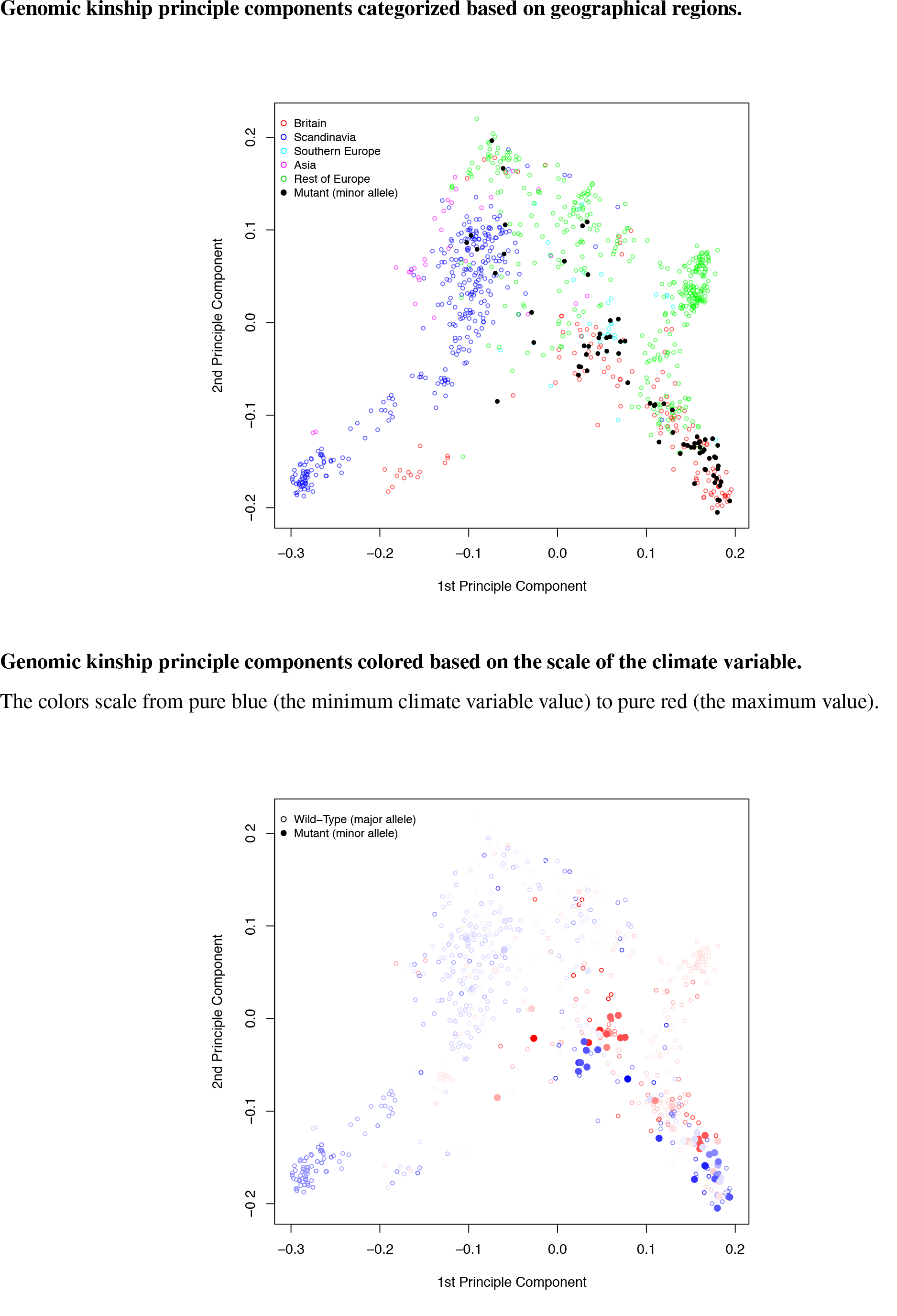
Principle components of the genomic kinship for the two alleles on chromosome 1 at 6463065 bp. Corresponding climate variable: number of consecutive frost-free days.

**Supplementary Figure 34.**
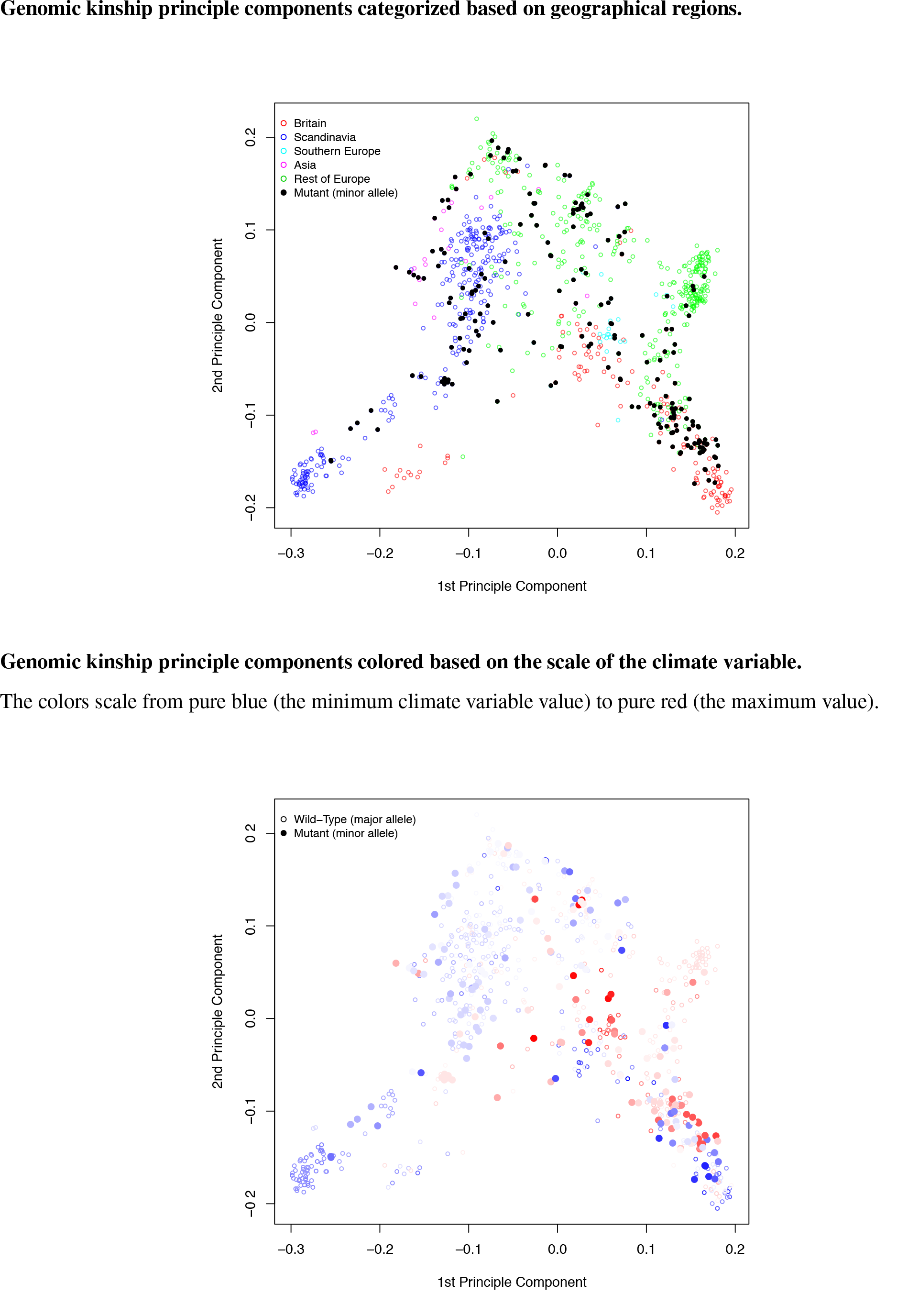
Principle components of the genomic kinship for the two alleles on chromosome 2 at 9904076 bp. Corresponding climate variable: number of consecutive frost-free days.

**Supplementary Figure 35.**
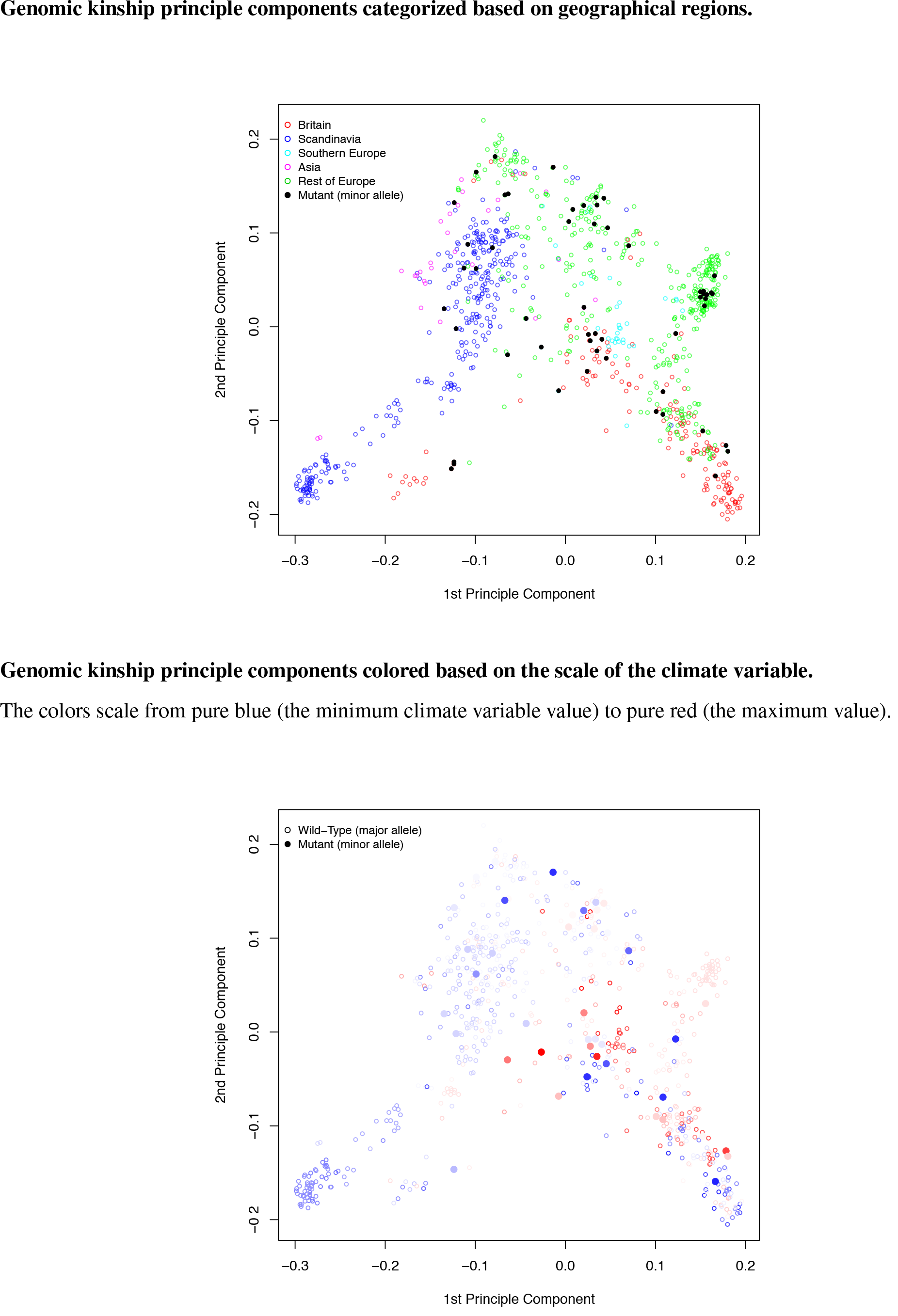
Principle components of the genomic kinship for the two alleles on chromosome 5 at 18061531 bp. Corresponding climate variable: number of consecutive frost-free days.

**Supplementary Figure 36.**
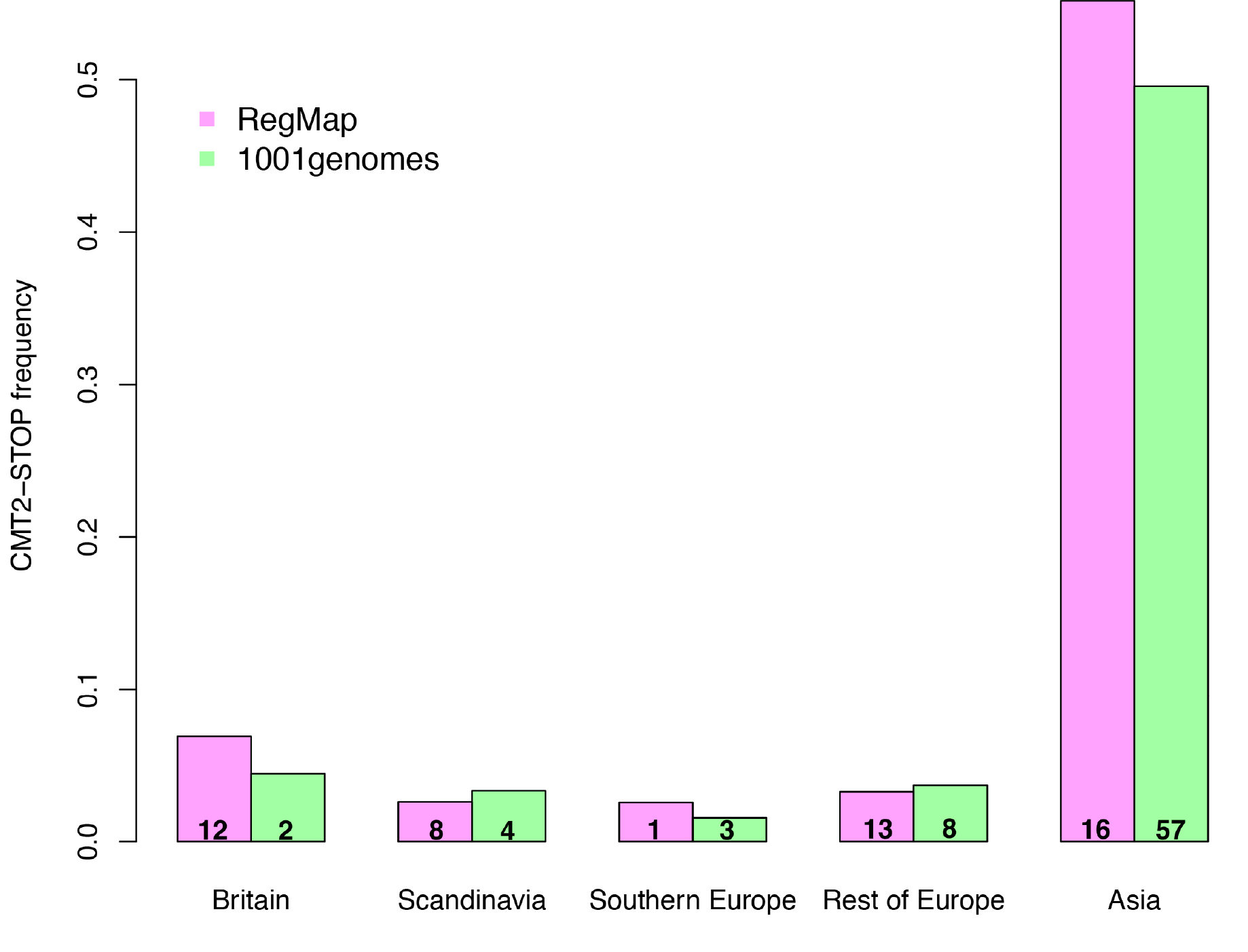
Comparison between the RegMap and 1001genomes collections in terms of the allele-frequency of CMT2_STOP_ across different geographic regions in the Euroasian *A. thaliana* population. The numbers in the bars are the number of CMT2_STOP_ alleles in this area.

**Supplementary Figure 37.**
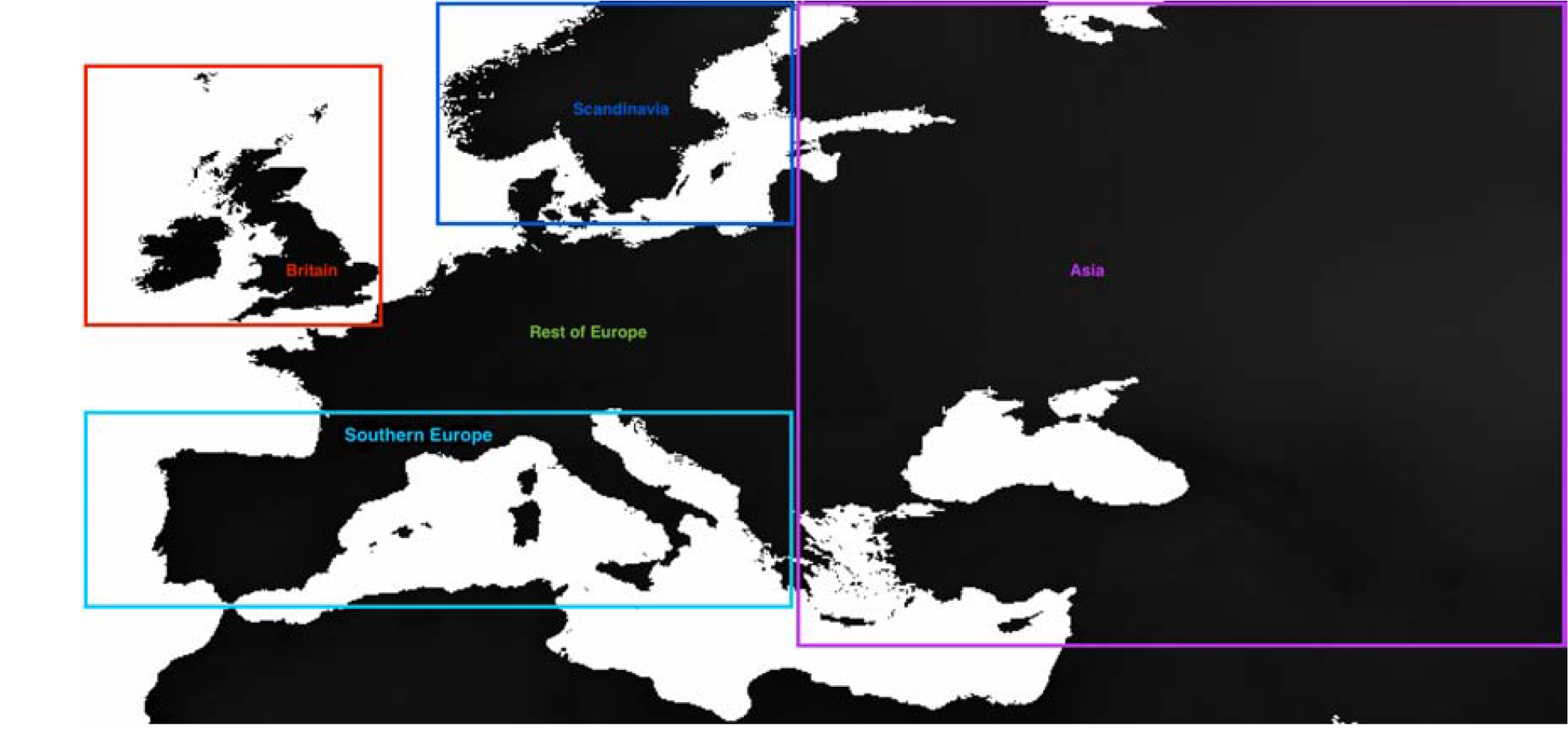
Defined geographical regions across the Euro-Asia sampling area.

**Supplementary Figure 38.**
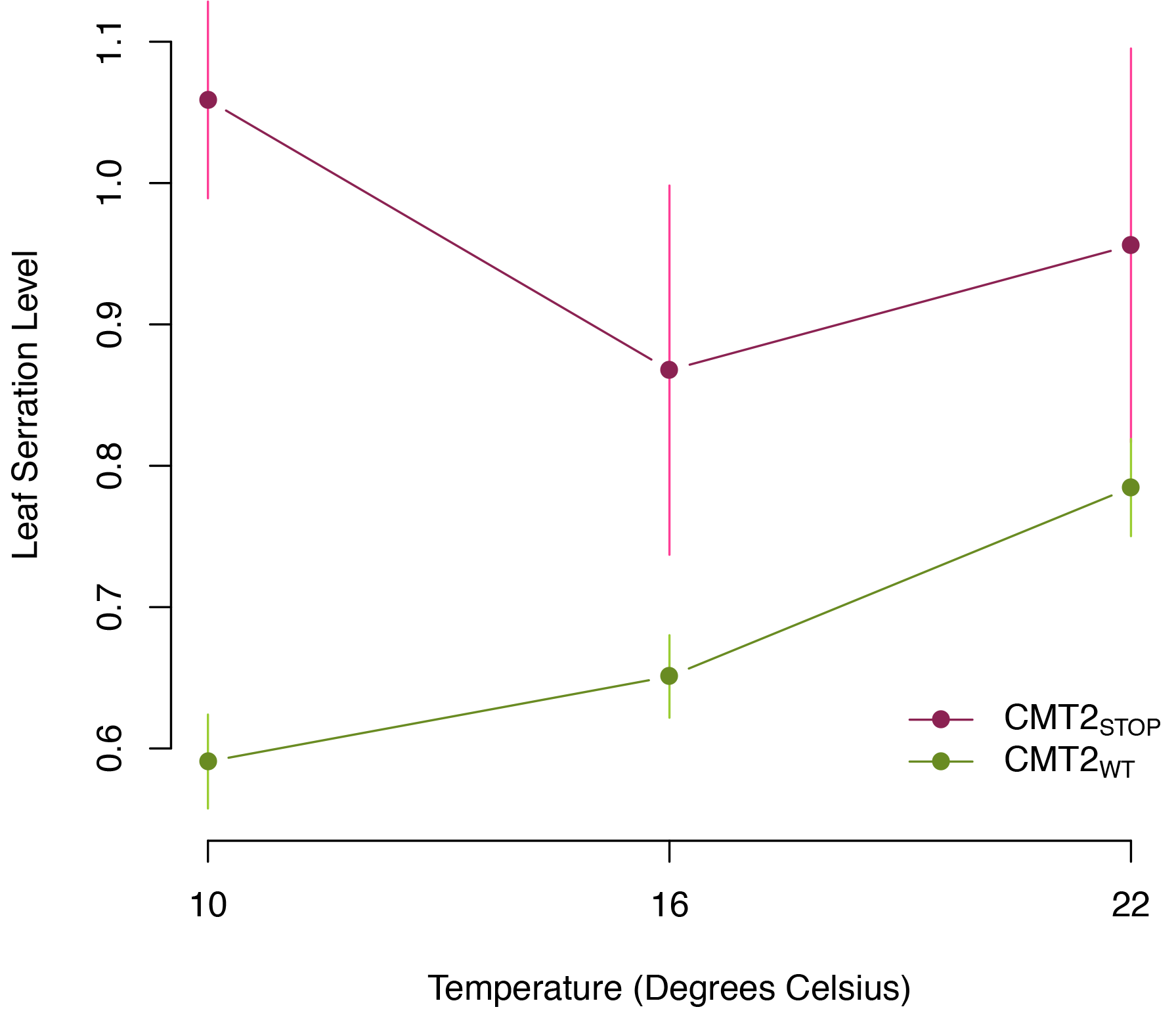
*CMT2*-by-temperature interaction effects on leaf serration. The analysis was performed using the genome-wide association data reported by Atwell et al. (2010). Each point is the mean leaf serration level of a combination of CMT2 genotype and temperature. The vertical bars represent standard errors of the mean estimates.

**Supplementary Figure 39.**
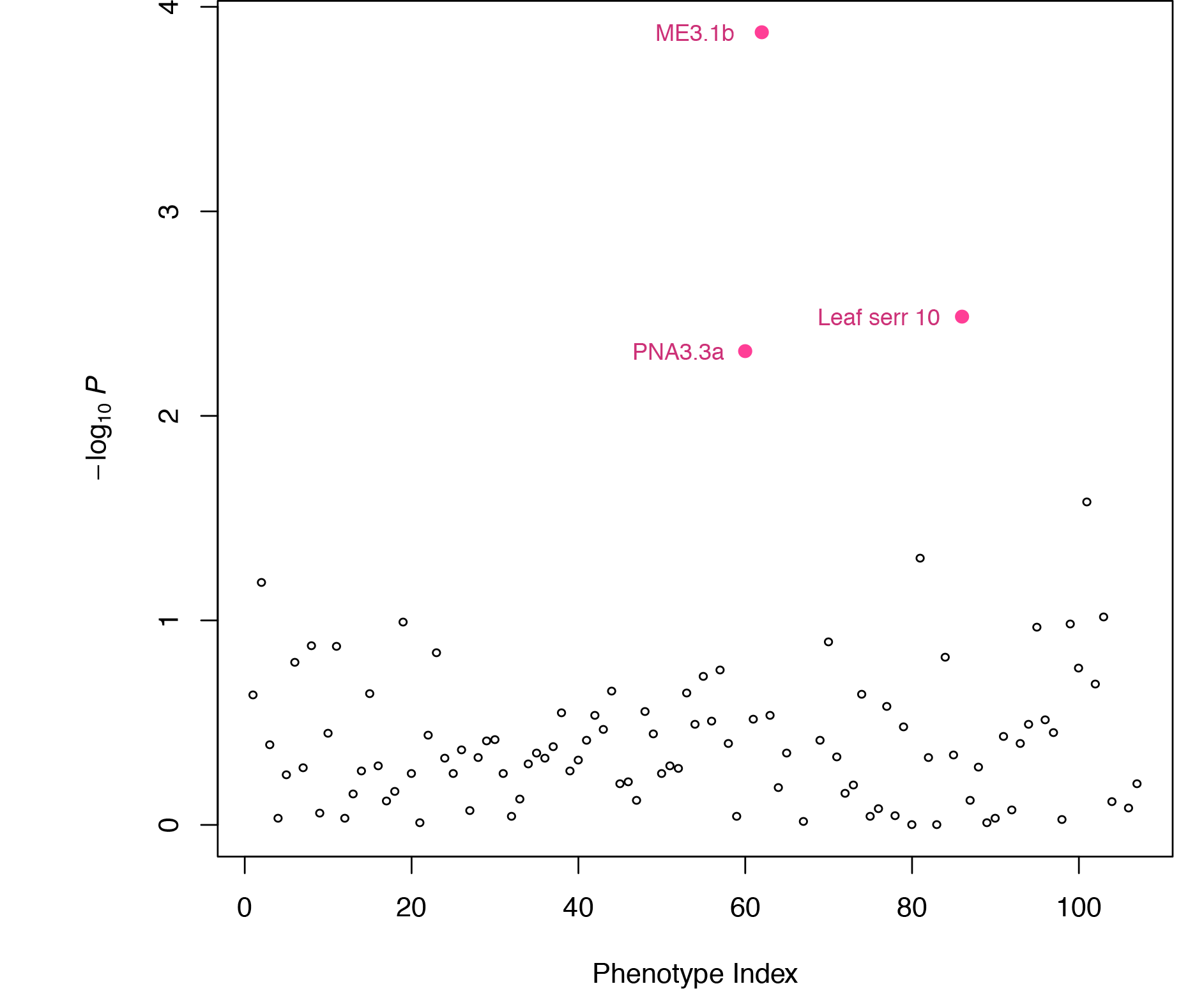
Associations between the *CMT2*_*STOP*_ genotype and the 107 scored phenotypes in Atwell et al. (2010) The most significant three associations are labeled in pink,with a false discovery rate of 0.17. The definition of each labeled phenotype should be referred to the Supplementary Tables in Atwell et al. (2010).

**Supplementary Figure 40.**
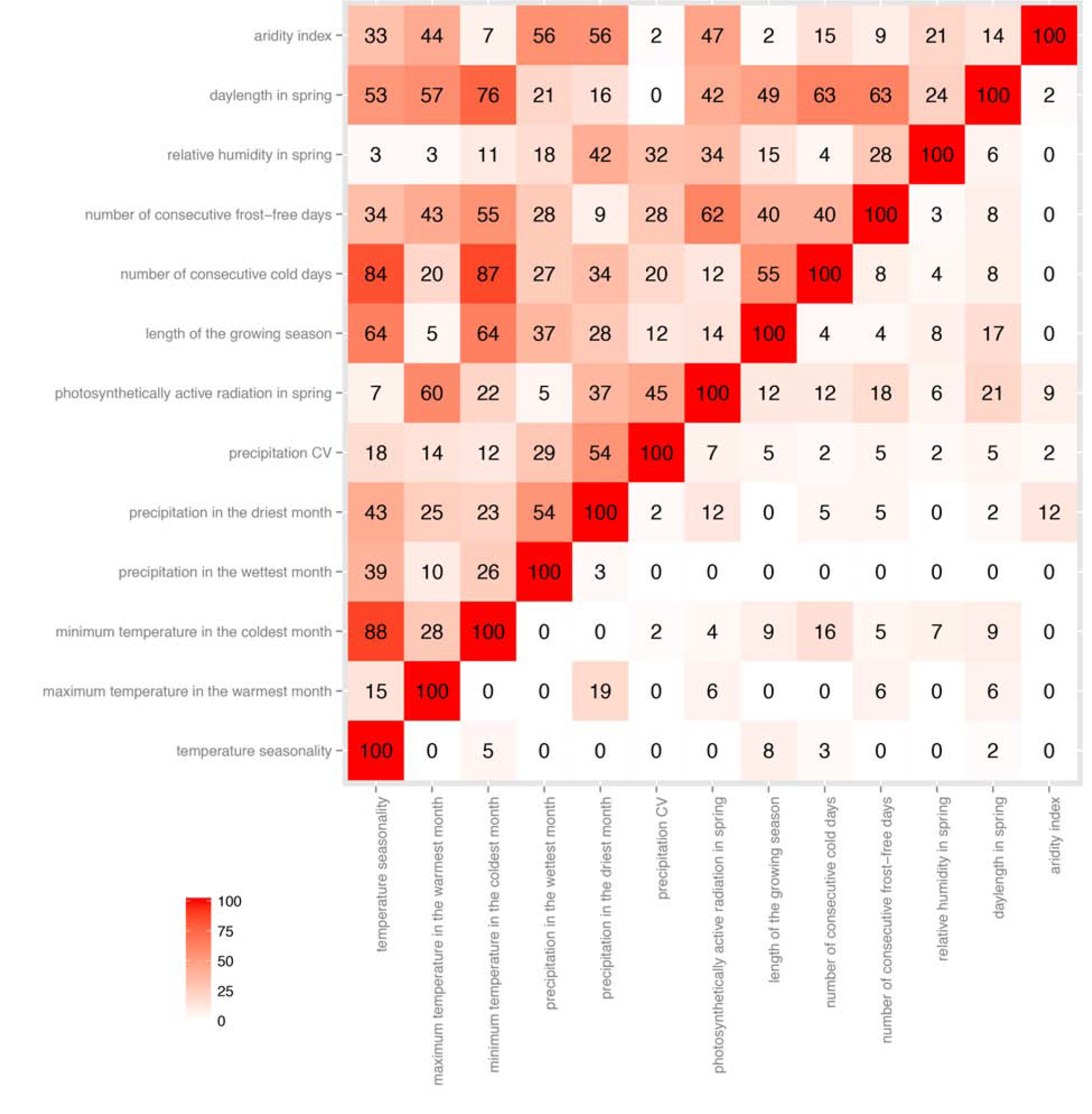
Comparison between the correlations among the climate variables (upper triangle) and the overlap in variance-heterogeneity GWA profiles (lower triangle). Numbers shown in the figure are percentages. Pearson’s correlation coefficients were calculated for each pair of the climate variables. Overlaps in GWA profiles were calculated as the proportion of shared SNPs above the threshold of 1.0 × 10^−4^.

**Supplementary Figure 41.**
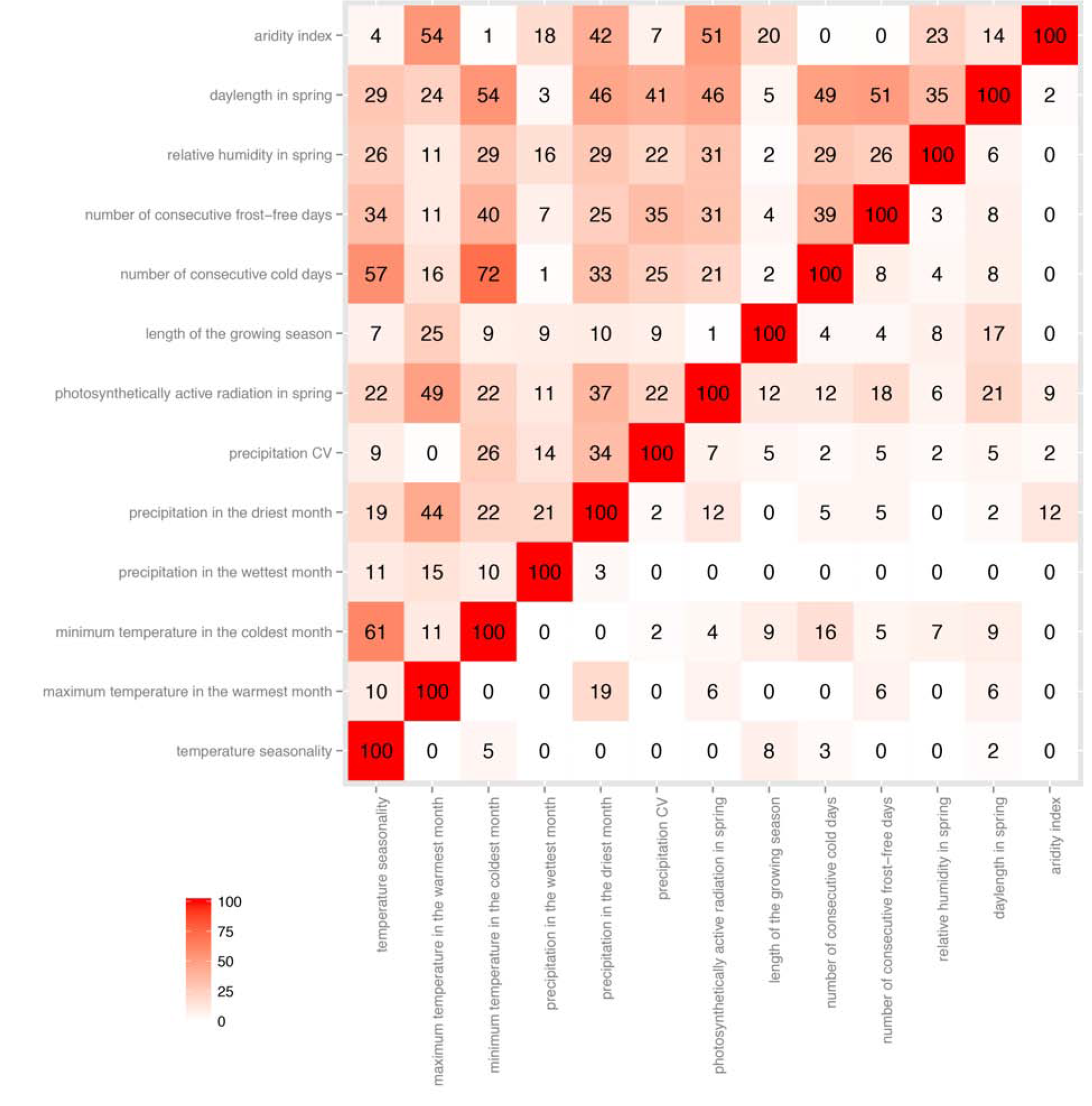
Comparison between the correlations among the residual climate variables after genomic
kinship correction (upper triangle) and the overlap in variance-heterogeneity GWA profiles (lower triangle). Numbers shown in the figure are percentages. Pearson’s correlation coefficients were calculated for each pair of the climate variables. Overlaps in GWA profiles were calculated as the proportion of shared SNPs above the threshold of 1.0 × 10^−4^.

**Supplementary Figure 42.**
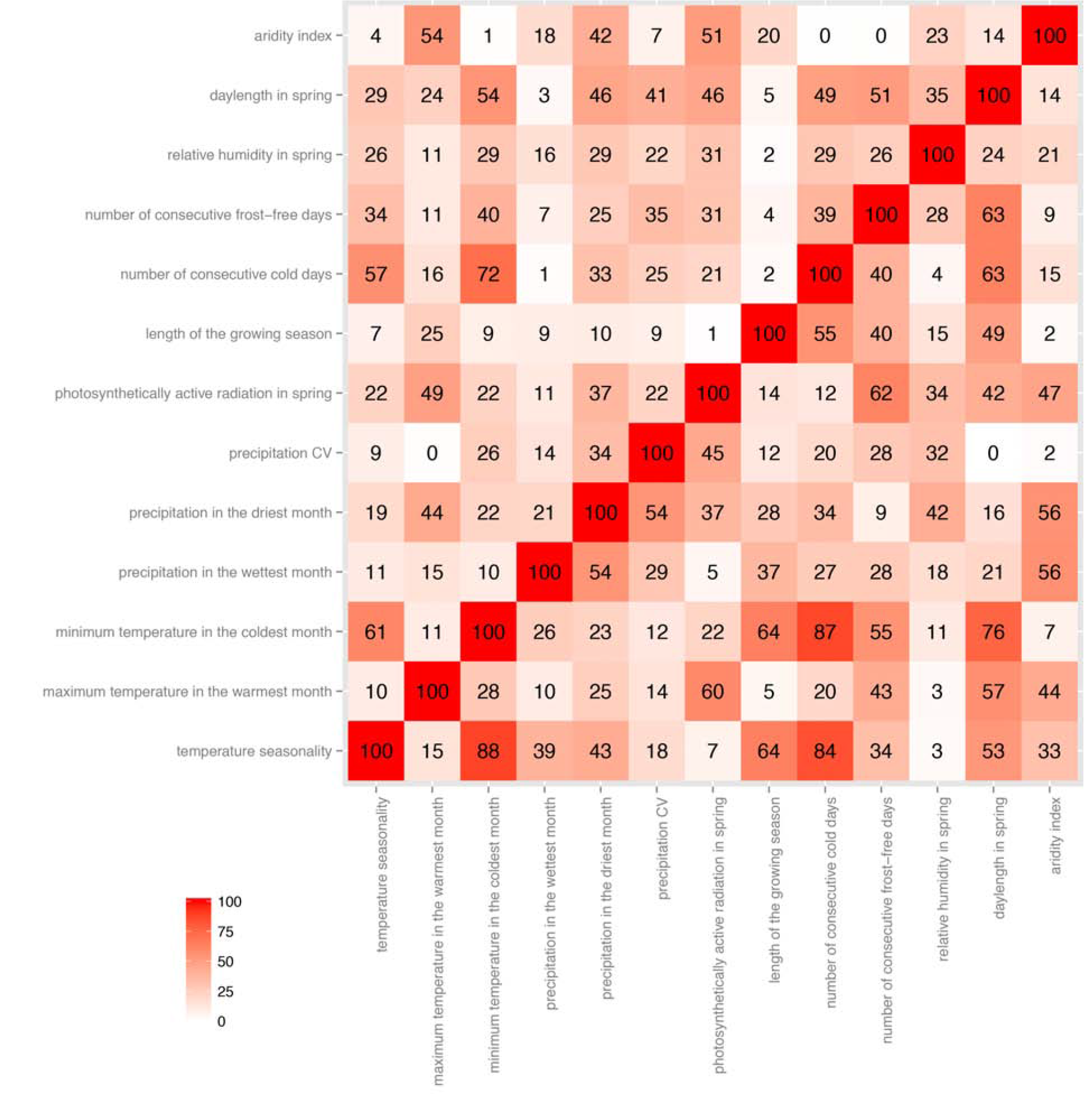
Comparison between the correlations among the residual climate variables after genomic kinship correction (upper triangle) and the correlations among the original climate variables (lower triangle). Numbers shown in the figure are percentages. Pearson’s correlation coefficients were calculated for each pair of the climate variables.

**Supplementary Figure 43.**
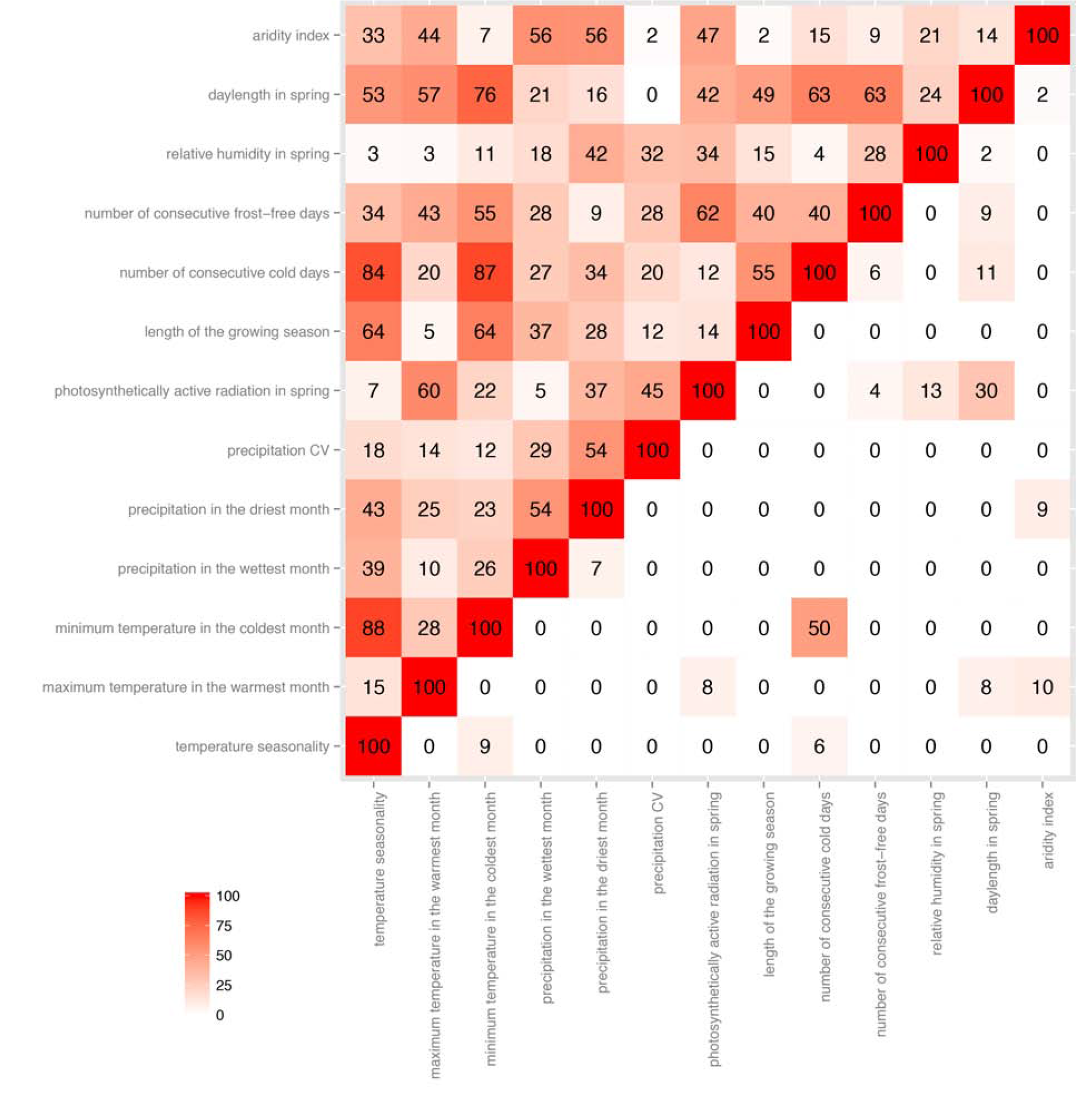
Comparison between the correlations among the climate variables (upper triangle) and the overlap in ordinary GWA profiles (lower triangle). Numbers shown in the figure are percentages. Pearson’s correlation coefficients were calculated for each pair of the climate variables. Overlaps in GWA profiles were calculated as the proportion of shared SNPs above the threshold of 1.0 ×10^−4^.

**Supplementary Figure 44.**
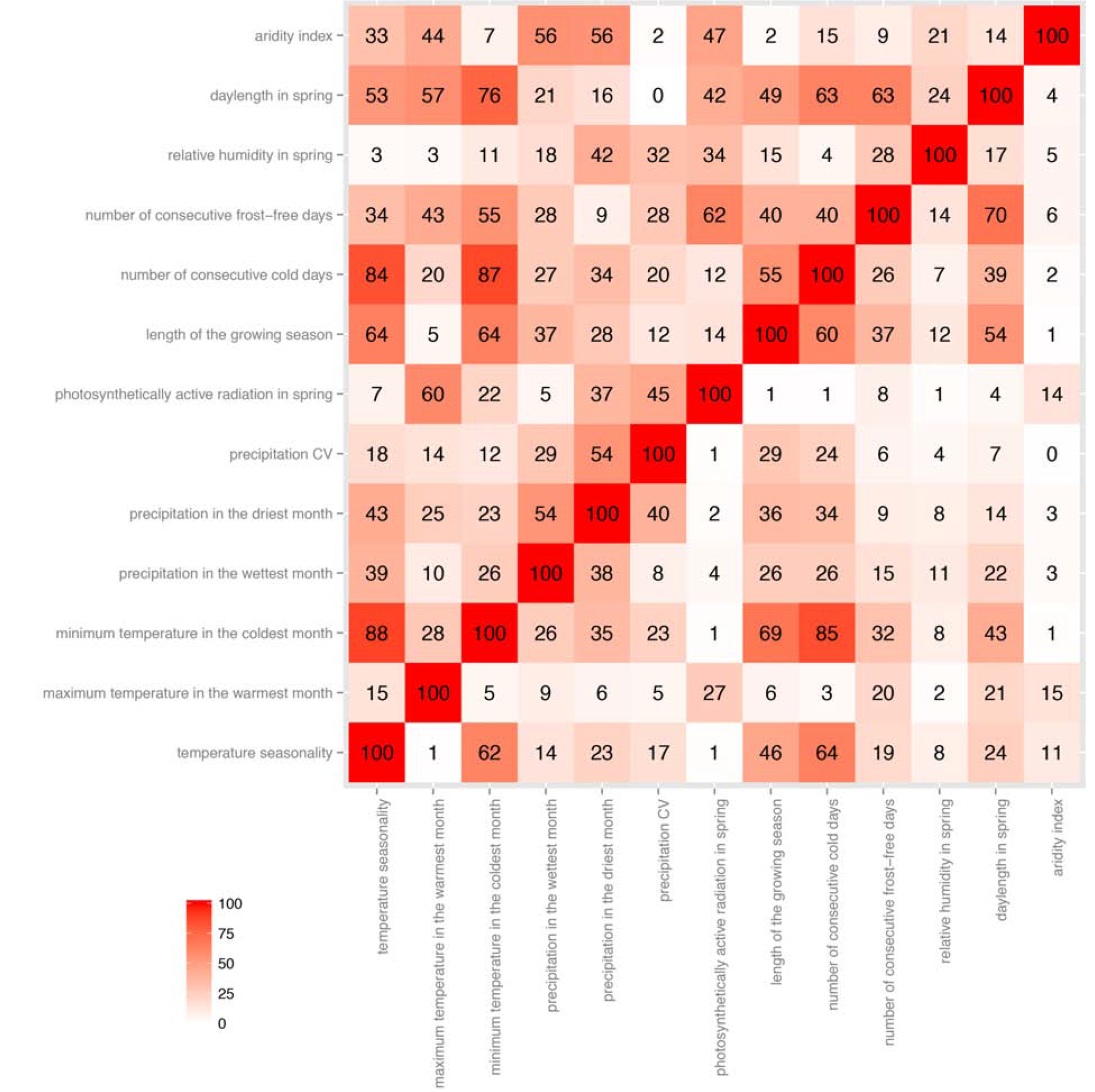
Comparison between the correlations among the climate variables (upper triangle) and the overlap in simple GWA profiles without correction for population structure (lower triangle). Numbers shown in the figure are percentages. Pearson’s correlation coefficients were calculated for each pair of the climate variables. Overlaps in GWA profiles were calculated as the proportion of shared SNPs above the threshold of 1.0 ×10^−4^.

**Supplementary Figure 45.**
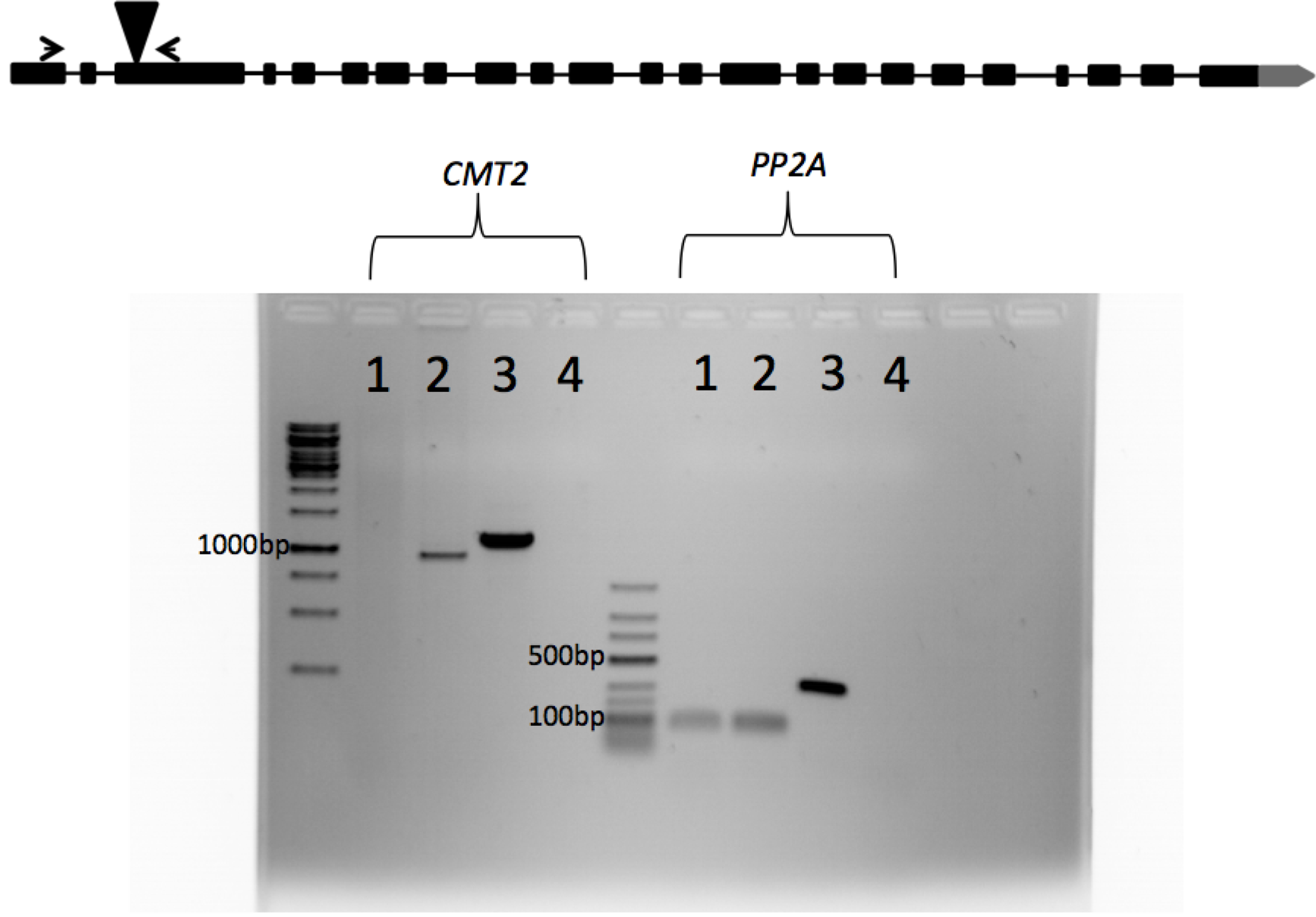
Gene-model of CMT2 and T-DNA insertion confirmation. Boxes indicate exons, lines represent introns. The triangle shows the T-DNA insertion site. Arrow heads indicate the location of primers that were used to assay *CMT2* transcripts. *CMT2*: PCR reaction with *CMT2*-specific primers, PP2A: PCR reaction with PP2A-specific primers. Lanes 1:*cmt2-5* cDNA, lanes 2: Col cDNA, lanes 3:Col genomic DNA, lanes 4: no template controls. *CMT2* cDNA and genomic DNA are predicted to give 940bp and 1159bp bands, respectively. PP2A cDNA and genomic DNA are predicted to give 84 bp and 210 bp bands, respectively.

**Supplementary Figure 46.**
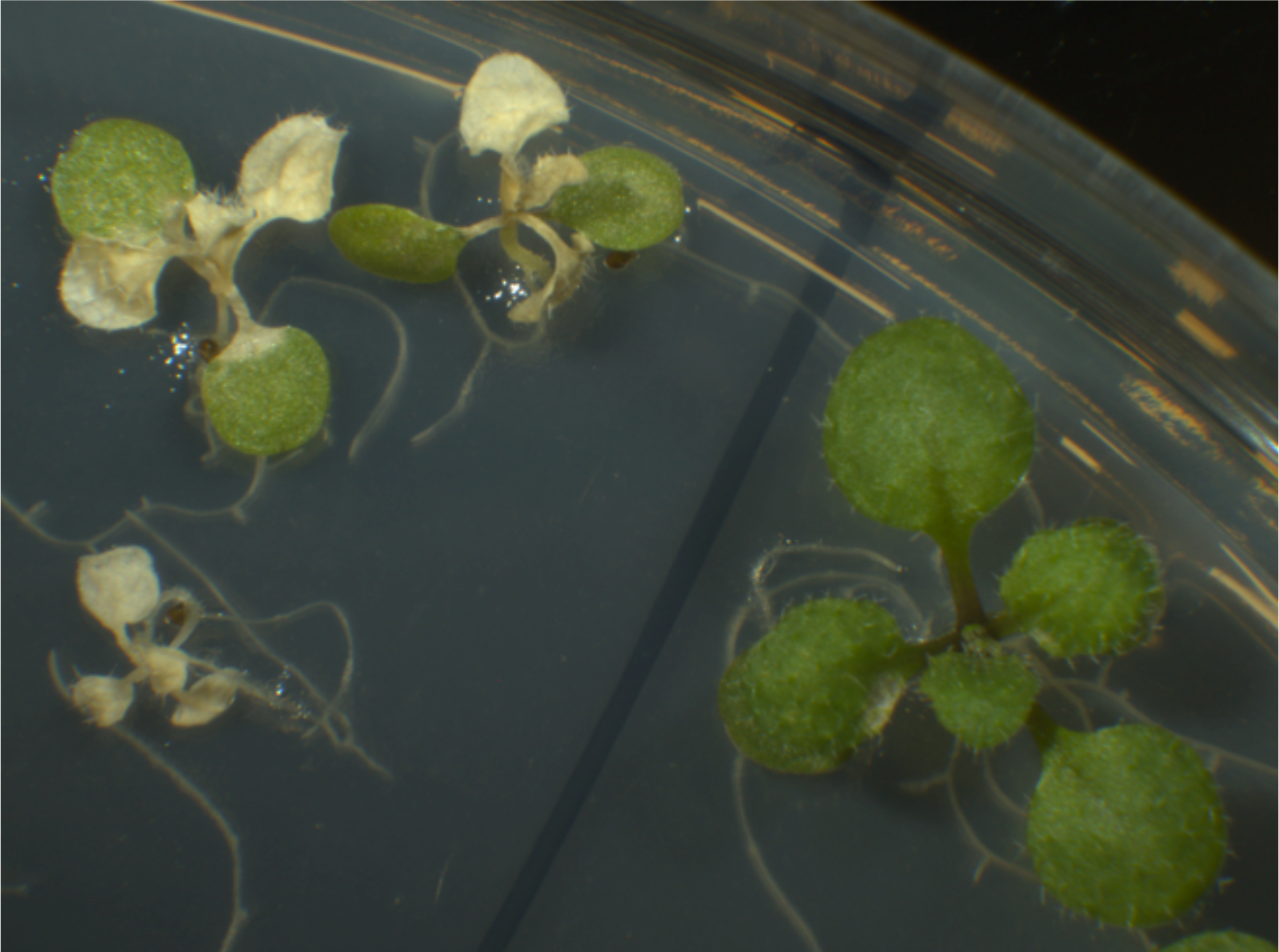
Prolonged heat stress is often lethal. 10-days-old seedlings were heat-stressed at 37.5°C for 24 h based on a published protocol (Ito et al. 2011). Plants were counted as non-viable if shoot apices were completely bleached. Note that the lamina of cotyledons often remains green for a longer time but no recovery was observed if apices were
bleached.

**Table S1.**
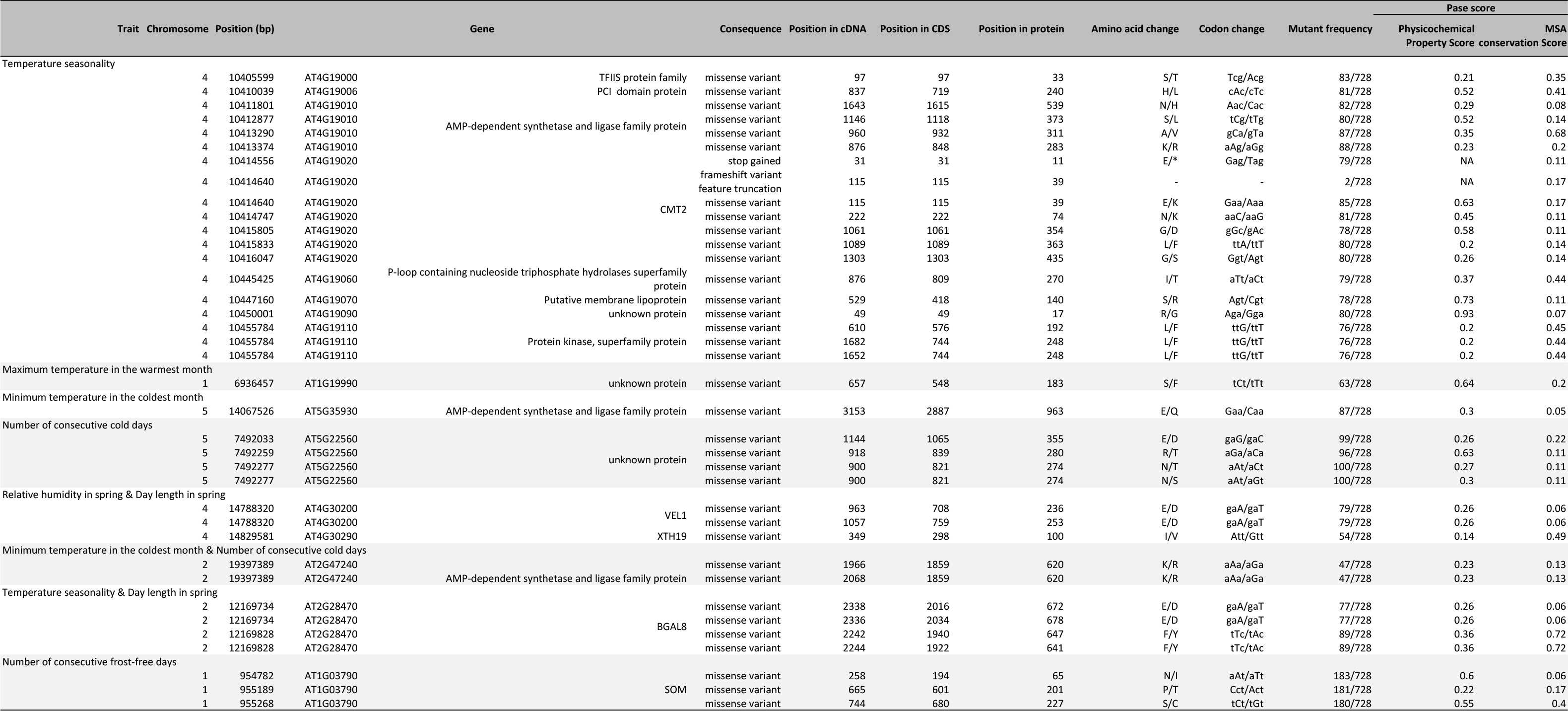
Detailed information about the missense mutations significantly associated with climate adaptability of Arabidopsis thaliana.

**Table S2.** Loci significantly associated with climate adaptability of Arabidopsis thaliana but without non-synonymous mutations in high LD detected. P-values were obtained from linear regression of squared z-scores. GC P-values were the P-values after genomic control. Gamma P-values were obtained by fitting generalized linear models with Gamma response. Pleiotropic loci are marked with stars. bp = base pair; MAF = minor allele frequency.

| *Trait* | Leading SNP |  |  |  |  |  |
| --- | --- | --- | --- | --- | --- | --- |
|  | *Chromosome* | *Position (bp)* | *MAF* | *P-value* | *GC P-value* | *Gamma P-value* |
| Relative humidity in spring | 3 | 1816353 | 0.072 | 1.8E-10 | 1.2E-08 | 1.0E-14 |
|  | 5 | 8380640 | 0.073 | 1.4E-09 | 6.3E-08 | 9.2E-17 |
| Minimum temperature in the coldest month & Number of consecutive cold days* | 2 | 18620697 | 0.083 | 7.2E-11 | 3.5E-08 | 1.2E-11 |
|  |  |  |  | *7.2E-10 | *1.7E-07 | *1.4E-11 |
|  | 5 | 18397418 | 0.107 | 3.8E-10 | 1.2E-07 | 1.5E-13 |
|  |  |  |  | *8.8E-10 | *1.9E-07 | *3.2E-13 |
| Length of the growing season | 3 | 576148 | 0.084 | 1.1E-09 | 1.4E-07 | 8.9E-09 |
| Number of consecutive frost-free days | 1 | 953031 | 0.237 | 1.0E-08 | 8.0E-08 | 2.4E-21 |
|  | 1 | 6463065 | 0.081 | 2.7E-08 | 1.9E-07 | 2.9E-09 |
|  | 2 | 9904076 | 0.215 | 2.3E-09 | 2.2E-08 | 1.8E-10 |

**Table S3.** Experimental data of the heat-stress treatment on Col-0 and CMT2 individuals.

| experiment | accession | total | alive | dead |
| --- | --- | --- | --- | --- |
| 1 | cmt2-5 | 28 | 17 | 11 |
| 1 | cmt2-5 | 25 | 21 | 4 |
| 1 | Col-0 | 25 | 12 | 13 |
| 1 | Col-0 | 23 | 5 | 18 |
| 2 | Col-0 | 28 | 8 | 20 |
| 2 | Col-0 | 30 | 4 | 26 |
| 2 | cmt2-5 | 26 | 10 | 16 |
| 2 | Col-0 | 25 | 4 | 21 |
| 2 | cmt2-5 | 19 | 11 | 8 |
| 2 | Col-0 | 29 | 11 | 18 |
| 3 | cmt2-5 | 25 | 20 | 5 |
| 3 | Col-0 | 29 | 21 | 8 |
| 3 | cmt2-5 | 26 | 16 | 10 |
| 3 | Col-0 | 29 | 12 | 17 |
| 3 | Col-0 | 29 | 13 | 16 |
| 3 | Col-0 | 29 | 14 | 15 |

**Table S4.** Experimental data of root growth (mm) of Col-0 and CMT2 knockouts, with and without 6 h heat stress.

| Col |  | cmt2 |  |
| --- | --- | --- | --- |
| control | 6h heat | control | 6h heat |
| 11.839 | 13.48 | 17.375 | 20.214 |
| 10.911 | 0.364 | 28.15 | 22.313 |
| 20.818 | 22.539 | 26.546 | 23.846 |
| 8.357 | 0 | 20.006 | 22.25 |
| 13.656 | 1.214 | 19.581 | 18.644 |
| 23.621 | 21.834 | 8.576 | 9.141 |
| 16.425 | 18.095 | 23.061 | 18.037 |
| 26.866 | 7.785 | 26.788 | 3.843 |
| 7.801 | 14.861 | 23.047 | 9.019 |
| 23.397 | 8.013 | 33.445 | 24.753 |
| 11.484 | 0.243 | 27.716 | 6.607 |
| 8.684 | 10.881 | 21.39 | 33.157 |
| 18.94 | 10.902 | 23.379 | 19.084 |
| 21.055 | 1.093 | 14.33 | 5.901 |
| 21.355 | 0 | 19.115 | 22.115 |
| 26.758 | 16.586 | 29.795 | 1.75 |
| 25.958 | 6.19 | 18.046 | 1.996 |
| 23.438 | 0.777 |  | 6.376 |
| 23.419 | 12.242 |  | 17.121 |
| 22.792 | 0.384 |  | 15.948 |
| 21.526 | 1.341 |  | 7.113 |
| 26.55 | 0.738 |  | 19.502 |
| 16.929 | 0.619 |  | 6.085 |
| 29.122 | 24.244 |  | 14.354 |
| 27.851 | 25.941 |  | 1.573 |
| 24.558 | 10.512 |  | 16.381 |
| 24.275 | 0.243 |  | 5.892 |
| 30.535 | 23.916 |  |  |
| 30.152 |  |  |  |
| 18.197 |  |  |  |

